# Phylogenetic Biodiversity Metrics Should Account for Both Accumulation and Attrition of Evolutionary Heritage

**DOI:** 10.1101/2022.07.16.499419

**Authors:** James Rosindell, Kerry Manson, Rikki Gumbs, William D. Pearse, Mike Steel

## Abstract

Phylogenetic metrics are essential tools used in the study of ecology, evolution and conservation. Phylogenetic diversity (PD) in particular is one of the most prominent measures of biodiversity, and is based on the idea that biological features accumulate along the edges of phylogenetic trees that are summed. We argue that PD and many other phylogenetic biodiversity metrics fail to capture an essential process that we term attrition. Attrition is the gradual loss of features through causes other than extinction. Here we introduce ‘EvoHeritage’, a generalisation of PD that is founded on the joint processes of accumulation and attrition of features. We argue that whilst PD measures evolutionary history, EvoHeritage is required to capture a more pertinent subset of evolutionary history including only components that have survived attrition. We show that EvoHeritage is not the same as PD on a tree with scaled edges; instead, accumulation and attrition interact in a more complex non-monophyletic way that cannot be captured by edge lengths alone. This leads us to speculate that the one dimensional edge lengths of classic trees may be insufficiently flexible to capture the nuances of evolutionary processes. We derive a measure of EvoHeritage and show that it elegantly reproduces species richness and PD at opposite ends of a continuum based on the intensity of attrition. We demonstrate the utility of EvoHeritage in ecology as a predictor of community productivity compared with species richness and PD. We also show how EvoHeritage can quantify living fossils and resolve their associated controversy. We suggest how the existing calculus of PD-based metrics and other phylogenetic biodiversity metrics can and should be recast in terms of EvoHeritage accumulation and attrition.

**Candidate cover image:** 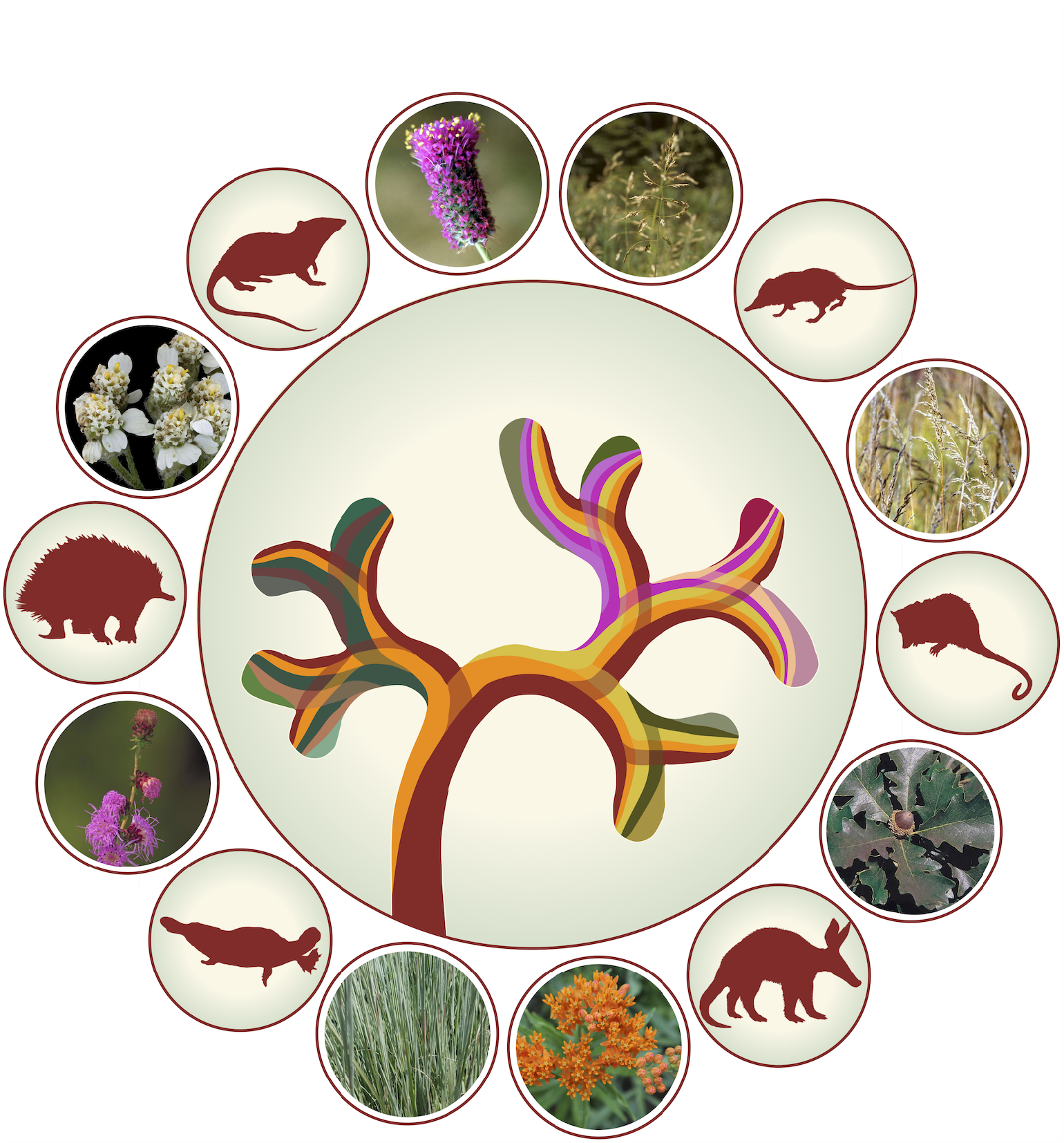

Artistic cover image prepared for this manuscript. The central tree depicts the gain and loss of Evolutionary Heritage (EvoHeritage) along each edge with its many coloured sections. EvoHeritage is proposed as an expansion of the concept of phylogenetic diversity. Around the outside of the tree are species that feature in our two practical applications of the EvoHeritage calculus: mammals identified as ‘living fossils’ and plants included in our study of community productivity. The cover image was generated by James Rosindell following discussions with co-authors. The Caenolestes outline (representing shrew opossums) and Dromiciops outline (monito del monte) used as components of this image are credited to Sarah Werning and provided under a CC BY 3.0 license; both images were recoloured in brown and placed over a shaded circle. All other images used as components are from the public domain. We thank Mina Mincheva for useful feedback on earlier drafts of the cover image.

## Introduction

Phylogenetic diversity (PD) is one of the most prominent measures of the variety of life, biodiversity. With biodiversity in crisis, PD was introduced by Faith (1992) to foster quantitative approaches to the problem of deciding where to focus limited conservation resources. It has since found many practical applications in conservation (Isaac et al., 2007; Gumbs et al., 2023) and as a biodiversity indicator, for example as highlighted by the Intergovernmental Science-Policy Platform on Biodiversity and Ecosystem Services (Brondizio et al., 2019). PD has become a mainstream feature of studies in ecology, evolution and conservation, and has been applied to taxa ranging from microbes (Bassett et al., 2015) to plants (Forest et al., 2007; Gonzalez-Orozco et al., 2016; Molina-Venegas et al., 2021) and vertebrates (Rosauer and Jetz, 2015; Gumbs et al., 2020).

PD has remained remarkably consistent with its original definition, which now underpins a large ‘calculus’ of related PD metrics (Faith, 2013), all derived from the same principles. The PD of a set of extant vertices (species) on a tree is defined as the sum of the lengths of all edges (branches) needed to connect those species (Faith, 1992) to the root vertex of the tree (though explicitly not right back to the origin of life; Faith and Baker, 2006). PD is typically measured on dated phylogenetic trees and therefore in units of millions of years, but could be measured on trees that are not ultrametric or that have other units. By taking edge lengths into account, PD goes beyond species richness and aims to capture biodiversity more completely including feature diversity, unknown features, and features with unknown future utility (future options for humanity) (Faith, 1992).

Alongside PD, other measures of biodiversity (Purvis and Hector, 2000a) such as functional diversity (Tilman et al., 1997) and trait diversity have also been broadly applied, with a myriad of different approaches (Petchey and Gaston, 2006). The question of how well PD captures feature diversity (and, relatedly, functional and trait diversity) has recently received increased attention from both data-driven (Mazel et al., 2017, 2018) and theory-focused (Wicke et al., 2021) studies. The answer undoubtedly depends on which features are being considered and how they are measured (Faith, 2018), leading to some debate (Owen et al., 2019; Mazel et al., 2019) and suggesting fertile ground for future research. However, there are also conceptual reasons why PD might not accurately capture feature diversity or future options for humanity.

Consider, for example, the features that evolved along the stem edge of all birds that connects them to their common ancestor with crocodilians (explicitly not the additional features that evolved later along the descendant edges). Notably, the feature of flight likely evolved along this stem edge. According to PD, all the features from the stem are captured by any one species of bird combined with any crocodilian because the edges needed to connect those two species must include the stem edge of birds. If we choose a flightless bird, however, this leads to a contradiction with the edge corresponding to the evolution of flight being fully ‘captured’ by a species that has lost its ability of fly.

An alternative to PD incorporating feature loss has been briefly considered, but it was concluded that PD was more suitable because of its simplicity (Faith, 1994b). It seems to have been assumed that the net effect of feature gains and losses is already reflected in edge lengths (Faith, 2018), or at least could be, with suitable changes to edge lengths.

There are, however, few options proposed for how to re-scale edge lengths (Faith, 1994c; Letten and Cornwell, 2015) and the majority of work continues to follow the convention of using edge lengths in terms of geological dates. Furthermore, whilst re-scaling would likely be an improvement, such an approach could not resolve the problem in our example. This is because the stem edge of birds would have to be shorter if we happened to be measuring a set of flightless birds, but not otherwise, so there isn’t a single length that we could coherently use to enable PD to capture the feature of flight correctly.

A related problem is apparent if we consider the PD of a mixed group of both extant and extinct species. The PD of a bird species and a crocodilian species together is not increased by the addition of a third extinct species that is a direct ancestor of birds exactly on the edge connecting them to the crocodilians (say the Archaeopteryx or a close relative). This is because the connecting edges of the two extant species already pass through the third extinct species in our example. Yet, in contradiction, we intuitively expect the extinct ancestor would hold features that have been lost in both of the extant species we have chosen. We cannot solve this problem by changing edge lengths because the extinct species will always be on the path between the other two species. By explicitly accounting for feature loss, however, the extinct ancestor may add to the diversity of extant species by representing lost features. Although extinct species may not seem relevant, inconsistency in such an example is likely to be a symptom of conceptual problems that are of more general concern. Furthermore, there are studies that calculate PD on trees including fossil taxa (Faurby et al., 2019) highlighting that phylogenetic diversity is already being used on mixtures of extinct and extant species.

In contrast to the problems posed by extinct ancestral species, homoplasy (convergent evolution) has long been acknowledged as a reason why PD may not capture feature diversity (Faith, 1992); two species with divergent evolutionary histories sometimes have similar features. This is not as troublesome for PD as one might assume because convergence may only apply to a (potentially superficial) subset of features. A blind snake has a remarkable resemblance to an earthworm, but closer inspection of their bodies would reveal the clear divergence of their less obvious features (Laver and Daza, 2021).

Convergent evolution requires two different ancestor species to have evolved to become similar in certain respects. This may occur as a result of acquiring the same feature independently more than once, but can also occur through the *loss* of a feature that once distinguished the two lineages. For example, the blind snake lost its sight and limbs along part of its path to convergence with the earthworm.

In the cases where homoplasy occurs without feature loss, the consequences for PD may also be less concerning because two (or more) independent, though convergent, solutions to the same adaptive need may indeed represent more underlying features and variety than one solution. We therefore suggest that the gradual loss (attrition) of features, and not homoplasy, is the real challenge for the classic PD narrative that has been left unanswered. Whilst we have illustrated this with obvious features such as flight and vision, we believe the same applies with equal force to other sources of heritable novelty and variation. The central thesis of this manuscript is that the attrition of such biological heritable variation affects PD, and other phylogenetic biodiversity metrics, in fundamental ways that cannot be fully accounted for by re-scaling edge lengths. We offer a solution in the form of a suite of new metrics that we illustrate with example applications: i) as a predictor of community productivity using data from the Cedar Creek grassland experiments (Tilman et al., 1996); ii) as a way to quantitatively identify so-called ‘living fossils’ (Grandcolas and Trewick, 2016) as species preserving the rarest ancestral features.

## Introducing EvoHeritage

We adopt the term EvoHeritage after ‘evolutionary heritage’ that was first coined by Mooers et al. (2005) to describe PD-based measures with a narrative that is more appealing for conservation initiatives working at the national level. However, we use the term for a new purpose: to highlight an important conceptual distinction between evolutionary history and evolutionary heritage. We take EvoHeritage to mean features including functions, future options for humanity, purely aesthetic or intangible features, and other sources of novelty and variation in the most general sense, but explicitly only the components that are heritable and present in a given set of organisms. These may be measured in a discrete or continuous manner. In contrast, evolutionary history may mean the total number of years over which lineages have been independently evolving, or the total EvoHeritage that ever existed in ancestors, importantly, this includes components that were since lost (see Fig. 1). We suggest that EvoHeritage is conceptually preferable to evolutionary history as a target for practical conservation. We avoid referring to EvoHeritage explicitly as ‘features’ so as not to confuse it with direct empirical measurement of an observed subset of features or traits and to frame EvoHeritage primarily as a conceptual generalisation of the PD calculus and with similar applications.

**Fig. 1:**
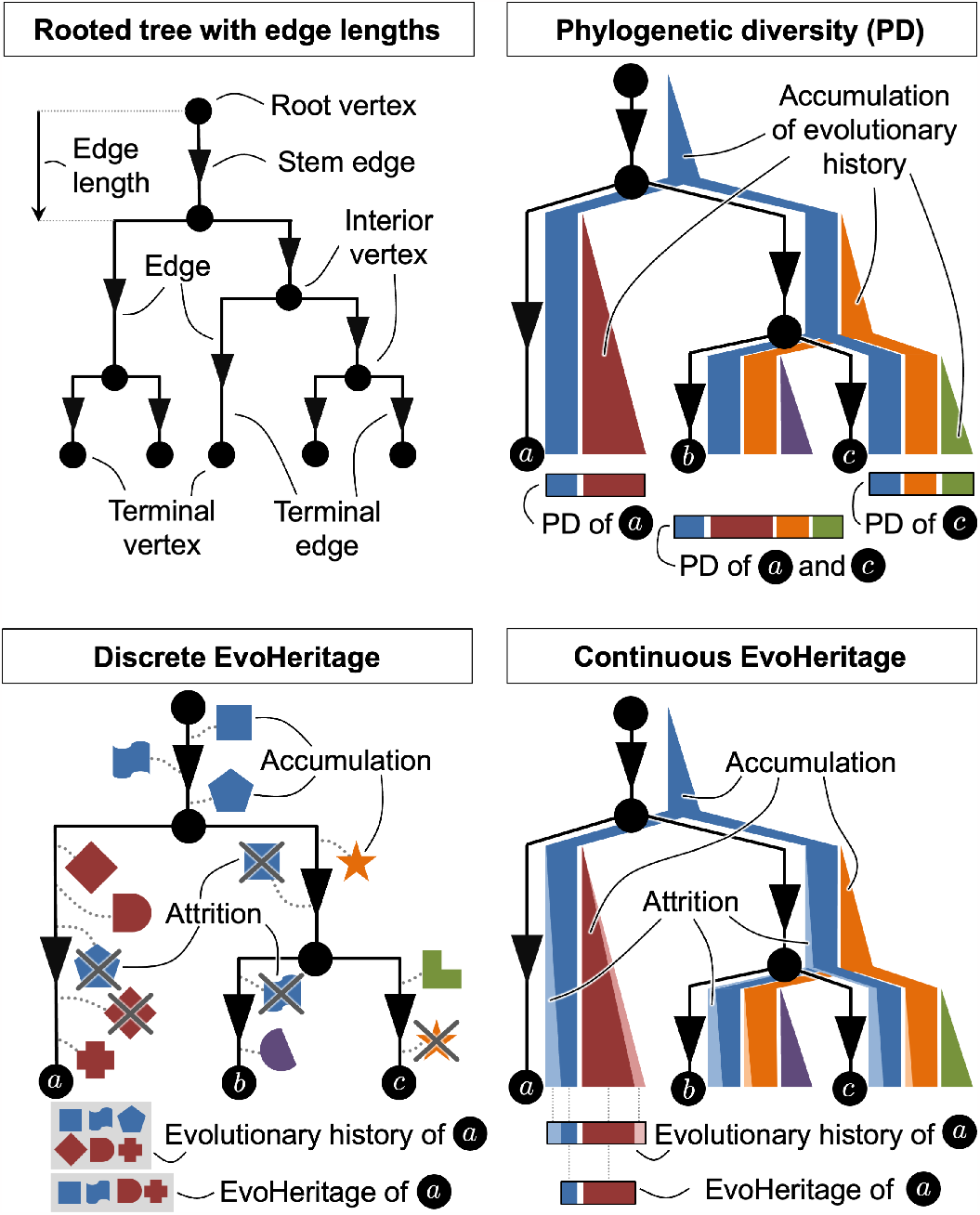
Visual glossary: an illustration of key terms and concepts from this manuscript. Panel a) shows tree terminology. Panel b) shows Phylogenetic Diversity as a continuous quantity framed as the accumulation of evolutionary history proportionate to edge lengths. Panel c) shows EvoHeritage framed as discrete components that are subject to accumulation and attrition, and distinct from evolutionary history. Panel d) shows EvoHeritage framed as a continuous quantity that is subject to accumulation and attrition, illustrating the distinction from evolutionary history. The concepts may straightforwardly be generalised to multiple ‘forms’ of EvoHeritage, each with their own rules for accumulation and attrition (see supplementary material).

We define ‘accumulation’ as a natural process by which novel EvoHeritage is generated along edges with copies being passed on to descendants. Conversely, we define attrition as a natural process by which EvoHeritage may be lost along the paths of a phylogenetic tree without extinction occurring (see Fig. 1). EvoHeritage attrition is akin to ‘substitution saturation’ that can be found at the genetic level, where the same areas of the genome can be overwritten multiple times. Substitution saturation leads to problems when building phylogenetic trees: mutations are not precisely cumulative, which results in skewing the apparent length of some particularly long edges (Xia et al., 2003). Similarly, attrition highlights problems with interpreting particularly long edge lengths in terms of heritable phenotypic change.

We define EvoHeritage in general terms to support various future applications (please refer to Fig. 1 in this manuscript and to the supplementary glossary https://doi.org/10.5061/dryad.z08kprrgs). We avoid referring to ‘species’ or ‘higher taxa’ and instead talk about the vertices of a tree. We imagine that ultimately species may equate to either a single vertex or a set of vertices. There may be different ‘forms’ of EvoHeritage potentially represented in an EvoHeritage tree (or graph), each form with its own characteristics of accumulation and attrition. For example, we may have ‘immortal’ EvoHeritage as one form that is never lost to enable rare and truly irreversible evolutionary steps to be accounted for. We consider only one form of EvoHeritage in the main text because to sum over multiple independent forms is straightforward, but to make it explicit would add complexity to the notation that follows.

### From tree to ‘EvoHeritage tree’

We define an EvoHeritage tree as a tree that specifies the accumulation and attrition of EvoHeritage explicitly along its edges. EvoHeritage trees are novel objects, so have to be constructed in a bespoke manner, depending on the research objectives and available data. This approach enables us to maintain conceptual separation between the calculations one can do on an EvoHeritage tree and the methods one uses to acquire the EvoHeritage tree to begin with. For each edge *e* we will need to calculate the net accumulation of EvoHeritage (which we write as *α*(*e*)) and the proportion of ancestral EvoHeritage to survive attrition (which we write as *β*(*e*)). The approach we follow for this is akin to the ‘Dollo model’ of binary characters that are gained and lost, but cannot be regained (Le Quesne, 1974), which has since also been applied to gene genesis and loss (Huson and Steel, 2004).

In our applications here, we build an EvoHeritage tree from a tree with edge lengths. We assume that EvoHeritage accumulates at a rate of *λ* units of EvoHeritage per unit of edge length. If edges correspond to time measured in millions of years, this means *λ* has dimension EvoHeritage *·* time^−1^. Attrition for a single unit of EvoHeritage occurs at a rate of *ρ* per unit of edge length and thus *ρ* would have dimension time^−1^. The total rate of attrition is thus proportionate to the total amount of EvoHeritage. These rates may be deterministic with EvoHeritage as a continuous quantity (‘standard conditions’) or may be stochastic with EvoHeritage as a discrete quantity (‘standard stochastic conditions’) see Fig. 1. The stochastic version naturally produces a distribution of outputs rather than a single value. This opens up the potential to study the properties of the resulting distribution, particularly the variance, which may be used as way of quantifying some of the potential error in using EvoHeritage as a proxy for feature diversity.

Consider a tree with specified edge lengths expressed with a function *L* defined so that *L*(*e*) gives the length of edge *e*. Attrition under standard conditions for a single unit of EvoHeritage occurs at rate *ρ* ⩾ 0 per unit of edge length (see Fig. 2). Therefore, the proportion of EvoHeritage surviving attrition, *R*(*l*), as a function of edge length *l*, satisfies the differential equation:

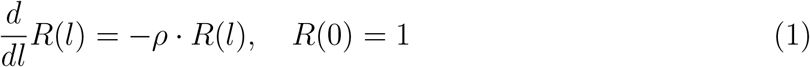

Solving this equation yields:

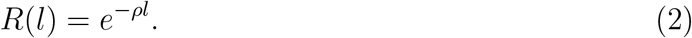

Note that here, *e* = 2.71828 … (rather than referring to an edge) and *ρl* is dimensionless because *l* has the same dimension as edge length (time) whereas *ρ* has the dimension of the reciprocal of this: time^−1^. We now derive *β*(*e*), which is also dimensionless, and gives the proportion of EvoHeritage that survives attrition along the edge *e*. We use *l* = *L*(*e*) in Eqn. (2) to write:

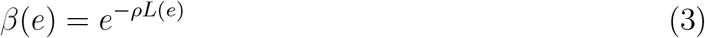

We now derive an expression for *α*(*e*), net accumulation of EvoHeritage, which includes EvoHeritage accumulation along the edge *e* but also the attrition of some of this EvoHeritage occurring later along the same edge. To do this, we integrate over edge *e*. This accounts for the EvoHeritage generated at every point along the edge, each having a different length over which it can be lost before reaching the end vertex:

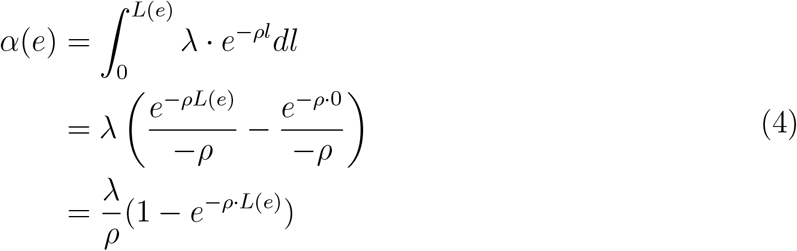

The dimension of *α* is thus simply EvoHeritage. For very small *ρ*, we can write 1 −*e*^−*ρ·L*(*e*)^ ≈ *ρ· L*(*e*). In the limit *ρ* →0 we can therefore say that *α*(*e*) = *λ· L*(*e*). This is also consistent with the exact expression of *α*(*e*) given by the first line in Eqn. (4) when *ρ* = 0.

**Fig. 2:**
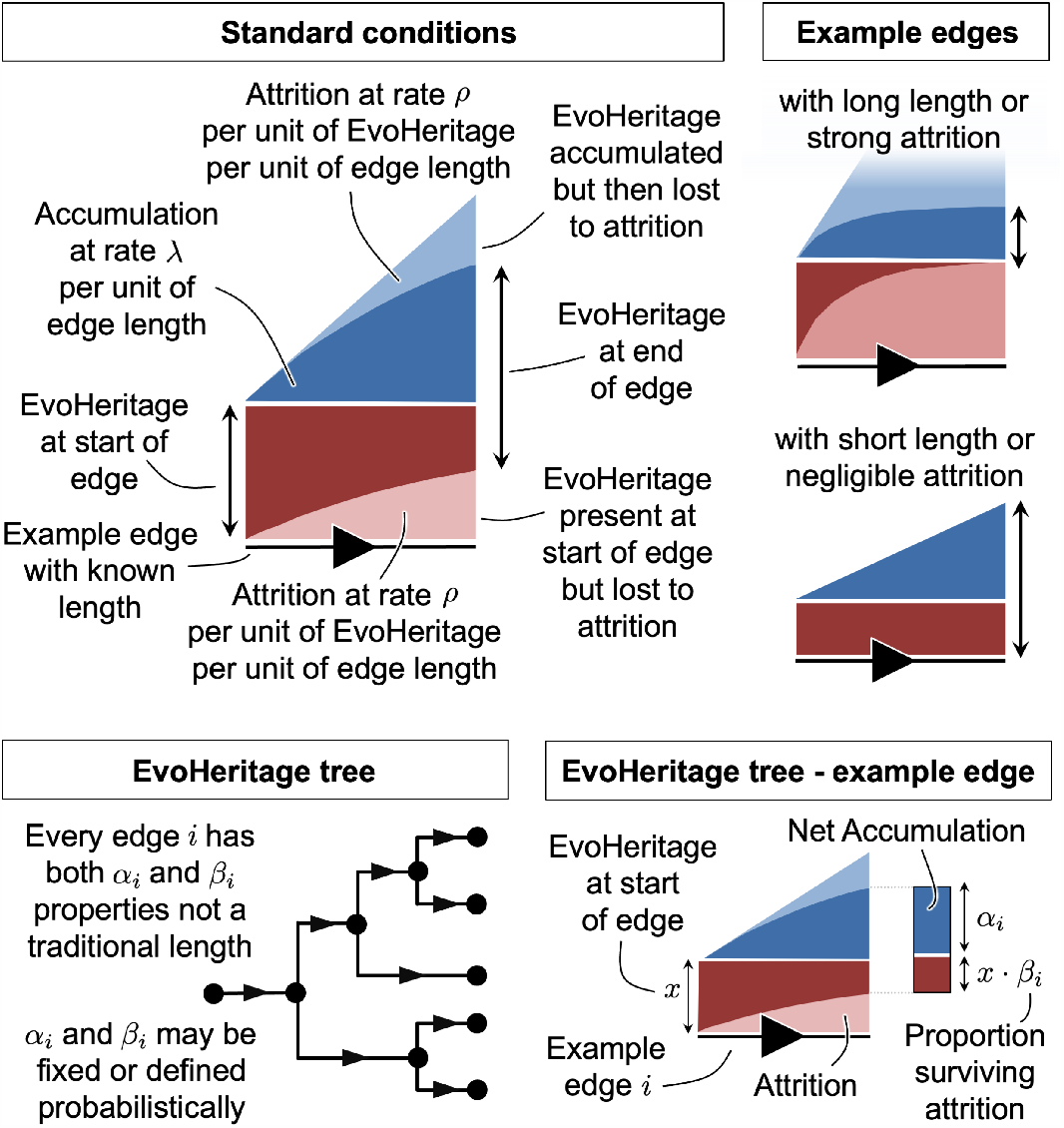
Standard conditions and the EvoHeritage tree concept. Panel a) shows standard conditions explained for a single example edge. Panel b) shows long and short extreme example edges under standard conditions. Panel c) shows an EvoHeritage tree and panel d) shows an example edge from that tree.

For a tree with edge lengths and values of *ρ* ⩾ 0 and *λ* ⩾ 0, we can now define an EvoHeritage tree under standard conditions by using Eqns. (3) and (4) for each edge of the tree. In case we are considering standard stochastic conditions *β*(*e*) denotes the expected proportion of EvoHeritage that survives attrition along the edge *e* and *α*(*e*) denotes the expected net accumulation of EvoHeritage, but the calculations are the same.

Edges where *ρL*(*e*) is small show very little or no attrition. In contrast, no ancestral EvoHeritage can survive attrition on the very longest edges (where *ρ· L*(*e*) is large). This means that net accumulated EvoHeritage eventually reaches saturation if an edge is sufficiently extended (see Fig. 2).

### Derivation of φ (total EvoHeritage) for a set of vertices on an EvoHeritage tree

In this section we give the formal derivation of *φ*, the total EvoHeritage of an arbitrary set of vertices (Eqn. 6 below) on a given EvoHeritage tree. If the values of *α* and *β* on the EvoHeritage tree have stochastic interpretations (as described earlier), then the same derivation would give *φ* as the *expected* total EvoHeritage. Throughout this manuscript, we describe an EvoHeritage tree mathematically by letting *V* represent the set of vertices and *E* be the set of directed edges (see Fig. 1). For a directed edge *e* = (*v*_1_, *v*_2_) in *E*, let *d*^*V*^ (*e*) = *v*_2_, the vertex descending from *e*, and let *a*^*V*^ (*e*) = *v*_1_, the immediate ancestor vertex to *e*. For a vertex *v* in *V*, let *D*^*E*^(*v*) be the set of edges descending from *v* and let *a*^*E*^(*v*) be direct ancestor edge to *v*. Thus *a*^*V*^ (*e*) = *v* if and only if *e ∈ D*^*E*^(*v*), and similarly, *d*^*V*^∈ (*e*) = *v* if and only if *a*^*E*^(*v*) = *e*.

To derive *φ* for an arbitrary set *X* ⊆*V* of vertices, we first iteratively calculate a term *p* (*v, X*) for every vertex *v* in *V* . The term *p*(*v, X*) represents the proportion of EvoHeritage associated with the vertex *v* that survives to be counted on at least one vertex in the set *X*. Our Eqn. (5) gives the formula for calculating these values, where the iteration proceeds in a way such that *p*(*v, X*) is calculated only after the values on all vertices descending from *v* have been calculated. The tree structure allows the iterative calculation of *p*(*v, X*) by giving a natural order to the calculations on each vertices.

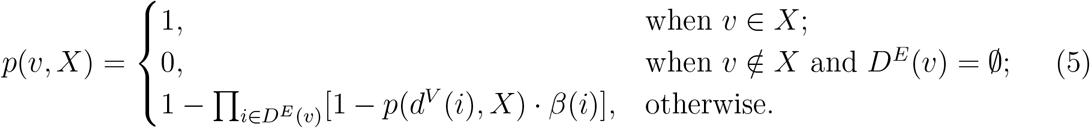

The first case in this Eqn. 5 terminates the iteration for vertices (usually terminal vertices) that are in set *X*. The second case terminates the iteration for other terminal vertices. The third case is for interior vertices only. It uses proportion *p* for all descendent vertices, and multiplies each by proportion of survival *β*(*i*) (see Fig. 2) along the edge *i* leading to that descendent vertex. Finally, the product term combines these to give the proportion of EvoHeritage survival down any descendent edge *i* ∈ *D*^*E*^(*v*). We are now in a position to write an expression for *φ* as a function of a set of vertices *X*.

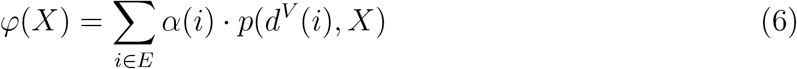

In words, the total EvoHeritage *φ*(X), of an arbitrary set of vertices *X*, is a sum across all edges in the EvoHeritage tree. Inside the sum is the EvoHeritage accumulated along each each edge *α*(*i*) multiplied by the proportion of that EvoHeritage surviving attrition to be counted in the set *X*.

### Derivation of φ_ρ_ for a set of vertices on a tree with edge lengths

We now define the quantity *φ*_*ρ*_, a special version of *φ* that can be calculated from a tree with edge lengths. This can be defined based on standard conditions or standard stochastic conditions. Under standard stochastic conditions, discrete EvoHeritage units are gained and lost along edges. We model this gain and loss of discrete EvoHeritage by a continuous time Markov chain. The model incorporates ‘constant birth’ and ‘linear death’ processes. The expected number of surviving EvoHeritage units can be calculated and it matches the deterministic expression for *φ*_*ρ*_ derived here based on standard conditions (see Supplementary Material).

We measure *φ*_*ρ*_ in terms of ‘standardised units’ defined as the total EvoHeritage at the end of a single disconnected edge of unit length. A standardised unit *S*_*u*_ is thus given by substituting *L*(*e*) = 1 into Eqn. (4) to give:

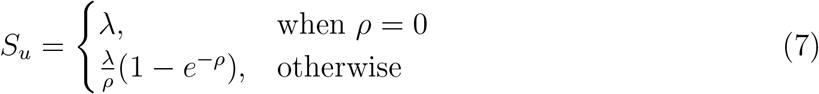

Thus, for a set of vertices *X*, the value *φ*_*ρ*_(*X*) is given by:

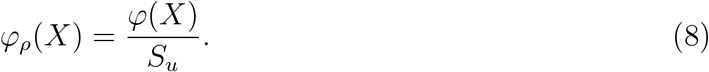

The quantity *φ*_*ρ*_ can be considered formally dimensionless because it is framed as the ratio between two components each of which have the same dimension (EvoHeritage). By substituting Eqns. (4), (6) and (7) into Eqn. (8) we can see that *φ*_*ρ*_(*X*) depends only on *ρ* and not on *λ*.

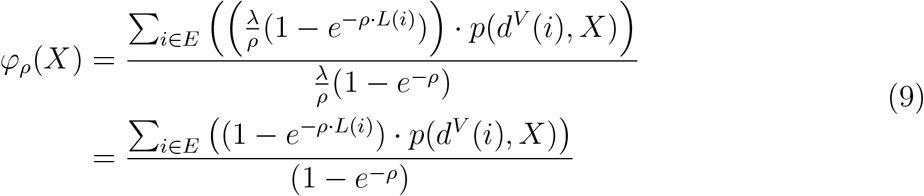

The quantity *φ*_*ρ*_(*X*) can therefore be used in place of PD in existing calculations on a tree with edge lengths for any given *ρ*.

### Relationship between φ_ρ_, PD and species richness

EvoHeritage expressed as *φ*_*ρ*_(*X*) converges to species richness (if the vertices represent species) when *ρ* → ∞; and *φ*_*ρ*_(*X*) converges to the PD of the set of vertices when *ρ* →0. We formalise this as a theorem the proof of which is given in supplementary material:

#### Theorem 1

Let *G* = (*V, E*) be an EvoHeritage tree, and let *X* ⊆ *V*.

1. lim_*ρ*→∞_ *φ*_*ρ*_(*X*) = |*X*|
2. lim_*ρ*→0_ *φ*_*ρ*_(*X*) = *PD*(*X*)

The convergence of *φ*_*ρ*_ to both PD and species richness may seem strange because it is dimensionless whereas PD is usually framed as having the dimension of time. However, one can represent classic PD as being technically dimensionless by re-scaling all edge lengths on a dated tree to multiples of any fixed, standardised length of time (Chao et al., 2010) and this is what we have effectively done in the limit when *ρ*→ 0.

In the ideal world, we would measure *φ*_*ρ*_ on a tree that includes the origin of life as a vertex *ô* (see supplementary material). This is a departure from the standard practice with PD, which does not trace back to the origin of life (Faith and Baker, 2006). However there reasons to include the origin of life: first, it enables results to be universally comparable between independent studies, being free from any bias brought by the size of tree for each study; second, it means the PD (or equivalently *φ*_0_) of a single vertex will be equal to the date of the origin of life. In our view this is more consistent with the concept of ‘evolutionary history’: any single extant species (or organism) arguably represents about four billion years of evolutionary history regardless of its position in the tree of life. We note that by adopting EvoHeritage, the deepest parts of the tree will have negligible effect apart from when we are very close to the PD limit (*ρ*→ 0). This is beneficial because it enables the origin of life to be included without any results being disproportionately affected by the large numbers involved.

### φ is not the same thing as PD on a tree with scaled edge lengths

Let us now ask whether *φ* can be considered as a restatement of PD with appropriately chosen edge lengths. To do this, we take an arbitrary EvoHeritage tree with a given topology and values of *α*(*e*) and *β*(*e*) on every edge *e*. We wish to find out if there exist some edge lengths for the tree that would enable PD to give us the same answer as *φ*. To formalise this, we say that *PD* corresponds to *φ* on an EvoHeritage tree *T* if we can calculate some edge lengths *L*(*e*) *>* 0 on *T* so that, for every possible set *X* of terminal vertices, *PD*(*X*) based on summation of the newly calculated edge lengths, gives the same value as *φ*(*X*) based on the original *α*(*e*) and *β*(*e*) values of *T* . If *β*(*e*) = 1 everywhere on *T*, there is no attrition and *φ* corresponds to *PD* by setting *L*(*e*) = *α*(*e*) for all *e*∈ *E*. If we had set *α*(*e*) and *β*(*e*) according to standardised conditions with *ρ* →0, then *β*(*e*) = 1 and *α*(*e*) = *λ· L*(*e*). The *λ* would be cancelled as we express the result in standardised units and hence *φ* would correspond to PD as we would expect from Theorem 1. Supposing that the special case of zero attrition (*β*(*e*) = 1 everywhere) does not apply (i.e. *ρ >* 0 if we’re using standardised conditions to get *α* and *β*), we can consider which arrangements of edges and vertices in *T* would nevertheless permit *PD* to correspond to *φ*.

#### Theorem 2

Let *T* be an EvoHeritage tree with terminal vertices given by *Y*⊂ *V* and *β*(*e*) *<* 1 for every edge *e* ∈ *E*. Then *φ*(*X*) = *PD*(*X*) for all sets *X* ⊆*Y* if and only if no edge is ancestral to more than two terminal vertices.

The proof of Theorem 2 follows as a consequence of a more general result proved in the supplementary material. In simpler terms, the theorem shows that PD can only correspond to *φ* on a tree when either attrition is explicitly turned off, or where the tree has a very simple shape: at most a polytomy at the root with each edge only branching at most once more (See Fig. 3). Such trees will not be encountered even in the simplest real-world applications. It can also be shown that a set (of a fixed number of vertices) chosen to maximise PD will not always capture the maximal *φ*_*ρ*_ although the difference may be small (see supplementary material). Another way to frame this is that PD no longer corresponds to *φ* if it is possible for accumulation and attrition to leave a non-monophyletic group representing any proportion of EvoHeritage (see Fig. 4). These findings are consistent with Wicke et al. (2021) who highlight that PD with suitable edge lengths can correctly capture feature diversity only when features appear in a pattern that could have arisen without any feature loss.

**Fig. 3:**
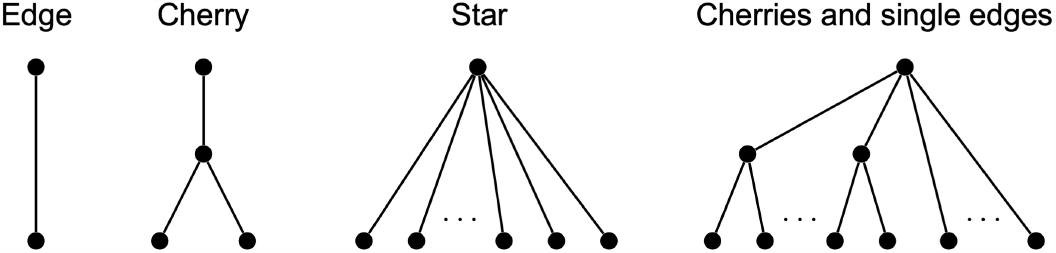
Characterisation of all tree topologies where PD corresponds to *φ*. This means that with EvoHeritage attrition occurring on every edge, we can define suitably re-scaled edge lengths so that, for every set of leaves, *φ* on the tree with attrition gives the same result as PD on the tree with re-scaled edges. These trees where PD corresponds to *φ* are very limited in complexity. If there is a stem edge leading down from the root then there are only two possibilities: a single edge or a ‘cherry’ (a stem and two descendent edges). In case there is more than one edge leading from the root (so the root is framed as a crown) the only possibilities are a ‘star tree’ (many edges connecting separately to the root) and a tree composed entirely of cherries and single-edges all connected directly to the root. PD does not correspond to *φ* on any other tree more complex than the ones shown.

**Fig. 4:**
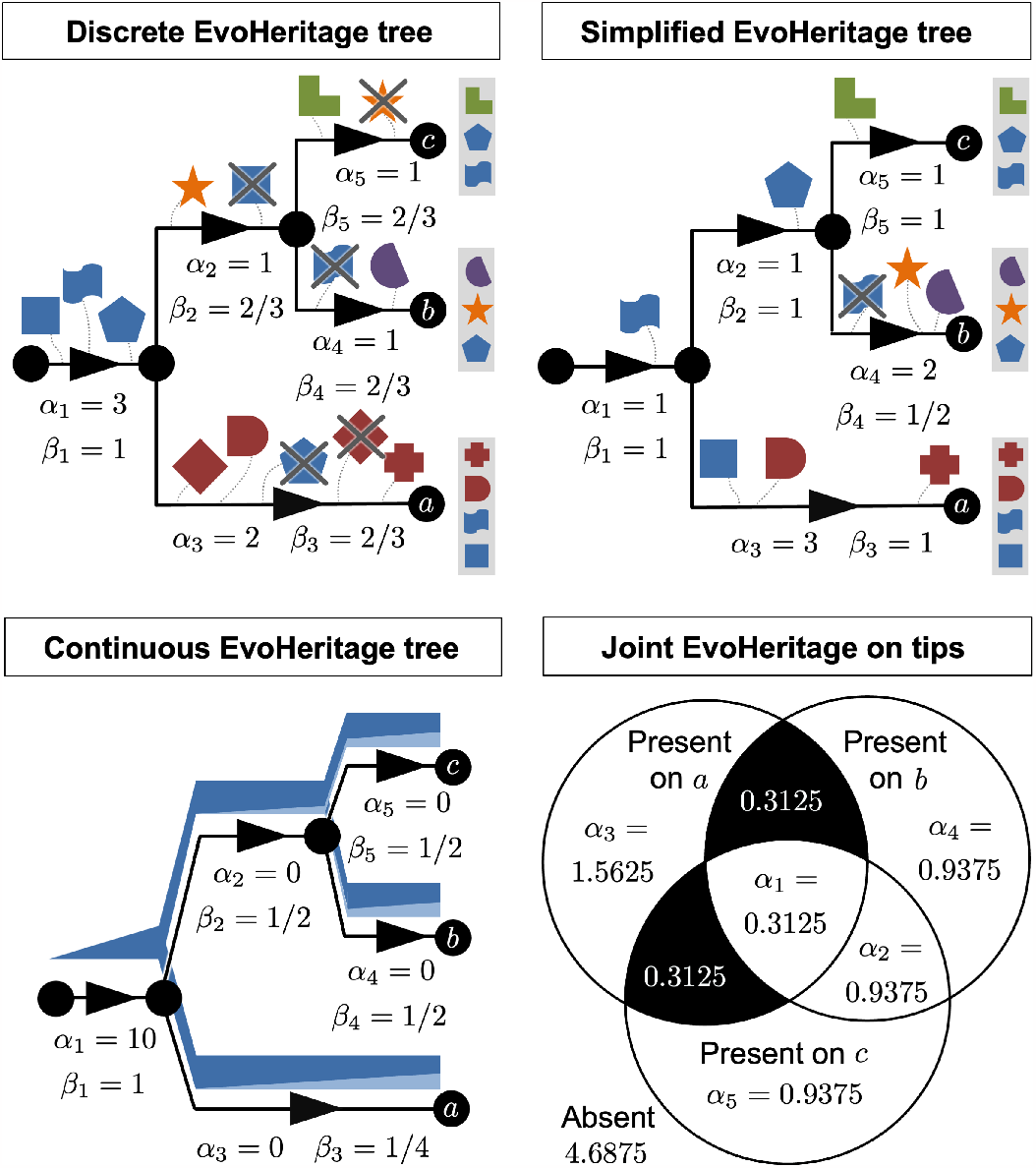
The fate of EvoHeritage on a simple tree to illustrate why it is impossible to frame accumulation and attrition in terms of PD on a tree with scaled edge lengths. Panel a) shows an example tree with discrete EvoHeritage units that are gained and lost on different edges. Each edge has a net gain of EvoHeritage, *α*, and a proportion of ancestral EvoHeritage that survives attrition, *β*. Of these, *α* is most like the classic edge length on a tree. Certain types of EvoHeritage loss can be absorbed by re-scaling edges (just in the *α* values). For instance, gain and loss of a discrete EvoHeritage unit along a single edge cancel out and shorten the effective length - this results in the *α* values on an EvoHeritage tree (e.g. the diamond on the terminal edge of vertex a. In other cases where EvoHeritage gain and loss leaves a monophyletic group with that EvoHeritage, we could express the gains and losses as a single gain at the stem of that group (e.g. the pentagon on the stem). Panel b) shows a new equivalent tree where some of the attrition (*β <* 1) has been eliminated by re-scaling of *α* values on each edge. We cannot eliminate the attrition of the EvoHeritage represented by the flag shape however, because this forms a non-monophyletic group. If we try to add it separately to the terminal edges of both vertex a and vertex c then it presents as homoplasy and we may double count the same EvoHeritage. If we try and compensate for the loss by reducing the *α* for the terminal edge of vertex b then we risk penalising for loss of an EvoHeritage unit that survives on another vertex. The same broad principles apply to continuous EvoHeritage e.g. under standard conditions. Panel c) shows an example of this with an EvoHeritage tree and panel d) shows an accompanying Venn diagram of the proportion of EvoHeritage shared between sets of terminal vertices. The values of *α* given in the Venn diagram are what we would need to assign to edges if we attempted to find an equivalent tree in which *β*(*e*) = 1 everywhere (i.e. a PD equivalent). Most sections of the Venn diagram (panel d) map to edges of the tree and to values of *α* but two shaded sections expresses a non mono-phyletic proportion of EvoHeritage that can never be eliminated or expressed in terms of *α* with *β*(*e*) = 1 everywhere.

## Developing a complete Evoheritage calculus

PD refers not only to the original definition (the minimum spanning tree of a set of terminal vertices together with the root) but also to a whole ‘calculus’ of derived measures based on the same underlying principles. To replicate the entirety of the PD calculus for EvoHeritage is beyond the scope of this work; however, we can begin the task and demonstrate that the rest will be achievable through incremental future work. For worked examples of parts of the EvoHeritage calculus, refer to Fig. 5.

**Fig. 5:**
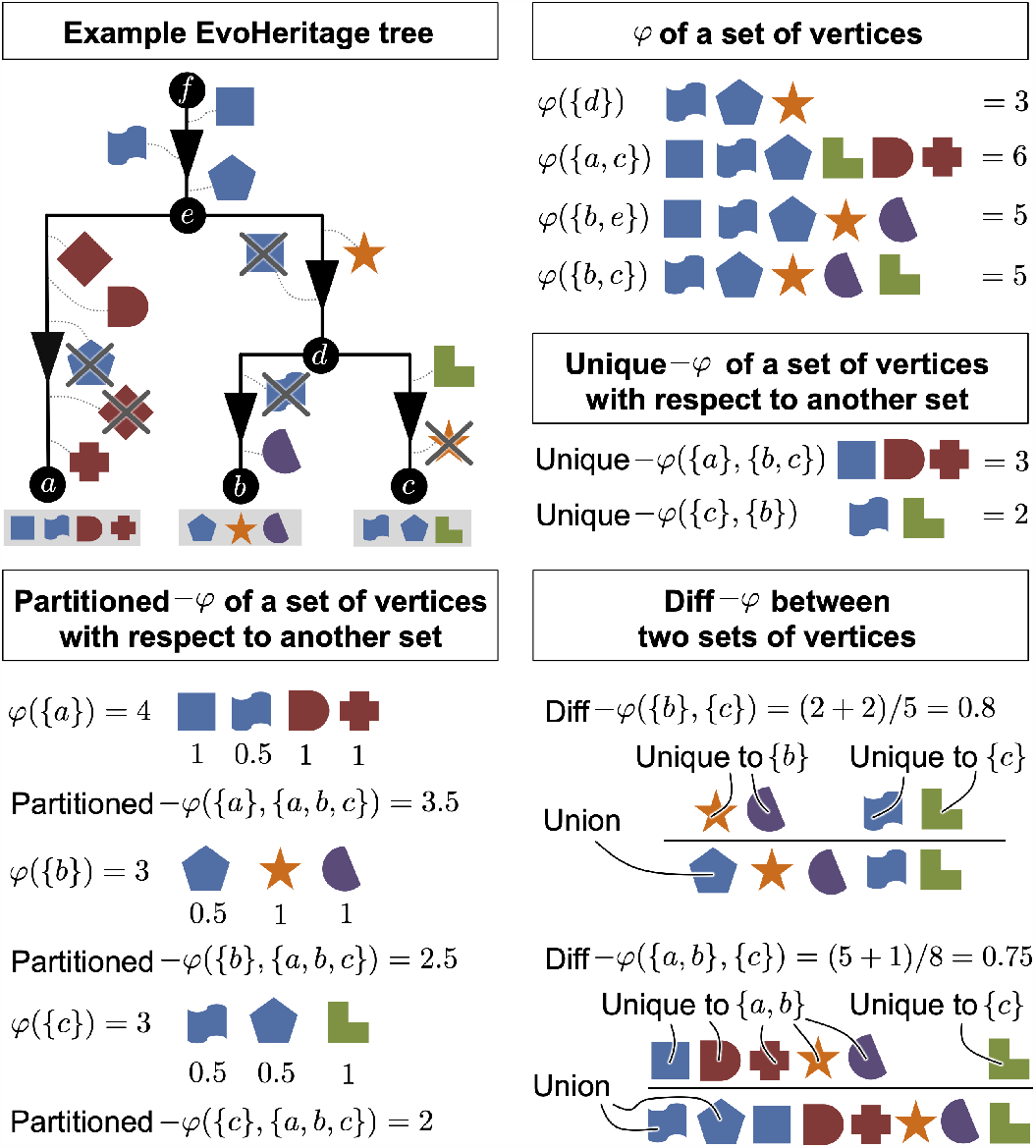
Illustrative examples of EvoHeritage, *φ*, unique-*φ*, paritioned-*φ* and diff-*φ*. We use discrete EvoHeritage units on the tree as this makes it easier to picture the underlying principles. If there were very large numbers of such discrete EvoHeritage units, the result would be a system where EvoHeritage is effectively a continuous quantity.

### Unique-φ

Unique EvoHeritage (unique-*φ*) is the EvoHeritage captured by a subset *X* of vertices and not captured by any of the vertices in a second subset *Y* of vertices. For most applications, we would expect researchers to choose that *Y* contains all extant vertices that are not in *X*. This is a generalised counterpart to ‘PD loss’ (Faith, 2013), also referred to as ‘unique PD’ and is calculated in a similar manner. The need to define the set *Y* more explicitly for unique-*φ* arises because extinct vertices are now potentially included. We define Unique EvoHeritage as follows:

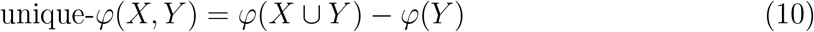

### EvoHeritage difference, diff-φ

The EvoHeritage difference (diff-*φ*) between two subsets (*X* and *Y*) of vertices is the EvoHeritage present within only one of the two sets, expressed as a proportion of the total EvoHeritage present in both subsets together. This can be written as follows:

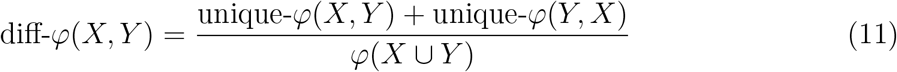

Note that diff-*φ* is symmetric so that diff-*φ*(*X, Y*) =diff-*φ*(*Y, X*). This metric is based on similar principles to the ‘UniFrac’ metric (Lozupone and Knight, 2005), which, in turn, is a phylogenetic version of the Jaccard index. Under standard conditions with *ρ*→ 0, diff-*φ*(*X, Y*) will converge precisely to the same result as UniFrac. The Phylosor metric of phylogenetic difference (Bryant et al., 2008) is similar to UniFrac but is based on the Sørensen index; an EvoHeritage counterpart to Phylosor along similar lines to diff-*φ* would be straightforward to develop.

In the supplementary material, we explore the relationship among diff-*φ*({*x}, {y}*), i.e. the EvoHeritage difference between two individual vertices, and phylogenetic difference between a pair of vertices (twice the date of common ancestry between a pair of extant vertices on a dated tree, or more generally the sum of all edge lengths along a path between the vertices). We also show that under standard conditions, diff-*φ* between a pair of vertices saturates as the length of the edges separating them increases (or more precisely as the product between *ρ* and edge length increases). Conversely, as *ρ* → 0, we see that diff-*φ*(*{x}, {y}*) converges to the sum of the edges required to connect *x* and *y* divided by a normalisation factor. In the other limit, as *ρ* → ∞, we see that diff-*φ*(*{x}, {y}*) converges to 1.

### Expected future-φ

The expected future-*φ* gives the expected value of *φ* at some point in the future, given *ϵ*(*v*), a probability of extinction by this future time point for each extant vertex *v* ∈ *V* where *D*^*E*^(*v*) =∅ . This can be considered a counterpart to expected PD. Calculation is straightforward, we simply cast the probability of extinction as an increased expected proportion of EvoHeritage loss on each terminal edge. This is done by subsituting *β*(*e*) with *β*(*e*) *·* (1− *ϵ*(*D*^*v*^(*e*))) for all terminal edges on an EvoHeritage tree then calculating *φ* on the new EvoHeritage tree.

### Relationship to genetic diversity

EvoHeritage has been constructed to avoid obligatory ties to the species concept. It therefore has some of the prerequisites to be applied in population genetics and for measuring both within- and between-species genetic diversity. There are many different measures of genetic diversity in general use (Hughes et al., 2008), and these may not necessarily fit the context of EvoHeritage. However, one early genetic diversity metric conceived by Crozier (1992) has a quantitative equivalent in terms of EvoHeritage. Crozier considered an unrooted tree with edge lengths between 0 and 1 corresponding to the probability of a difference being generated along the edge. This is directly comparable with our values of *β* (attrition) on each edge but without any values of *α* (accumulation). Crozier’s genetic diversity is the probability that all vertices will not be identical. If we add a stem to the tree in any location, to give directionality to the unrooted tree, then the proportion of the ancestral EvoHeritage accumulated on the stem, that does not survive attrition to exist on every extant vertex, is the same as Crozier’s genetic diversity (see supplementary material). It may seem counter intuitive that loss increases diversity, but attrition is interpreted as a multiplicative source of difference, and so not being inherited equally on all extant vertices is a source of variation and diversity. One could, perhaps, more closely equate this special case of EvoHeritage to ‘genetic *history*’ (Crozier et al., 2005) which is equivalent to the genetic diversity of Crozier (1992) but defined on a rooted tree.

### Power set approach

For a set *X*, we let 𝒫 (*X*) denote the power set of *X* (the set of all subsets of *X*). Here, we use the power set to introduce a new notation for EvoHeritage. We will see how this paves the way for a more elegant description of the broader EvoHeritage calculus, and for the use of Monte Carlo methods in numerical EvoHeritage calculations.

Let *T* = (*V, E*) be an EvoHeritage tree with vertex set *V* and edge set *E*. Intuitively, the functions we introduce here will be used to provide a measure of connection between vertices. The connection described is from a single vertex *v* ∈ *V* utilising only the edges in a particular subset of edges, say *G* ∈ 𝒫 (*E*). Formally, a function *F* of this type can be written as *F* : *V ×*𝒫 (*E*) *× H* → ℝ, where the set *H* varies depending on what we are trying to calculate. For example, if we seek to calculate *φ* using this approach, *H* = 𝒫 (*V*) and *X* ∈ *H* is a subset of vertices for which we are calculating *φ*. We now define a function *κ*(*F, X*) which calculates the percolation of EvoHeritage based on the measure of connection provided by *F* and a value of *X* ∈ *H*.

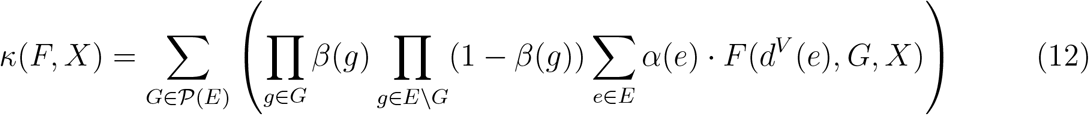

The calculation of *κ*(*F, X*) involves a sum over all possible subsets of edges in the tree. The broad idea is to view an EvoHeritage tree with attrition as a weighted distribution of trees (or disconnected sets of trees) without attrition. To see this in an example, consider a tree on which only one edge *e*_1_ has any attrition, so *β*(*e*) = 1, ∀*e* ∈ *E\ {e*_1_ *}*. We could express a calculation regarding this tree as a weighted average across two possibilities: first, with weight *β*(*e*_1_), a calculation of *F* on the tree with the edge *e*_1_ included (for EvoHeritage not lost along that edge); second, with weight 1 −*β*(*e*_1_), the same calculation of *F* on the tree (or disconnected pair of trees) that result from edge *e*_1_ being removed. Scaling up this thinking to a case where any possible subset of edges may be disconnected by attrition in different scenarios, we obtain the remainder of the terms given in Eqn. (12). We multiply factors of *β*(*e*) and 1 −*β*(*e*) for different edges to weigh across the possible combinations of attrition. The format of the function *κ* shows how the sum over 𝒫 (*E*) can be replaced with a Monte Carlo method for numerical approximation (see empirical applications – methods).

We will now describe a specific choice of function for *F* so that *κ*(*F, X*) returns *φ*(*X*) as derived earlier (Eqn. 6). For a set of edges *G* ∈ *𝒫* (*E*), we wish to count the number of different vertices *x* in set *X* ⊆ *V* that can be reached from the given vertex *v* ∈ *V* using only edges from within *G*. We introduce a function 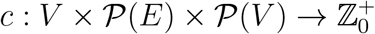, given by:

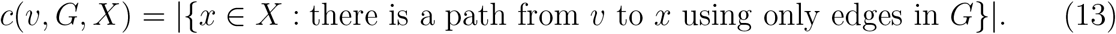

To differentiate between the sets of edges which contain at least one path from *v* to *X* and those that contain no such path, we introduce an ‘indicator function’, ℝ : *V ×*𝒫 (*E*) *×*𝒫 (*V*) → *{*0, 1*}*, defined by:

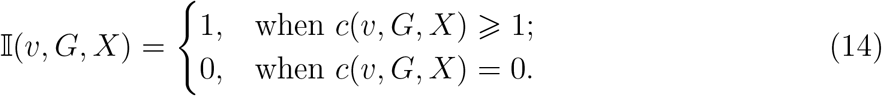

We can now express *φ* in terms of *κ* and ℝ.

#### Theorem 3

For all subsets *X* ⊆ *V, φ*(*X*) = *κ*(ℝ, *X*).

For a proof of Theorem 3, refer to the supplementary material.

### Partitioned-φ

PD can be partitioned among terminal vertices with evolutionary distinctiveness (also called fair proportion) or alternatively with equal splits (Redding, 2003; Redding and Mooers, 2006; Isaac et al., 2007). Fair proportion and Evolutionary Distinctiveness (hereafter ED) follow the same approach by sharing the length of each edge evenly between all extant descendant vertices regardless of tree topology (Redding, 2003; Isaac et al., 2007). Here, we provide a corresponding partitioning for EvoHeritage. We seek to fairly share out (partition) the ‘value’ of all EvoHeritage present on a set of vertices *Y* among the subsets of these vertices. Consider a subset *X* of *Y* . The partitioned-*φ* of *X* with respect to *Y* (denoted by partitioned-*φ*(*X, Y*)), is the fair share of the value *φ*(*Y*) which is attributed to the vertices in *X*. To calculate this fair share, we need to account for the number of copies of EvoHeritage across all vertices of *X* and across all vertices of *Y* (Fig. 5). The power set approach provides an elegant solution because attrition is accounted for automatically by weighted averaging in the function *κ*. We only need to write a function to partition the value of the EvoHeritage from one vertex passing through a known subset of edges.

More precisely, suppose that *X* ⊆ *Y* . The function *R*(*d*^*V*^ (*e*), *G, X, Y*) (defined in Eqn. 15) as follows:

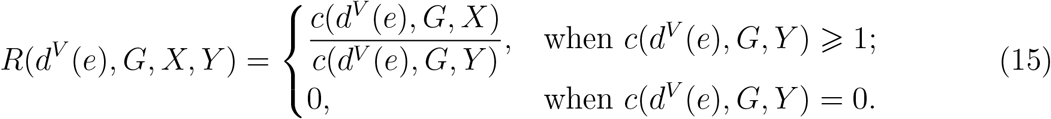

gives the number of vertices in set *X* that can be reached from edge *e* using only edges in *G* as a proportion of number of vertices in set *Y* ⊇ *X* that can similarly be reached from edge *e* using only edges in *G*. This can be framed as the fair share of EvoHeritage present in *Y* that is attributable to vertices of set *X* ⊆*Y* . Applying *κ* to the function *R* gives us partitioned-*φ*(*X, Y*).

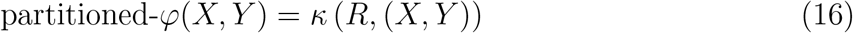

Many applications of partitioned-*φ* will use it to assign a value to each individual extant vertex *x ∈ X* within the context of all extant vertices. In supplementary material we show that the sum of all partitioned-*φ* values returns the total EvoHeritage of all extant vertices together: *φ*(*X*) = Σ_*x* ∈ *X*_ partitioned-*φ*({*x}, X*). Under standard conditions, partitioned-*φ*_*ρ*_ of a single extant vertex within the context of all extant vertices will converge precisely to ED as *ρ* → 0 (see supplementary material).

## Empirical applications –Methods

In this section, we will introduce two different empirical applications of our work that both rely on attrition: (i) an ecological application evaluating *φ*_*ρ*_ as a predictor of community productivity, (ii) an evolutionary application using the concepts of attrition and partitioned EvoHeritage to quantify living fossils. The results and discussion of each application will be given in later sections.

### i)Evaluating φ_ρs_ as a predictor of community productivity

The field of biodiversity-ecosystem function has yielded important insights for conservation, perhaps the most fundamental being that, in general and all other things being equal, greater biodiversity is associated with higher ecosystem function (Balvanera et al., 2006; Grace et al., 2016). A major contribution of phylogenetic conservation has been the general trend of phylogenetic diversity to correlate strongly with ecosystem productivity and, in notable cases, to do so more strongly than species richness or even functional diversity (Cadotte et al., 2009; Tucker et al., 2019). Perhaps the most influential study of these kinds of effects, by Cadotte et al. (2009), evaluated a range of diversity metrics, including species richness, PD and functional diversity. Using biomass and species lists from each of the 13m *×* 13m plots of the Cedar Creek Biodiversity Experiment (itself an iconic system in this strand of the literature; Tilman et al., 1996, 2001), the authors ranked 16 different individual predictors by their ability to explain plot-scale community productivity, measured as biomass accumulation. PD was second only to the presence of nitrogen fixers, and was an improvement over both species richness and a range of other functional diversity measures.

We repeat parts of the analyses of Cadotte et al. (2009), introducing *φ*_*ρ*_ alongside the PD and species richness predictors. One aim is to demonstrate our theoretical results (Theorem 1) with empirical data, showing that *φ*_*ρ*_ scales naturally as expected between species richness and PD, depending on the strength of attrition. Another aim is to showcase the possible insights that may follow from seeking a value of *ρ* to optimise some payoff (such as the ability of EvoHeritage to act as a proxy for feature diversity).

We sourced the Cedar Creek Biodiversity Experiment data from (Tilman, 2021) to provide both plot biomass and a matrix of species presence by plot, and the phylogenetic tree directly from Cadotte et al. (2009). All analyses were performed in R version 4.1.2 (R Core Team, 2021). We used the chronos function within the ape R package version 5.6-2 (Paradis et al., 2004) to make the tree ultrametric, similar to the original study (Cadotte et al., 2009). The tree was rate-smoothed (Sanderson, 2002) and dated by setting the crown, a common ancestor with *Amborella trichopoda* at 234.5 million years ago following (Zanne et al., 2014). The total tree depth was 1 in Cadotte et al. (2009), but the effect of our change is only a linear scaling in edge lengths. We note that when calculating *φ*_*ρ*_, edge length only appears in Eqns. (3) and (4) as a product with *ρ*, which we are going to vary. As a result, adding dates to the tree will not change any of our main results, it will only linearly scale the values of *ρ* at which we see those results, making the *ρ* values comparable to those calculated on other dated trees. We developed an R function to iteratively calculate *φ*_*ρ*_ under standard conditions using Eqns. (6), (3) and (4) (code available on GitHub - see later). We then applied this function to calculate *φ*_*ρ*_ for every one of the empirical plots. We repeated this process for a range of attrition parameters between *ρ* = 10^−4^ and *ρ* = 10^4^. To ensure we would see both the PD and species richness limits, and to help verify the correctness of our new function, we also calculated PD and species richness for each plot directly using the *pez* package version 1.2-3 in R (Kembel et al., 2010; Pearse et al., 2015).

Given that the phylogeny of species present in the Cedar Creek plots has a single deep split dividing the true grasses Poaceae (8 species) from 10 other species that are all within the Pentapetalae, a clade of Eudicots (see Fig. 6), we calculated the same diversity metrics for just the Poaceae and again for just the Pentapetalae as well as for both groups together. We calculated a Pearson correlation coefficient separately for plot biomass against every one of our plot diversity measures: PD, species richness and *φ*_*ρ*_ with each value of *ρ*, for all plants, for Poaceae only and for Pentapetalae only. We also repeated the whole analyses on filtered community data that incorporates only plots containing at least one representative from each of Pentapetalae and Poaceae. A sample of the data, together with the methods workflow, can be seen in Fig. 6. We discuss our findings in the Results section.

**Fig. 6:**
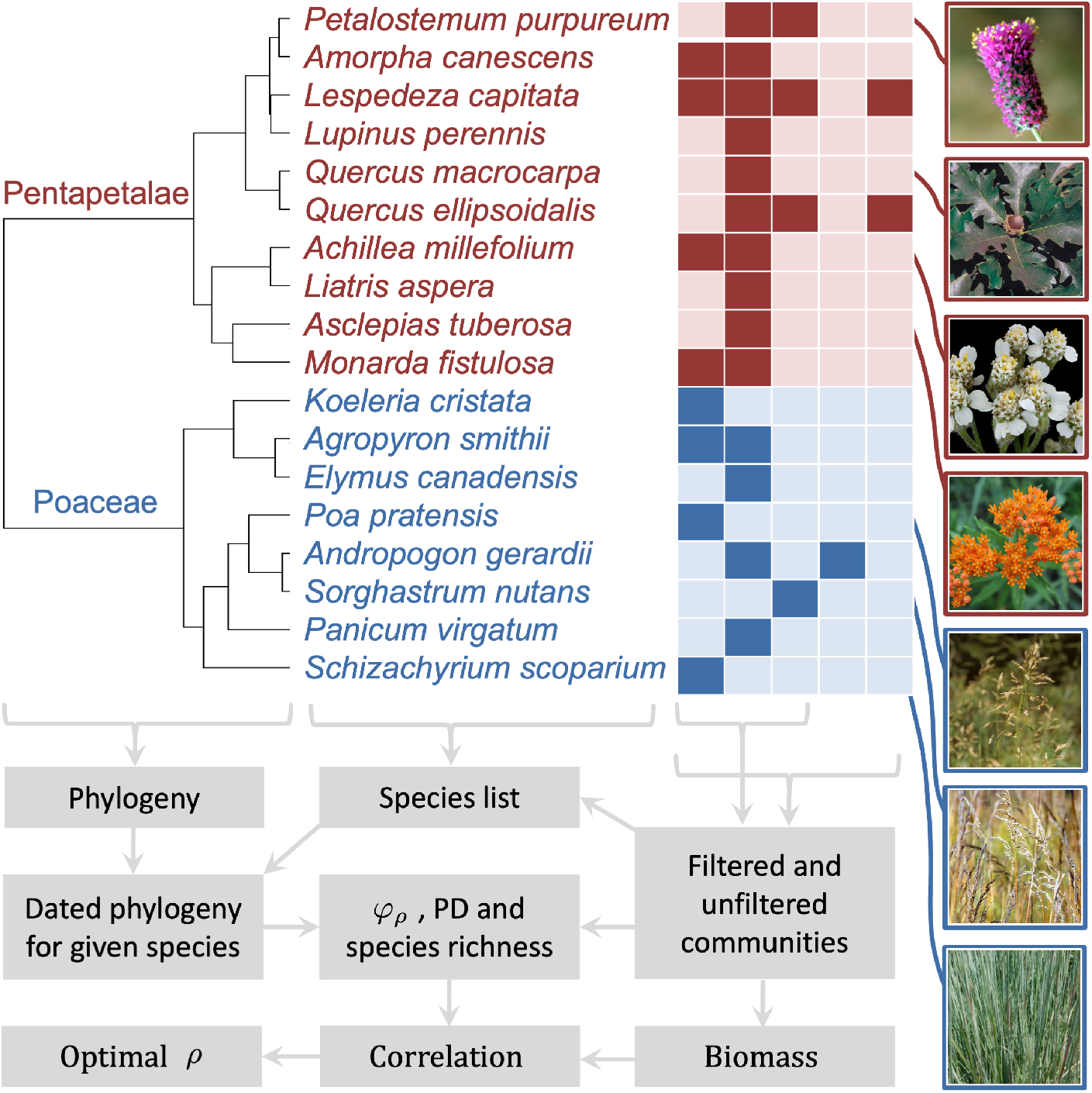
Selected empirical data and workflow for our community productivity example application. The phylogeny shows all species present in the study with edge lengths to scale, including the deep split between Pentapetalae and Poaceae. Next to the phylogeny we show a sample of the community presence-absence data across five communities (columns). Presence is indicated by a solid box, absence is indicated by lighter shades. Public domain images of selected species are shown. The filtered community data excludes communities that lack any representatives of either Pentapetalae or Poaceae. The flowchart shows the calculation steps leading to an optimal *ρ* for each scenario as the end product.

### ii)Applying attrition and partitioned-φ to quantify living fossils

The idea of ‘living fossils’ is contentious (Casane and Laurenti, 2013; Bennett et al., 2018; Lidgard and Love, 2021). Proponents of the concept say that living fossils are useful as motivation for a wide range of research questions (Lidgard and Love, 2018) and as a focus for conservation attention (Turner, 2019). One reason for the contention is perhaps the lack of a clear definition of a ‘living fossil’ and the wide range of different identifying characteristics that are mentioned across numerous studies (Bennett et al., 2018).

Commonly claimed features of living fossils are slow rates of evolution, “gross similarity to an ancestral fossil” (Lidgard and Love, 2018, 2021) and long periods of morphological stability (Turner, 2019). Opponents argue that these characteristics are in conflict with mainstream evolutionary thinking (Casane and Laurenti, 2013) and reinforce misinterpretations about evolution because they appear to support the idea of a ‘ladder of progress’ (Omland et al., 2008), where some extant species can be considered more ‘primitive’. It is also argued that even the most iconic living fossils, such as the coelacanths, do not really display morphological stability or slow rates of change at all (Casane and Laurenti, 2013). These are strong arguments, but, at the same time, it is hard to simply dismiss a large body of work on living fossils outright. It should be possible to distil some special property that connects all those species and clades that have been called living fossils.

Some attempt has been made to resolve the controversy surrounding living fossils by recasting them as ‘relict species’ whilst giving warnings about misinterpretation (Grandcolas and Trewick, 2016). In contrast, Bennett et al. (2018) argue that a quantitative concept of living fossils is needed to underpin research and resolve some of the controversy. They went on to describe the ‘evolutionary performance index’, which, to the best of our knowledge, remains the only pre-existing quantitative index focused on living fossils. Whilst evolutionary distinctiveness (ED) has also been considered as a potential indicator for living fossils (Isaac et al., 2007), it was raised primarily to defend high ED species against the idea that they have lower evolutionary potential (Erwin, 1991). The evolutionary performance index focuses on monophyletic clades and consists of three components: species richness, clade age and the number of evolutionary and ecological changes since divergence. Bennett et al. (2018) were emphatic that their approach does not invoke a ladder of progress. However, for living fossils to be defined partly on the basis of having few morphological and ecological changes since divergence still feels to us reminiscent of the ladder’s problematic characteristics. We will apply the idea of attrition and EvoHeritage to conceptualise living fossils and thereby show how the phenomenon of living fossils can be explained from tree topology alone without the need to invoke any heterogeneity in rates of genetic or morphological change. We will see that whilst our approach reduces to ED in a special case, it bears a better conceptual and mechanistic link to living fossils away from this special case.

We propose that the taxa (say, the extant vertices of a dated EvoHeritage tree) called living fossils do not possess more ancestral features *in total* than other extant vertices; rather, the features they have are *rarer*, or even *unique* to the living fossil among the extant vertices. EvoHeritage includes more than the morphological features that are typically preserved in fossils, but we assume that EvoHeritage can be used as a proxy for these features. Consider EvoHeritage accumulation and attrition occurring together at an equal pace across every edge in a dated EvoHeritage tree (under standard conditions). Now consider an ancient vertex with two descendant edges, one leading to a single extant vertex (a candidate living fossil) and the other leading to a diverse group of extant vertices.

Because attrition is constant everywhere, all extant vertices will inherit an equal total amount of ancestral features. The candidate living fossil vertex, however, inherits an independent sample of ancestral features, meaning those features are more likely to be rare or unique. In contrast, the vertices from the diverse sister group share a greater amount of evolution with each other, meaning many of the features they inherit, surviving attrition, will also tend to be shared and thus will not be rare.

Let us now explore how EvoHeritage concepts can be used to quantitatively identify living fossils from first principles on a dated EvoHeritage graph. One founding principle is that being a living fossil is not a binary categorisation; it is a continuous metric (Bennett et al., 2018; Turner, 2019). We use the term ‘living-fossil-ness’ first coined by Bennett et al. (2018). However, our approach beyond this is very different from that of Bennett et al. (2018).

In practice, clades containing more than one species may be identified as clades of living fossils (Bennett et al., 2018). For example, the coelacanths (Latimeria) consist of two known extant species. We consider only individual extant vertices on a species-level tree as the candidate entities. Although closely connected groups of vertices might happen to all be living fossils, this should be viewed as an emergent property from calculations that work at the level of a single vertex. EvoHeritage graphs intended for living fossil identification should thus be constructed so that all extant vertices are at comparable taxonomic levels in order to avoid bias in the results.

Fossils come from many different periods in geological history and may be preserved in different patterns on a dated EvoHeritage graph. The features represented in these fossils have also originated at different points in evolutionary history, but not later than the date of the fossil itself. We introduce the concept of a ‘targeted EvoHeritage tree’. This is a dated tree on which attrition, *β*(*e*), is defined under standard conditions for each edge whilst *α*(*e*) is augmented to include accumulation of only that EvoHeritage originating during a specified period of geological history. For example, if we want to consider living fossils from the Cretaceous then we only accumulate EvoHeritage from 66 million years ago or earlier because any EvoHeritage accumulated more recently than this could not possibly be represented in a fossil from this period. We could go one step further and only count EvoHeritage that arose between 66 and 145 million years ago. This means we’re looking specifically at features that appeared for the first time in fossils during the Cretaceous, a more specific case. We quantitatively define the ‘living-fossil-ness’ of an extant vertex *v* on a targeted EvoHeritage graph as log_10_(partitioned-*φ*({*v}, X*)) where *X* is the set of all extant vertices and *v* ∈ *X*. Attrition will typically be an exponential process, so we take the log of partitioned-*φ* to make the comparisons more natural. Our living fossil concept and associated methods workflow are described in Fig. 7.

**Fig. 7:**
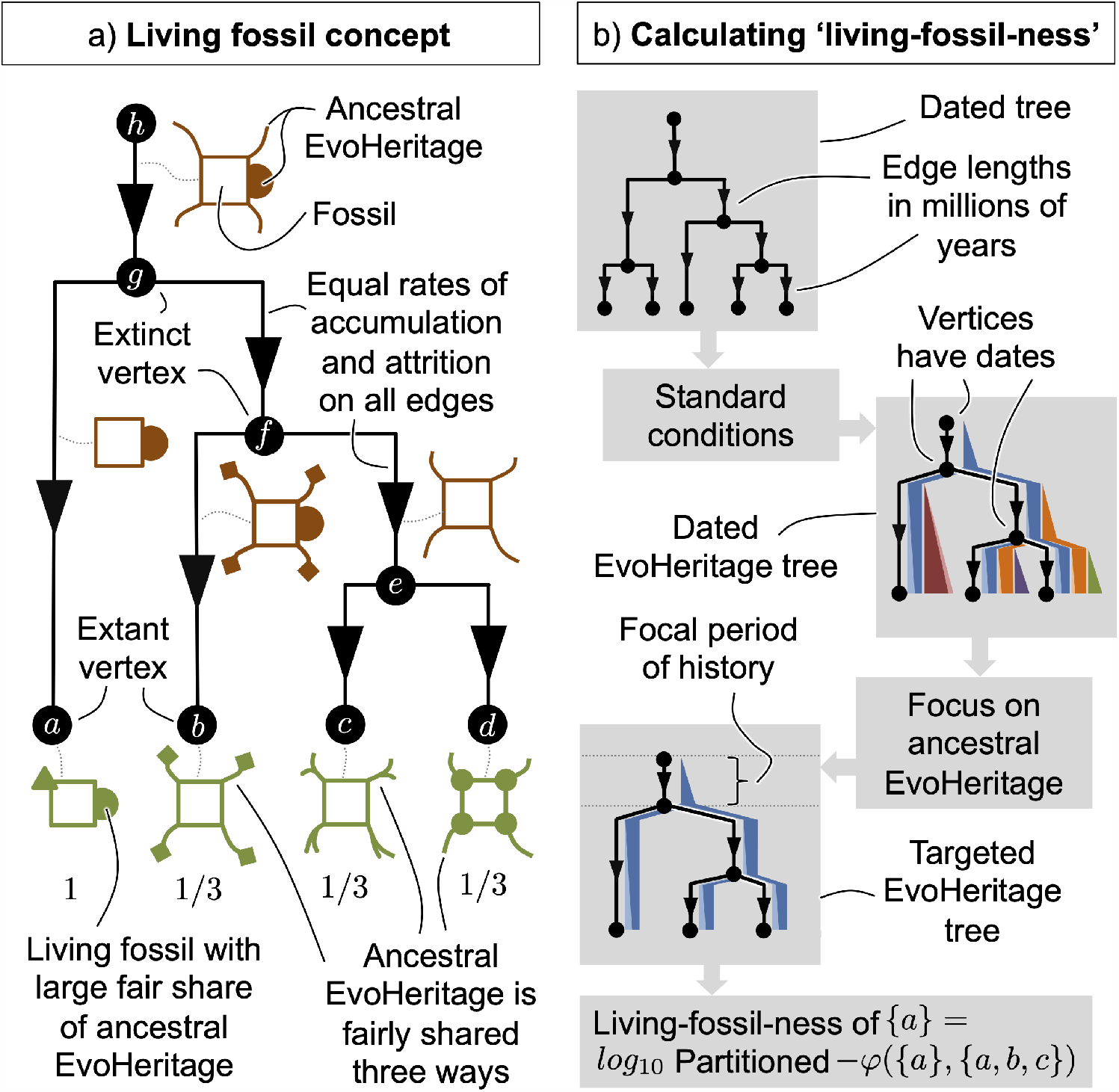
Panel a) shows an Illustration of the living fossil concept from the perspective of EvoHeritage. Each of the four extant vertices retains an ancestral feature, loses an ancestral feature, and gains a novel feature, in this sense they are equal. A fair partitioning of ancestral features, however, reveals an outlier with a larger share of ancestral features that could also appear in the fossil record. Panel b) shows our workflow for calculating living-fossil-ness quantitatively. EvoHeritage accumulation is deliberately suppressed other than during a specified period of history from which we are considering fossils - in this illustration, the stem edge. The features being partitioned are therefore only those that would have arisen on the stem, and thus been present in species being fossilised during this period. The period of interest could include everything from the origin of life to a key extinction event - to partition features present in fossils from this event. Alternatively, it could include just a key period of geological history - to partition features that first appeared in fossils during this period.

Our approach may appear superficially similar to ED, but there are important differences. First, we consider only EvoHeritage that may have been present in fossils from a particular period, likely an ancient period to match the narrative around living fossils. ED, in contrast, partitions all sections of edge in the same manner, even the most recent ones. Secondly, whilst partitioned-*φ*_*ρ*_ does converge to ED as *ρ*→ 0, we believe that *ρ >* 0 is the more realistic scenario.

Our method requires no data beyond a phylogenetic tree with edge lengths, and a value of *ρ*. We sourced mammal trees as a posterior distribution from Upham et al. (2019) with edge length units in millions of years. To construct targeted EvoHeritage graphs for each scenario, we first calculate *β*(*e*) under standard conditions using Eqn. (3). To calculate the net EvoHeritage accumulation *α*(*e*) for an edge spanning time the time period from *ℓ*_1_ to *ℓ*_2_ million years ago under a scenario of EvoHeritage accumulation between *t*_1_ and *t*_2_ million years ago there are several cases. Bearing in mind that *t*_1_ *< t*_2_ and *ℓ*_1_ *< ℓ*_2_, we have

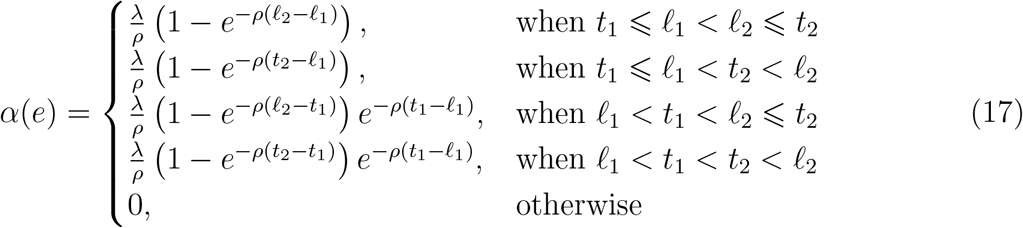

The first case is where EvoHeritage accumulates as normal along the edge. The second case is where the edge continues further back in time than the end of the EvoHeritage accumulation period, and so the formula remains the same, but the duration of accumulation is shorter than the length of the edge. The third and fourth cases mirror these concepts but the edge begins more recently than the start incorporate a period of EvoHeritage accumulation, and so there is a period of attrition only (without accumulation) along this part of the edge to be accounted for as well. The last case occurs otherwise, in practice when *ℓ*_2_ *< t*_1_ and the edge ends before the period of EvoHeritage accumulation or when *ℓ*_1_ *> t*_2_ and the edge starts after the period of EvoHeritage accumulation - in either case the entire edge lies outside the targeted accumulation period.

To illustrate our ideas, we will detect mammalian living fossils for three periods in geological history: the Jurassic (EvoHeritage accumulation from 145 million years ago or older), the Cretaceous (EvoHeritage accumulation from 66 million years ago or older) and the Quaternary (EvoHeritage accumulation from any point in time).

We calculate partitioned-*φ*_*ρ*_ on the resulting targeted EvoHeritage tree for every extant vertex using Monte Carlo methods. To see how, consider Eqn. (12): the sum over the power set 𝒫 (*E*). For this calculation, both of the product terms involving functions of *β* will be replaced with a repeated stochastic process that considers each edge *e* of the

EvoHeritage graph and disconnects it with probability 1− *β*(*e*), then calculates the inner part of the expression from Eqn. (12). After substituting this and Eqn. (15) into Eqn. (16), then reordering the two summations, we can write:

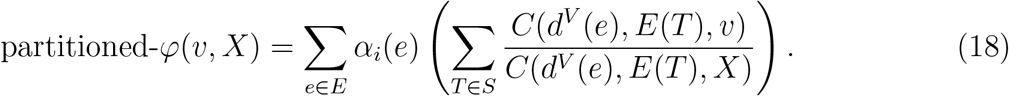

In this equation, *S* is the set of EvoHeritage trees resulting from repeated random removal of edges from the edge set *E*, and *T* is an example EvoHeritage tree from the set *S*, and *E*(*T*) is the set of all edges remaining in *T* . Notice from Eqn. (17) that *α*(*e*) = 0 for many edges *e*, we catch these cases in the outer sum of Eqn. (18) to quickly skip past them without evaluating the inner sum. For any edge *e* where *α*(*e*) ≠ 0 we need to generate a random set of trees for the inner sum by removing edges. One efficiency gain comes from recognising that it is only necessary to randomly remove edges descended from *e* because no other edges can affect the outcome of EvoHeritage generated on *e*. Furthermore, once an edge has been disconnected, we need not consider whether any of its descendent edges are disconnected because they will not inherit the EvoHeritage from *e* in any case.

We repeated the stochastic process and took the mean of those values, expressing the final result in standardised units. The number of repeats was 10^4^ except for the Quaternary period where it was 100, and the case where *ρ* = 0, which yields *β*(*e*) = 1 and removes all stochasticity from the result. All calculations were repeated for 100 mammal trees from the posterior distribution of the trees provided by Upham et al. (2019). In all cases we extended the stem of the tree back to the origin of life 4.025 billion years ago. We then took the median of our 100 values to calculate the living-fossil-ness for each vertex because the distributions are often skewed. For comparative purposes we calculated the ED for all species in the same 100 trees. We plotted the distribution of ranked living-fossil-ness across all mammals and showed box and whisker plots for the top 16 species, showing uncertainty from the 100 posterior trees. We produced a pairwise comparison of the living-fossil-ness ranks for all species between the Jurassic, Cretaceous and Quaternary periods for both *ρ* = 0 and *ρ* = 0.01.

## Empirical applications –Results

### i)Evaluating φ_ρ_ as a predictor of community productivity

The correlation coefficients of PD and species richness coincide exactly with those of *φ*_*ρ*_ at each of its limits, as expected. Considering the unfiltered community data first, the strongest correlation of plot biomass against plot *φ*_*ρ*_ for all species was achieved for the smallest *ρ*, where *φ*_*ρ*_ converges to PD. Considering only the species richness of the Pentapetalae, ignoring the contribution of any of the Poaceae, achieves a better correlation with biomass than any *φ*_*ρ*_ calculated for all species together. The PD of Pentapetalae correlated better with total biomass than the species richness of Pentapetalae. The strongest correlation with biomass was achieved with *ρ* ≈ 0.016, calculated for Pentapetalae only. The diversity of Poaceae was the poorest predictor of biomass, regardless of how it was measured, but unlike the other cases, it performed better as species richness (Poaceae) than it did as PD(Poaceae). For Poaceae, *φ*_*ρ*_ was optimised when *ρ* = 0.04 where it performed better than either species richness or PD (see Fig. 8).

**Fig. 8:**
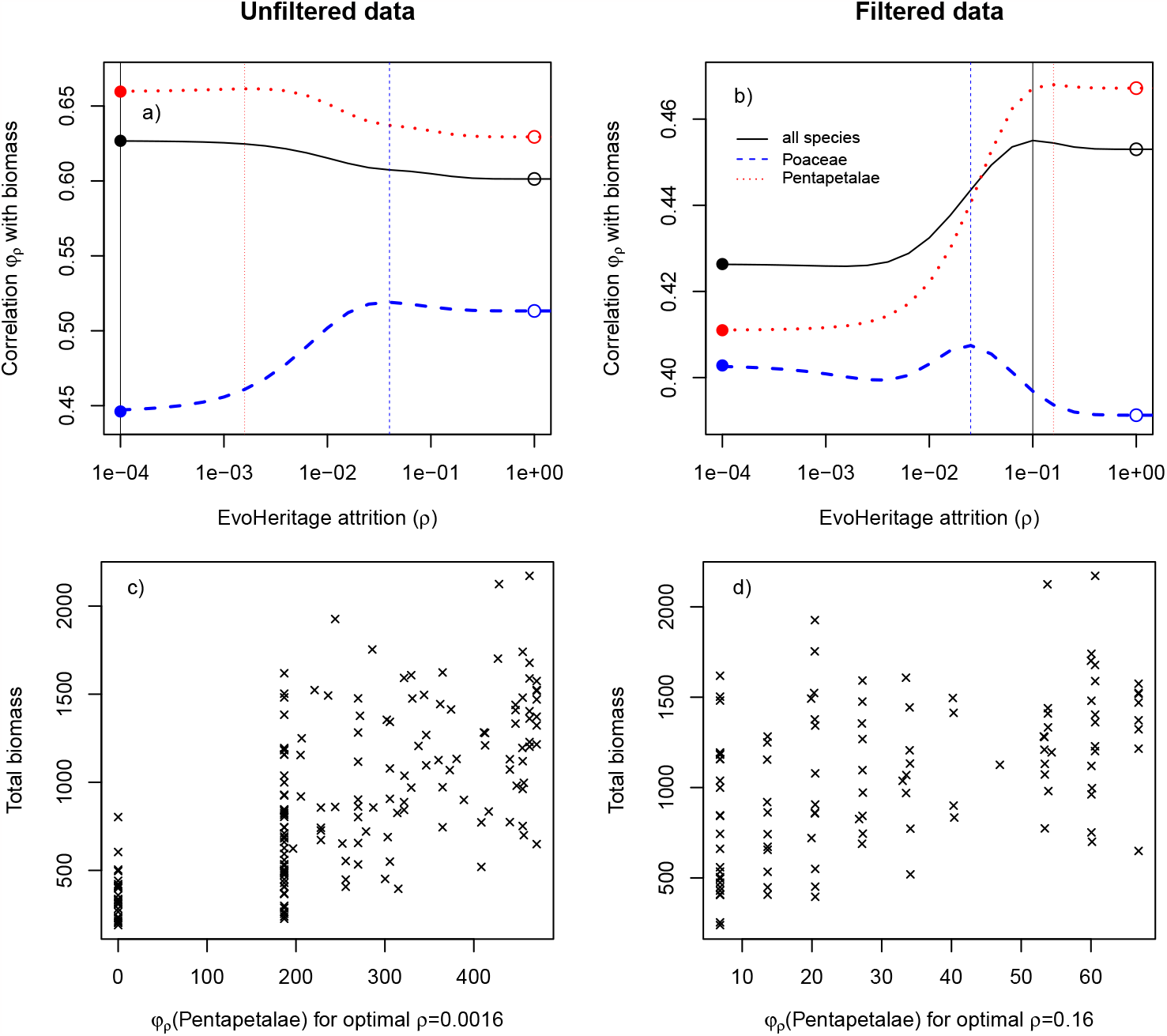
Relationship between EvoHeritage *φ*_*ρ*_ and community productivity. Panels a) and b) show Pearson’s correlation coefficient between *φ*_*ρ*_ and total biomass in the Cedar Creek plots. The horizontal axis shows attrition *ρ*. The solid circles give the same correlation, only with PD (corresponding to *φ*_*ρ*_ with small *ρ*) and the open circles give the correlation with species richness (corresponding to *φ*_*ρ*_ with large *ρ*). Vertical dotted lines show the optimal value of *ρ* that maximises the correlation coefficient. Data plotted in black corresponds to biodiversity measurement of all species present, blue corresponds to only Poaceae and red corresponds to only Pentapetalae. Panel a) shows results for all communities in the dataset ‘unfiltered data’. Panel b) shows results excluding communities that lack any species from either Pentapetalae or Poaceae. We do not show our data for *ρ >* 1 because it is clear *φ*_1_ already corresponds to species richness. Panels c) and d) show scatterplots of biomass against *φ*_*ρ*_ for the value of *ρ* and scenario in the panel directly above that gave the strongest correlation. This was always for measurement of Pentapetalae only because this was what maximised the correlation coefficient.

When community data was filtered to exclude communities that lack either Pentapetalae or Poaceae the correlation coefficient was poorer under almost all scenarios. The relative importance of species richness and PD switched to the exact opposite of that found for the unfiltered data. With filtering, species richness of all species and of Pentapetalae is superior to the corresponding measurement of PD whilst PD is superior to species richness when looking at Poaceae. The optimal values of *ρ* were 0.0025 for Poaceae, 0.1 for all species and 0.16 for Pentapetalae. All these values of *ρ* are far from both the species richness and PD ends of the spectrum (see Fig. 8).

### ii)Applying EvoHeritage attrition and partitioned-φ to quantify living fossils

Living-fossil-ness based on EvoHeritage from the Jurassic period yielded a stepped distribution of values with clear gaps between groups. The top species was *Ornithorhynchus anatinus*, the Duck-billed platypus. Next came all four species of echidna (Tachyglossidae) without much distinction between the four species. Then all seven species of shrew-opossum (Caenolestidae), followed by *Dromiciops gliroides* (known as the monito del monte) and then various other opossums. Overall, the top 16 living fossils from the Jurassic period are all Monotremes or South American marsupials. The rankings did not appear to be sensitive to the value of *ρ*, though the precise Living-fossil-ness values were dependent on *ρ* (Fig. 9). Living-fossil-ness from the Cretaceous yielded a smoother distribution of values, especially for *ρ* = 10^−2^. There are several noteworthy new entries to the top 16. The aardvark *Orycteropus afer* and the three species of Solenodon. Uncertainty in the placement of *Orycteropus afer*, which has a long terminal edge, leaves open the outside possibility for it to have much lower (or much higher) living-fossilness, but the median remains high.

**Fig. 9:**
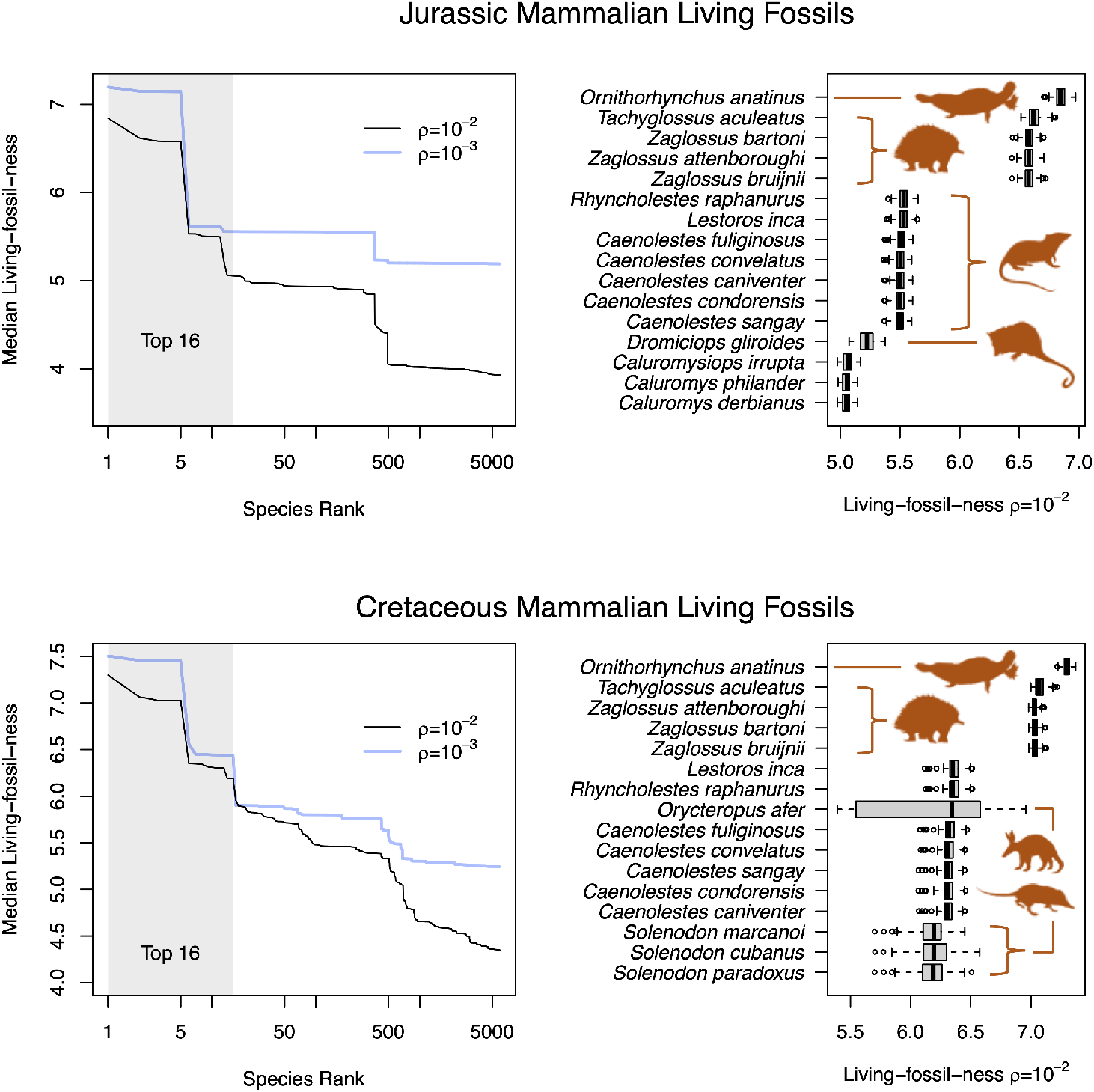
Living-fossil-ness for mammal species. Panels a) and b) are calculated for the Jurassic period, partitioning all EvoHeritage arising before 145 Million years ago. Panels c) and d) are for the Cretaceous (66 Million years ago and before). Jurassic living fossils may contain EvoHeritage arising before the Jurassic period, but surviving attrition to exist in species that were fossilised during this period. The same applies to Cretaceous living fossils. Panels and c) show the living-fossil-ness with rank in log scale for two different values of attrition: *ρ* = 10^−2^ and *ρ* = 10^−3^. The top 16 species are shaded. The box and whisker plots in panels and d) focus on *ρ* = 10^−2^ and top 16 species only. The uncertainty represented in the box and whisker plots is from 100 posterior trees. Each tree gives rise to one value in the distribution shown; each value is the result of 10^4^ random draws of attrition via a Monte Carlo approach. Image credit: Caenolestes (representing shrew opossums) and Dromiciops (monito del monte) Sarah Werning CC BY 3.0 both images recoloured in brown; other outline images are from the public domain.

Living-fossil-ness ranks of species based on ED are closely correlated with those based on paritioned-EvoHeritage (*ρ* = 0.01) for the Quaternary period. There is, however, a very substantive quantitative difference between partitioned-EvoHeritage and its ED counterpart partitioning only the PD corresponding to the Cretaceous or Jurassic periods (see supplementary material).

## Discussion

### i)Evaluating φ_ρ_ as a predictor of community productivity

Our results showed, as expected from the theory, that *φ*_*ρ*_ gives the same result as PD and species richness at its two extremes (see Fig. 8 and Thm. 1). This approach changes the nature of debates in this field from a stark choice between species richness and PD (Mazel et al., 2018, 2019; Owen et al., 2019) to a question about the extent of attrition captured by *ρ*. Previous work has suggested a metric that will converge to species richness and PD in different limits, based on pruning parts of the tree older than a given cutoff (Chao et al., 2010) in order to make values comparable across trees with different stem ages. This previous work may now be reinterpreted as an implicit form of attrition cutting away the deeper parts of the tree. EvoHeritage remains distinct however, because its approach is mechanistically led by the concept of the biological process of attrition and the need to account for it. Attrition exists as a biological reality, occurring at a rate that is neither zero (PD) nor infinity (species richness); it should be somewhere in between. The incorporation of phylogeny into ecology has, in the past, been criticised on the grounds that phylogeny’s role is relegated to a proxy for missing functional trait data (Swenson, 2013). By fitting the *ρ* parameter to a phylogeny, our approach allows us to better quantify how history predicts ecological process, and thus enabling eco-phylogenetics to give mechanistic insight into two disciplines (Pearse et al., 2019; Davies, 2021). Phylogenetic diversity is more than a proxy for function (Faith, 1992), it also captures other aspects of biodiversity including non-functional features such as future options for humanity and aesthetic value (Faith, 1992; Mooers et al., 2005). We claim, based on the conceptual arguments in this work, that the same applies with even greater weight to EvoHeritage.

We found that *φ*_*ρ*_ of all species present (that is the unfiltered community data), for any value of *ρ*, could not improve on PD as a predictor of community productivity. This result is not a failure of *φ*_*ρ*_ but is rather an example of how it can be informative. Specifically, it suggests that the deepest edges of the tree are the most important predictors of productivity, because attrition, which can only reduce the influence of such edges, is optimal when set to zero. The phylogeny of the species we considered has a deep split between the true grasses (Poaceae) and the other species, all of which are Pentapetalae (Fig. 6). PD of all species present in a plot will therefore primarily reflect the significant distinction between these two groups. This was confirmed by our finding that the species richness of Pentapetalae alone performed better as a predictor of biomass than any measure of total diversity. These results are probably the result of ecological factors such as species–species interactions and the special role of grasses in community productivity.

The best predictor of community productivity overall was the *φ*_*ρ*_ of Pentapetalae with an optimal *ρ* of 0.016 (see Fig. 8). This outperformed both species richness and PD and thus demonstrates the potential of *φ*_*ρ*_ as a general diversity measure. Although we found that the diversity of Poaceae is a poor predictor of community productivity, we highlight it as one example where species richness outperforms PD, though *φ*_*ρ*_ still outperforms both species richness and PD for a small range of values centered around *ρ* = 0.04 (Fig. 8).

Our tests of the filtered community data exclude any role of presence and absence at the level of Pentapetalae and Poaceae because any community with only one of the groups present was excluded. These filtered community results reinforced the conclusion that neither species richness nor PD extremes are optimal, as was the case for all three tests (measuring diversity of only Pentapetalae, only Poaceae, and both together). Species richness tends to prevail over PD in these filtered data, suggesting that the performance of PD was tied to its ability to pick up on key innovations for productivity occurring along the long edges separating Pentapetalae and Poaceae. In some sense this shows that PD is working as a paradigm to capture key innovations that may be unlikely to be reversed. We argue, however, that the majority of elements of biodiversity that are measurable from the tree of life are likely reversible evolutionary sources of variation. The prevalence of intermediate optimal values of *ρ* in our results, as well as the performance of species richness over PD in half of the cases, show that EvoHeritage has the flexibility to capture a wider range of evolutionary sources of variation than PD.

The differences in correlation coefficient are quite small across all our scenarios, which is consistent with the mixed results that comparisons between species richness and PD have so far yielded (Mazel et al., 2018). Furthermore, the best measure of overall diversity would not be one that completely ignores the presence of Poaceae as a component of diversity, yet this is what we find here if the goal is solely to predict community productivity. Our findings thus highlight the reasons why the ‘best’ measure of biodiversity in the most general sense of the word is not the same as the best predictor of community productivity in particular. It can thus be misleading to compare PD with species richness, as a measurement of biodiversity, on the basis of a single performance factor. Some debates around the role of PD have pointed to the conceptual foundations of PD in its defence (Owen et al., 2019). Our foundation for EvoHeritage as an alternative to PD is likewise based more on the conceptual reasons supporting its use, rather than its encouraging performance in this particular case.

### ii)Applying EvoHeritage attrition and partitioned-φ to quantify living fossils

Many of the top mammal species that we identified have a history of being referred to as living fossils in the primary literature, including the monotremes (Phillips et al., 2009), the Aardvark (Gózdziewska-Har-lajczuk et al., 2018) and solenodons (Wezel and Bender, 2002). Other than *Dromiciops gliroides* (Bozinovic et al., 2004), we only found a mention of South American marsupisals as living fossils in the grey literature. Our measure of living-fossil-ness therefore demonstrates that the taxa historically identified as living fossils can be recovered from our EvoHeritage-based concept of living fossils and requires no heterogeneity in the rate of evolution (Fig. 9). The key to our proposed solution of the living fossil controversy comes from recognising that living fossils show ancestral features that are rare in extant species, but need not show more ancestral features in total. Indeed, it has been pointed out before that scarcity is an important characteristic, otherwise we would have to admit bacteria as living fossils (Werth and Shear, 2014). Given that the identity of the living fossils themselves was originally established by a non-quantitative thought process, it is easy to imagine how the discovery of *rare* ancestral features in an extant species would carry greater weight and be more worthy of commentary. In contrast, the discovery of ancestral features that are commonly seen in extant species could easily be taken for granted. It also seems likely that rare ancestral features in an extant species would be more informative for reconstructing the features of extinct ancestors. Therefore, the phenomenon of living fossils as real and noteworthy objects of study might have been masked by the subtlety of rarity and a shortage of quantitative approaches. Our explanation still requires an imbalanced phylogenetic tree and does not seek to provide an explanation for why such imbalance occurs. It may be due to lower speciation or elevated extinction within a clade - if the latter it would make our living fossils ‘relict species’ in the spirit of Grandcolas et al. (2014). Perhaps in reality imbalance is caused by a mixture of factors. Regardless of the reason behind any phylogenetic imbalance, we have shown that the concept of living fossils does not need to invoke the ‘ladder of progress’ or be in conflict with mainstream thinking.

To the best of our knowledge, our method of using partitioned-*φ* on targeted EvoHeritage graphs is the only attempt focused on quantifying living fossils, other than the work of Bennett et al. (2018). Although Bennett et al. (2018) provided inspiration for our quantification procedure, our approach is different in several important respects. First, we calculated the ‘living-fossil-ness’ for individual vertices of a dated EvoHeritage tree, whereas Bennett et al. (2018) primarily designated clades of species as living fossils. Second, we were able to calculate the ‘living-fossil-ness’ differently depending on which period in geological history the fossils of interest came from, thus revealing a difference between the living fossil species that best represent rare ancient features present in fossils from different periods. Third, our calculations are process-based, incorporating the concept of attrition.

We note that ED (fair proportion) (Redding, 2003; Redding and Mooers, 2006; Isaac et al., 2007) has also been suggested as a metric of living fossils, though this was not its primary purpose (Bennett et al., 2018). Using ED to measure living-fossil-ness is a special case of our method corresponding to the scenario having zero attrition (*ρ* = 0) and including feature accumulation from the very recent past (see supplementary material). Besides the lack of attrition, this special case does not, in our view, correspond to the commonly intended meaning of a ‘living fossil’ as a species with a special similarity to an *ancient* fossil; not a special similarity to the remains of a recently departed species, which would be unremarkable and expected. We found that under both the ED and partitioned-*φ* paradigms there was a big difference between rankings of Quaternary living fossils, and living fossils from the older periods (Jurassic or Cretaceous - see supplementary material). This is likely because much of topology of the mammal tree has grown since the Cretaceous. The partitioned-*φ* case with *ρ* = 0.01 treats these recent parts of the tree very differently from the older parts because only attrition, and not accumulation, is permitted to occur. The ED case (*ρ* = 0) creates an even starker divide because it prevents the more recent parts of the tree from having any influence on the results at all. The effect of *ρ* is relatively weak in the Quaternary case. In contrast, the effect is strong in the Cretaceous case and stronger still in the Jurassic case (see supplementary material). This is because as we move from the Quaternary to the Cretaceous and Jurassic cases we are increasing the portion of the tree within which the biggest influence of attrition can be seen.

To bring the ED and partitioned-*φ* cases into even starker relief, let us contemplate Triassic living fossils from the mammals. The end of the Triassic occurs before the crown of the phylogeny of living mammals. According to the ED paradigm, we fairly partition the stem between all extant mammals and there is nothing to choose between them. According to partitioned-*φ*, however, all the younger edges of the tree have a role in determining the pattern of attrition and thus in determining the distribution of stem features among living species. For the same reason, the Jurassic period living fossils based on ED (*ρ* = 0) produces a distribution of only three possible values whilst *ρ* = 0.01 provides a way to distinguish between all the species that have equal living-fossilness ranks under ED (see supplementary material). Using our living-fossil-ness measure, older edges naturally emerge as the ones on which the EvoHeritage seen in fossils has accumulated. Younger edges in contrast give us the pattern of EvoHeritage attrition, and describe how ancient EvoHeritage is shared unevenly between tips as an emergent result.

Future technical work may improve the Monte Carlo algorithms by reweighing the probabilities of attrition along each edge to explore the parameter space more easily. This step may be necessary for tractable calculations in cases where *ρ* is large or extremely small. Here we have partitioned EvoHeritage fairly among extant vertices, but equally weighted all EvoHeritage from a given period of interest, regardless of how common this EvoHeritage would have been in fossils. It would be interesting to consider how this may change if we weight EvoHeritage according to how common it was in the fossils themselves, taking into account preservation bias and the effects of some fossils having come from now entirely extinct lineages. Future work may also seek to more formally test whether species labelled as living fossils match our findings, or seek to find *ρ* that optimises the outcome of such a test.

#### Implications for conservation applications in future work

Applications of PD to conservation focus on the idea that PD, usually calculated on a dated phylogenetic tree, captures something different and more valuable to protect than species richness, for example future options for humanity (Faith, 1992). The PD calculus may thus be used in conservation prioritisation protocols (Isaac et al., 2007; Steel et al., 2007; Gumbs et al., 2023), and metrics of success (Gumbs et al., 2021). It is outside the scope of this work to trial the natural counterparts of PD-focused conservation in terms of EvoHeritage, but we discuss the conceptual reasons why it may be beneficial to apply EvoHeritage to conservation.

Let us consider the lengths of the terminal edges that connect each terminal vertex (species) to the rest of the tree. PD incorporates (as a minimum) the length of the terminal edge for any species under consideration, because that edge is required in a connection to any other place on the tree. It is clear that species with long terminal edge lengths contribute more unique features to biodiversity than others. But how much more? The tuatara (*Sphenodon punctatus*), an extraordinary reptile endemic to New Zealand, has a terminal edge length of 275 million years and many unique features. It seems plausible to us that it possesses more unique features than, say, the duck-billed platypus (*Ornithorhynchus anatinus*), with a terminal edge length of about 64 million years, but having over four times as many features, as PD would assert, seems disproportionate. This issue is mirrored by an idea in population genetics that net differences between groups will become saturated, given sufficient time, rather than grow indefinitely, even if evolution remains rapid (Crozier et al., 2005). Saturation was addressed at the stage of tree construction for Crozier’s genetic diversity measure, where the edge lengths are bounded between 0 and 1, capturing the probabilities of a genetic difference (Crozier, 1992) meaning that edge lengths interacted in a multiplicative fashion, rather than additive as PD would assert. Framing in terms of EvoHeritage and attrition produces a similar effect because older features have more opportunities to be lost.

For terminal edges, these effects could be addressed within PD by re-scaling edge lengths appropriately (though this is not done in practice on dated trees). Indeed, edge lengths measured in years may not be the best choice for PD (Faith, 1994a) or other downstream analyses (Letten and Cornwell, 2015). Whilst our work gives a new way to re-scale edges in the form of *α* values, we also show that, for interior edges the effects of attrition are more complex and cannot be entirely captured by rescaling. This is because features may be gained on one edge but lost on a different edge, resulting in a non-monophyletic pattern of features (Wicke et al., 2021). More extreme examples than the tuatara with long interior edges are not hard to find; the Filasterea clade of just three species is sister to animals and choanoflagellates together (Shalchian-Tabrizi et al., 2008) and over 900 million years old (Ferrer-Bonet and Ruiz-Trillo, 2017; OneZoom Core Team, 2021) incorporating interior edges of well over 100 million years. The PD framework, especially on dated trees, seems incapable of assigning a reasonable value to such edges and their descendent species. Reconstructing such deep parts of the phylogeny is notoriously difficult, and it could be that the tree will rearrange around, say the Filasterea clade, in some future work, radically changing the ED scores of its species in the process. A weakness of the PD calculus is that on big trees it remains very sensitive to rearrangements of the deep and unstable parts of the tree of life. The same is not true of EvoHeritage because the deeper parts of the tree are subject to more attrition and thus carry less influence for comparing extant vertices. Indeed, the difficulty of reconstructing ancient parts of the tree of life is perhaps prominent evidence that attrition is a real force that cannot be ignored in phylogenetic analyses at such scales.

Drilling into the still deeper past, the PD of a group of species, based on the minimum spanning tree between those species and the root of the tree explicitly excludes the origin of life (Faith and Baker, 2006). It is arbitrary which point in the tree of life we choose to define the scope of a study, so excluding the origin of life makes PD comparable only up to the root of the clade being analysed and no further. In practice, this means results are not directly comparable between, for example, a study of mammals and a study of birds, unless both studies root their tree on the crown of Amniotes. Furthermore, it seems inconsistent to refer to PD as Evolutionary History if it in fact excludes an arbitrarily large portion of ancient evolutionary history from its total. We note that the concept of *unique* PD has none of these problems because the path to the origin of life is never unique (unless we’re measuring all life). We suppose that the real reason why PD has been set to exclude the origin of life is that to include it would lead to apparent absurdity with many studied groups having values only negligibly different to four billion years. In our view, this is treating the symptom (disproportionate quantitative values), but not the cause, which is failure to account for attrition. EvoHeritage resolves these issues because deeper parts of the tree, right back to the origin of life, can still add contemporary features, but have less weight than more recent parts of the tree (see Fig. 2).

More recent work into use of PD for practical conservation prioritisation has used mappings of IUCN Red List status onto the probability of extinction (Mooers et al., 2008) to optimise future PD with a metric known as HEDGE (Steel et al., 2007) and an associated protocol for using HEDGE, known as EDGE2 (Gumbs et al., 2023). One notable consequence of HEDGE is that the deeper edges of the tree have reduced influence because there is a small probability of all their descendants becoming extinct. The same mechanism attaches a value to there being more descendants of a deep edge because edges with more descendants are more likely to survive. HEDGE therefore mitigates some of the issues with PD that we have highlighted here, albeit as a byproduct of making the probability of extinction explicit. However, HEDGE is still based on PD, and one should not confuse the different underlying mechanisms and aims of HEDGE and EvoHeritage.

We need a coherent system for attributing value (EvoHeritage accounting for attrition) as well as for assessing the risk of losing parts of that value (HEDGE); neither should be ignored and future work may amalgamate both.

Work evaluating the rate of contemporary species extinction often compares this with background rates of extinction (and speciation) to establish that the present biodiversity crisis goes substantially beyond that expected from natural processes (De Vos et al., 2015). Studies focused on PD have concluded that it will take millions of years for mammal diversity to recover given current rates of diversity generation (Davis et al., 2018). PD generation, however, comes from two sources, extension of the terminal edges (tips), and speciation. The former has been noted to be sufficient to compensate for at most three species per year across all life (Purvis and Hector, 2000b). This is a small number compared to contemporary rates of biodiversity loss, however in the context of our work it may be smaller still because attrition adds an additional natural source of loss beyond background extinction, one that has not been previously accounted for. The inevitable growth in all terminal edges has two different effects: firstly, it adds new EvoHeritage just as happens with PD; secondly, it takes away old EvoHeritage because there are new opportunities for attrition to occur. Growth at the terminal edges of the tree of life over the natural passage of time does not therefore add much to net EvoHeritage because it has negative effects (attrition) that at least partially cancel the positive effects (accumulation). Future work may thus compare the background rate of EvoHeritage loss (through attrition and natural extinction together), gain (through accumulation and speciation together) and anthropogenic loss (through anthropogenic extinctions and loss of populations with distinct features).

#### The future of EvoHeritage trees and graphs

The foundations for EvoHeritage are based on the new concept of an EvoHeritage tree: a tree where the processes of both EvoHeritage accumulation and attrition are defined along the edges. We have provided standard conditions and associated calculations as a way to obtain an EvoHeritage tree from a classic tree with only edge lengths together with a value (or range of values) for the relative rate of attrition (*ρ*). It would be interesting to build EvoHeritage trees directly from genetic data by adapting tree-building methods and we expect this will be possible in future work. The key step in doing so would be to replace edge length with attrition (*β*) and accumulation (*α*) separately for each edge and allow them to be decoupled. We have shown through Theorem 2 that EvoHeritage accumulation and loss interact in a way that leads to non-monophyletic features and cannot be expressed with just a single dimension of data, such as an edge length. At least two dimensions are needed: one for EvoHeritage accumulation and the other for attrition. We would likely wish to retain geological dates as well in EvoHeritage trees to inform some kinds of analyses. Inference procedures for building dated trees seek to recover the passage of geological time along each edge. These methods account for mutation rate heterogeneity across nucelotides and the possibility of overwriting previous changes in parts of the genome with high mutation rate. This is the kind of information that could be used to separately inform *α* and *β* on each edge in future work, but is instead distilled to a single edge length. The fundamental units of EvoHeritage should probably not be nucelotides, but they may be genes or larger sections of genome. Alternatively an EvoHeritage graph may be constructed as the best fit to a matrix capturing the similarities among taxa. In any case, the concept of separating gain and loss will still apply.

To build an EvoHeritage tree instead of a phylogenetic tree would be a break in convention requiring a new data structure for storage and a new suite of methods. The potential reward might be an object with less compromise and uncertainty than a dated phylogenetic tree, which would enable more of the source data to inform downstream analyses as well as new questions. For example, how does the rate of attrition vary with body size? Did it change at important times of evolutionary pressure such as during past mass extinction events? Beyond this, future work may extend EvoHeritage trees to EvoHeritage graphs: directed acyclic graphs with attrition and accumulation specified on each edge, an extension of the concept of phylogenetic networks. These would be yet more general objects. For example, capturing hybridisation with *β* ≈ 0.5 because roughly equal EvoHeritage is inherited from each parent.

In order to facilitate practical applications of the EvoHeritage calculus, we developed a model of exponential EvoHeritage attrition that can be applied to trees with edge lengths, given *ρ*, a rate of attrition. This has the useful property that many existing measures are recovered when we set *ρ* = 0. The metric *φ*_*ρ*_ in particular was necessary to provide something that could directly stand in for PD in future analyses. We can see several approaches to determining the most appropriate *ρ* in such studies. The approach we used here was to experiment with a range of values and make the effect of *ρ* a subject of study rather than an inconvenience. This can be a strength, such as in our application to community productivity.

We see that in some cases, a fixed value of *ρ* will need to be decided upon. One approach is to use the value of *ρ* that optimises some payoff, such as a correlation with feature diversity. Another approach would be to use a rule of thumb. This may not seem ideal but if the alternative is to go to one extreme or the other (that is, to species richness or PD), then a rule of thumb may be better than nothing. If we expect parts of the tree more recent than a few tens of millions of years to still behave like PD, and edges are measured in millions of years, then *ρ* ≪ 0.1 would reflect these assumptions. Similarly, if we expect to see significant attrition over a billion years, then *ρ* ≫ 0.001 would be the approximate choice. Indeed, after investigating a wide range of *ρ* in our empirical study for living fossil detection, the interesting and intuitively correct results seem to be around *ρ* ≈ 0.01. The optimal values of *ρ* from our community productivity experiments ranged between 0.0025 and 0.16. The best correlation from any scenario was achieved from the EvoHeritage of Pentapetalae with *ρ* = 0.016. These results fit with the intuition that *ρ* ≈ 0.01 though a lot more research is needed with complementary tests for optimal *ρ* before a clear picture could begin to emerge.

#### Conclusion

In this manuscript, we began by pointing out a range of conceptual challenges with PD. First, following PD, a single extant descendant of a deep edge is enough to guarantee the survival of all ancestral features on that edge. This does not seem biologically reasonable; for example, it would mean one species of flightless bird alone could represent all features of the stem edge of birds. Second, the PD of a set of vertices is not increased by resurrecting an extinct common ancestor. For example, PD ignores any value that a direct ancestor such as the Archaeopteryx (or its close relative as the case may be) could add beyond that seen in living birds and crocodilians. Third, the features (including future options for humanity) cannot reasonably be directly proportionate to the edge length for extremely long edges. Whilst the last of the these challenges can be addressed for terminal edges by simply re-scaling longer edge lengths, the other challenges cannot be fully addressed by any kind of edge re-scaling. All the challenges are symptoms of the fact that EvoHeritage accumulates along edges but can also subsequently be lost; this leads to non-monophyletic patterns of feature distribution among vertices that could never be captured by PD, regardless of how edge lengths are set. A key realisation is that the same concerns about the existence of attrition are likely to apply to any other metric that tries to capture features from a tree, and is underpinned by the sums or means of inter-vertex distances in some way.

We introduced a new EvoHeritage calculus, which incorporates the process of ‘attrition’ capturing a gradual loss of features without extinction. Our mathematical results show that the independent forces of accumulation and attrition cannot be captured with the mainstream concept of a single value for ‘length’ of each edge. We have provided standard conditions as a way to proceed with calculations using phylogenetic trees and accounting for accumulation and attrition, until EvoHeritage trees (or graphs) can be produced directly by other methods. We have proved that on a tree, both the value of *φ*_*ρ*_ (total EvoHeritage under standard conditions with attrition at rate *ρ*), and the sets of terminal vertices that maximise it, cannot be perfectly captured by PD regardless of how edge lengths are set. The difference was small, however, for our simple quantitative examples (see supplementary material). A fruitful direction for future work would be to find edge re-scaling that maximises the ability of PD to replicate *φ*_*ρ*_ and quantify any remaining error. We expect that provided the product of crown age and *ρ* remains small, the existing PD paradigm may continue to apply either as it stands, or with some edge re-scaling of the tree (see Fig. 2). However, for a larger-scale phylogeny this is unlikely to be the case. The situation is reminiscent of the curvature of earth which may be ignored at the scale of local maps, but demands attention as we progress to continental scales, requiring a new geometry that incorporates euclidean geometry as a special case. As we scale our field of vision to the complete tree of life, we believe it will not be feasible to ignore the force of EvoHeritage attrition.

We have introduced *φ*, unique-*φ*, partitioned-*φ*, expected-*φ*, and diff-*φ* to illustrate EvoHeritage counterparts to wide ranging concepts from the PD ‘calculus’ (Faith et al., 2004). There are many more possible counterparts. For example, on an EvoHeritage tree with extant vertex range maps the phylogenetic endemism concept of Rosauer et al. (2009) could become ‘EvoHeritage endemism’ by spreading the value of EvoHeritage equally across the area in space where that EvoHeritage exists. The same is true of more recent spatially explicit metrics of PD (Gumbs et al., 2020). Another example of important potential future work would be to incorporate abundances of individuals representing each terminal vertex and Hill numbers following (Chao et al., 2010). We also imagine a reinterpretation of tree balance in terms of EvoHeritage rather than of lineage numbers.

We introduced standard conditions and *φ*_*ρ*_ to make the adoption of EvoHeritage easier as a straight swap for PD on any tree with edge lengths, given a value of *ρ*. To use the EvoHeritage calculus under standard conditions (fixed rate of accumulation and attrition for EvoHeritage), one only needs a phylogenetic tree with edge lengths and a value of *ρ* (or a range of values). Our two applications of EvoHeritage both rely on its attrition component. EvoHeritage can outperform PD and species richness as a predictor of community productivity and metrics on a targeted EvoHeritage graph can detect living fossils and resolve their controversy. We have also discussed the potential for future work to apply EvoHeritage in the field of conservation.

We are aware that there is already an apparent excess of phylogenetic metrics, many of them redundant (Tucker et al., 2017). Indeed, it has been argued that this ‘jungle of indices’ impedes, rather than facilitates, scientific progress and uptake in conservation applications (Pausas and Verdú, 2010). We view the jungle in the different light of the process that we think created it. Researchers have qualitative questions that they wish to answer; doing so requires them to have metrics with certain properties. It is easy to create metrics with the required properties from scratch and so researchers do so. The ‘cost’ of accidentally duplicating, or reinventing a subtly different version of, something that already exists is very small for a metric compared with other research endeavours. The main message of this paper is thus not our individual metrics, but rather the conceptual importance of attrition and the flaws associated with just summing edge lengths (even if edge lengths are re-scaled first). We urge researchers to either account for attrition in the metrics they create, or stick to questions where the symptoms of attrition are unlikely to manifest, such as on trees with relatively recent crown ages, or where the quantity of interest is extrinsically justified as the units of edge length. One should thus use PD for a question explicitly about evolutionary history, the number of years things have been evolving for. However, it is hard to ignore attrition if we are making value judgements on the basis of something that accumulates, such as features, including future options for humanity. To capture these things we need EvoHeritage and attrition, whilst this will increase complexity, it will also make the resulting metrics more coherent. Furthermore, alternative and simpler metrics of biodiversity may incur a greater danger that strategies for maximising them lead to loss of variety in some form that is not incorporated in the metric. If we may summarise the original message of PD as ‘species are not equal, their value depends on the edge lengths that represent their evolutionary histories’, our message is ‘unit lengths of edge are not equal, their value depends on context and the evolution of their descendants’.

## Supporting information

Supplementary Glossary

## Acknowledgements

We thank Arne Mooers, an anonymous reviewer and the editorial team for their really useful comments that have helped improve this work. We thank (in alphabetic order) Ryan Chisholm, Alexei Drummond, Dan Faith, Lucas Dias-Fernandes, Daniel Huson, Robert Noble, Nisha Owen, Roseli Pellens, Sebastian Pipins, Laura Pollock, Charles Semple, Bill Sherwin and Yan Wong for very useful discussions and suggestions. We also thank the Rosindell lab group members, and members of the IUCN Phylogenetic Diversity Task Force. Through JR and WDP, this study is an output of the Georgina Mace Centre for the Living Planet at Imperial College London. KM and MS thank the New Zealand Marsden Fund (MFPUOC2005). JR thanks the Leverhulme Trust for funding this research through a Research Fellowship (RF-2022-497).

## Author contributions

JR conceived of the initial idea of EvoHeritage, its limiting cases and basic results. JR and MS then developed the ideas further through meetings and exchange of notes. KM wrote and proved Thm. 2, developed results related to stochastic standard conditions, generated Fig. 3 and wrote large parts of the supplementary material and proofs. KM, MS and JR checked and clarified the mathematical results. WDP suggested the application to Cedar Creek and provided code for this analysis using PD; JR adapted this code to use Evolutionary Heritage. JR developed the application to living fossils, associated power set approach and Monte Carlo simulation with feedback from MS, KM and WDP. RG provided insights from an early stage into the conservation applications and phylogenetic diversity framing of the work. MS provided valuable guidance to KM and JR throughout. JR wrote the manuscript, produced the figures and graphs (excluding Fig. 3) and some parts of the supplementary material. All authors commented meaningfully on the manuscript and graphics at several stages prior to submission and during revision.

## supplementary material

Data and supplementary PDF files available from this Dryad Digital Repository link:

https://datadryad.org/stash/dataset/doi:10.5061/dryad.z08kprrgs

Code written for this work can be found on GitHub and is available for general use using the link:

https://github.com/jrosindell/EvoHeritage

