## Supplementary Material for "Phylogenetic Biodiversity Metrics Should Account for Both Accumulation and Attrition of Evolutionary Heritage"

September 4, 2023

James Rosindell[1,2,*], Kerry Manson[2], Rikki Gumbs[3],
Will Pearse[1] and Mike Steel[2]

[1] *Department of Life Sciences, Silwood Park Campus, Imperial College London, Buckhurst Road, Ascot, Berkshire, SL5 7PY, United Kingdom*
[2] *Biomathematics Research Centre, University of Canterbury, Christchurch, New Zealand*
[3] *EDGE of Existence Programme, Zoological Society of London, Regent's Park, London NW1 4RY, UK*

### 1 Introducing EvoHeritage ($\varphi$)

#### 1.1 Foundational results

##### 1.1.1 Generation of new vertices

We show that any edge can be split into two edges to create an additional vertex without affecting the results of EvoHeritage calculations on existing vertices. Consider edge $e \in E$ with the associated values $\alpha(e)$ and $\beta(e)$. Firstly $\alpha(e)$ gives the net amount of EvoHeritage accumulated along each edge $e \in E$. Secondly $\beta(e)$ gives the proportion of EvoHeritage present at the start of the edge $e$ that survives attrition along the entire edge to still be present at its end vertex $d^V(e)$.

We will describe a method to insert a new vertex $v^*$ along edge $e$, splitting the edge into two parts $e_1$ and $e_2$. Let $L(e)$ give the length of $e$. We describe the position of $v^*$ by the equations:

$$L(e_1) = L(e) \cdot (1 - x) \tag{1}$$

$$L(e_2) = L(e) \cdot x. \tag{2}$$

The EvoHeritage accumulated along both new edges and surviving attrition to arrive at $d^V(e)$ must remain the same as it was before before the new vertex was added. Therefore

$$\alpha(e_1) \cdot \beta(e_2) + \alpha(e_2) = \alpha(e). \tag{3}$$

In addition, the total EvoHeritage lost to attrition along both new edges must also remain unchanged and therefore

$$\beta(e_1) \cdot \beta(e_2) = \beta(e) \tag{4}$$

Provided that $\beta(e) \leq \beta(e_1) \leq 1$, $\beta(e) \leq \beta(e_2) \leq 1$, $0 \leq \alpha(e_2) \leq \alpha(e)$ and $0 \leq \alpha(e_1) \leq \frac{\alpha(e)}{\beta(e_2)}$ we can see numerous possible solutions for placement of the vertex that meet the condition that satisfy Eqns. 1 and 2. A general family of solutions is provided by taking a variable $0 \leq x \leq 1$ to define positioning of the new vertex. This leads to the natural solution below, which can be readily seen as having the required properties.

$$\beta(e_1) = \beta(e)^{1-x} \tag{5}$$

$$\beta(e_2) = \beta(e)^x \tag{6}$$

$$\alpha(e_2) = \alpha(e) \cdot x \tag{7}$$

$$\alpha(e_1) = \frac{\alpha(e) \cdot (1 - x)}{\beta(e)^x} \tag{8}$$

Under standard conditions, with accumulation parameter $\lambda$ and attrition parameter $\rho$, we can set the following values:

$$\alpha(e_1) = \frac{\lambda}{\rho}(1 - \exp(-\rho \cdot L(e_1))) \tag{9}$$

$$\alpha(e_2) = \frac{\lambda}{\rho}(1 - \exp(-\rho \cdot L(e_2))) \tag{10}$$

$$\beta(e_1) = \exp(-\rho \cdot L(e_1)) \tag{11}$$

$$\beta(e_2) = \exp(-\rho \cdot L(e_2)) \tag{12}$$

Now, let us verify that these solutions are consistent with Eqns. (3) and (4).

$$\begin{aligned}
&\alpha(e_1) \cdot \beta(e_2) + \alpha(e_2) \\
=&\frac{\lambda}{\rho}(1 - \exp(-\rho \cdot L(e_1))) \cdot (\exp(-\rho \cdot L(e_2))) + \frac{\lambda}{\rho}(1 - \exp(-\rho \cdot L(e_2))) \\
=&\frac{\lambda}{\rho}[\exp(-\rho \cdot L(e_2)) - \exp(-\rho \cdot (L(e_1) + L(e_2))) + 1 - \exp(-\rho \cdot L(e_2))] \\
=&\frac{\lambda}{\rho}[1 - \exp(-\rho \cdot L(e))] = \alpha(e)
\end{aligned} \tag{13}$$

$$\beta(e_1) \cdot \beta(e_2) = \exp(-\rho \cdot (L(e_1) + L(e_2))) = \exp(-\rho \cdot L(e)) = \beta(e) \tag{14}$$

Hence for a new vertex added anywhere along edge $e$, consistent new values of $\alpha$ and $\beta$ may be found for the newly created edges $e_1$ and $e_2$.

###### 1.1.2 Suppression of vertices of degree 2

As a counterpoint to the generation of new vertices above, we now show that we can suppress vertices of degree two without impacting the calculation of $\varphi$. The values for EvoHeritage accumulation and attrition on the resultant edge are constructed from the original values of the merged edges.

**Lemma 1.1** *Let $T = (V, E)$ be a rooted phylogenetic tree, and for some $n \geq 3$, let $P = v_1, v_2, \ldots, v_n$ be a path in $T$. Let $e_i = (v_i, v_{i+1})$ and suppose $v_2, \ldots, v_{n-1}$ are all vertices of degree 2. If $e^*$ is the edge that results from suppressing $v_2, \ldots, v_{n-1}$, then setting $\alpha(e^*) = \sum_{i=1}^{n-1} \alpha(e_i) \prod_{j=i+1}^{n-1} \beta(e_j)$, and $\beta(e^*) = \prod_{i=1}^{n-1} \beta(e_i)$ does not alter $\varphi$ values.*

###### 1.1.3 Proof of Lemma 1.1

We prove that one vertex of degree 2 can be suppressed. By repeated application of this process we can extend the result to the entire path $P$. Consider $v_1, v_2, v_3$, a subpath of $P$, and suppose that $v_1$ is, of the three vertices, the closest to the root of $T$. Let $R$ be the path from the root vertex to $v_1$, and let $T'$ be the subtree descending from $v_3$. If $X \subseteq V(T')$, then

$$\begin{aligned}\varphi(X) &= \sum_{e\in T'} \alpha(e)p(d^V(e),X) + \alpha(e_2)p(v_3,X) + \alpha(e_1)p(v_2,X) + \sum_{e\in R} \alpha(e)p(d^V(e),X) \\ &= \sum_{e\in T'} \alpha(e)p(d^V(e),X) + p(v_3,X)[\alpha(e_2) + \alpha(e_1)\beta(e_2) + \varphi(\{v_1\})\beta(e_1)\beta(e_2)].\end{aligned}$$

Where the term $p(v, X)$ is defined as in the main text: the proportion of EvoHeritage associated with the vertex $v$ that survives to be counted on at least one vertex in the set $X$. Hence if we set $\alpha(e^*) = \alpha(e_2) + \alpha(e_1)\beta(e_2)$ and $\beta(e^*) = \beta(e_1)\beta(e_2)$ we do not change $\varphi$(X) after the suppression of $v_2$.

□

#### 1.2 The independence of $\varphi_\rho$ from the choice of accumulation rate

Under standard conditions, EvoHeritage accumulates at a fixed deterministic rate of $\lambda$ units of EvoHeritage per unit of edge length. We wish to show that $\varphi_\rho$ does not depend on the choice of $\lambda$. Recall from the main text that

$$S_u = \begin{cases} \lambda & \text{when } \rho = 0; \\ \frac{\lambda}{\rho}\left(1 - e^{-\rho}\right) & \text{otherwise,} \end{cases} \tag{15}$$

and for edge $i$

$$\alpha(i) = \begin{cases} \lambda L(i) & \text{when } \rho = 0; \\ \frac{\lambda}{\rho}\left(1 - e^{-\rho L(i)}\right) & \text{otherwise.} \end{cases} \tag{16}$$

We now calculate $\varphi_\rho(X)$, for an arbitrary set of vertices $X$, for the two possibilities for $\rho$. When $\rho = 0$ we have

$$\begin{aligned}\varphi_0(X) &= \frac{\varphi(X)}{S_u} \\ &= \frac{\sum_{i\in E} \alpha(i) \cdot p(d^V(i),X)}{S_u} \\ &= \frac{\sum_{i\in E} \lambda L(e) \cdot p(d^V(i),X)}{\lambda} \\ &= \sum_{i\in E} L(e) \cdot p(d^V(i),X).\end{aligned} \tag{17}$$

When $\rho > 0$ this same calculation gives

$$\begin{aligned}\varphi_\rho(X) &= \frac{\varphi(X)}{S_u} \\ &= \frac{\sum_{i \in E} \alpha(i) \cdot p(d^V(i), X)}{S_u} \\ &= \frac{\sum_{i \in E} \frac{\lambda}{\rho}\left(1 - e^{-\rho L(i)}\right) \cdot p(d^V(i), X)}{\frac{\lambda}{\rho}\left(1 - e^{-\rho}\right)} \\ &= \frac{\sum_{i \in E}\left(1 - e^{-\rho L(i)}\right) \cdot p(d^V(i), X)}{\left(1 - e^{-\rho}\right)}.\end{aligned} \tag{18}$$

In both cases the final expression contains no $\lambda$ terms. Hence under standard conditions the value of $\varphi_\rho$ does not depend on our choice of $\lambda$. For this reason, we choose to set $\lambda = 1$ for the proofs in Section 1.4 of this supplementary material without any loss of generality.

#### 1.3 The role of the origin of life

##### 1.3.1 Why include the origin of life?

As discussed in the main text, it is beneficial to include the origin of life $\hat{o}$ as a vertex in our formulation. One reason is to make EvoHeritage of groups with different crown ages comparable. Without including the origin of life the total PD of a group with a shallow crown (or stem) age would be unfairly penalised when it comes to a comparison with a group that has a deeper crown (or stem) age.

Another motivation is to enable a consistent definition of the EvoHeritage (or PD) of a single vertex (or species). Let us begin by considering classic PD of a single species on a tree. The distinction between unique-PD and PD is important here. The unique-PD of a single species should clearly be that species' terminal edge (branch) length. However, the total (not necessarily unique) PD of a single species has often been taken to also mean the terminal branch length of that species, if it is defined at all. We argue that a more logically consistent answer for the total PD of a single species should be the date back to the origin of life vertex $\hat{o}$, approximately 4 billion years (Dodd et al., 2017). This is because any living species is the result of that number of years of evolutionary history (some of which is unique to that species and some of which is not).

Set against these reasons for incorporating the origin of life $\hat{o}$ as a vertex, is the uncertainty in the value of its date, represented hereafter as $\tau$. There may also be an intuition that the value of $\tau$ should not be relevant to measurement of contemporary diversity, as well as being a quantity that increases (albeit by a minute percentage of its total) with each further year that passes. So the value of $\tau$ measured now in 2023 will be 1 year shorter than the value measured in 2024. To resolve this we explore the sensitivity of our results to the value of $\tau$.

##### 1.3.2 Sensitivity to fluctuations in the origin of life

Consider the equation for a standardised unit based on the net EvoHeritage accumulation (the $\alpha$ value) of an edge with length $\tau$:

$$S_u = \frac{1 - \exp(-\rho \cdot \tau)}{\rho}. \tag{19}$$

We can also write this as

$$S_u = \frac{1}{\rho}\left(1 - \frac{1}{\exp(\rho \cdot \tau)}\right) \tag{20}$$

It is now clear, given that $\tau$ is certainly very large, that unless $\rho$ is extremely small, $S_u \approx \frac{1}{\rho}$ and is independent of $\tau$. We may also wish to constrain $\rho$ from being so small as to prevent current uncertainty in $\tau$ from having any noticeable effect. Take $\tau_{min} \leq \tau \leq \tau_{max}$ to express the uncertainty in $\tau$ and suppose that we wish our resulting value of $S_u$ to be accurate to within an error of $N$ significant figures. We therefore require

$$\frac{\frac{1}{\rho}\left(1 - \frac{1}{\exp(\rho \cdot \tau_{max})}\right) - \frac{1}{\rho}\left(1 - \frac{1}{\exp(\rho \cdot \tau_{min})}\right)}{\frac{1}{\rho}\left(1 - \frac{1}{\exp(\rho \cdot \tau_{min})}\right)} \leq 10^{-N} \tag{21}$$

Let us use the current best estimates of $\tau_{min} = 3.77 \times 10^9$ years and $\tau_{max} = 4.28 \times 10^9$ years following (Dodd et al., 2017). Let us take $N = 5$ for 5 significant figures, an accuracy greater than that used in many phylogenetic calculations. Eqn. (21) can now be solved numerically to give a value of $\rho$ sufficient to certainly remove undesirable dependency on the uncertainty around the date of the origin of life. The result is $\rho \geq 10^{-8.525}$, thus we probably should not consider values of $\rho$ below this value if we have opted to use the origin of life to define a standardized unit, and want to be certain that the origin of life uncertainty is not affecting our results.

#### 1.4 Relationship to species richness and PD

##### 1.4.1 Proof that $\varphi_\rho$ converges to a count of vertices (species richness) when $\rho \to \infty$

We seek to show that $\lim_{\rho \to \infty} \varphi_\rho(X) = \sum_{v \in X} |A^E(v)|$ where $X \subseteq V$. In words, the EvoHeritage of a set X, under standard conditions, converges to a count of terminal edges in X as attrition grows large. This can be seen as a convergence to species richness if terminal vertices represent species in a tree. Let us begin by considering the values of $\alpha(e)$ (net EvoHeritage accumulation along edge $e$), $\beta(e)$ (proportion of EvoHeritage lost along edge $e$) and $S_u$ (the size of a standardised unit) in this limit. For convenience, we repeat the definitions from the main text.

$$\alpha(e) = \frac{\lambda}{\rho}(1 - e^{-\rho \cdot L(e)}) \tag{22}$$

$$\beta(e) = e^{-\rho \cdot L(e)} \tag{23}$$

$$S_u = \begin{cases} \lambda, & \text{when } \rho = 0 \\ \frac{\lambda}{\rho}(1 - e^{-\rho}), & \text{otherwise} \end{cases} \tag{24}$$

Here we now have to assume that $L(e) > 0$. This is reasonable because one could collapse any zero branch lengths by merging the connected nodes, maybe resulting in a polytomy. Consequently $(\rho \cdot L(e)) \to \infty$ as $\rho \to \infty$, and so

$$\lim_{\rho\to\infty} \beta(e) = \lim_{\rho\to\infty} \exp(-\rho \cdot L(e)) = 0. \tag{25}$$

Let us now consider $S_u$.

$$\lim_{\rho\to\infty} S_u = \lim_{\rho\to\infty} \frac{\lambda}{\rho}(1 - e^{-\rho}) = 0 \tag{26}$$

However, the value of $\varphi_\rho$ will later be calculated as a fraction with $S_u$ on the denominator, so we cannot take this limit and should instead write $S_u$ as an approximation that will be valid in the limit of large $\rho$:

$$S_u = \frac{\lambda}{\rho}(1 - e^{-\rho}) = \frac{\lambda}{\rho} - \frac{\lambda}{\rho \cdot e^{\rho}} \approx \frac{\lambda}{\rho} \text{ if } \rho \gg 1 \tag{27}$$

Following the same logic and again assuming $L(e) > 0$ we can say that $\alpha(e)$ is independent of $L(e)$ and can be approximated by

$$\alpha(e) \approx \frac{\lambda}{\rho} \text{ if } \rho \cdot L(e) \gg 1. \tag{28}$$

Now let us apply these values of $\alpha$ and $\beta$ to give $\varphi$ under standard conditions. Recall that under standard conditions there is only one form of EvoHeritage, and thus taking equation 5 from the main text we can see that

$$p(v, X) = \begin{cases} 1, & \text{when } v \in X \\ 1 - \prod_{j \in D^E(v)}(1 - p(D^V(j), X) \cdot \beta(j)), & \text{otherwise.} \end{cases} \tag{29}$$

And so taking the limit for $\rho \to \infty$ we get $\beta(j) = 0$ and consequently that

$$p(v, X) = \begin{cases} 1, & \text{when } v \in X \\ 0, & \text{otherwise.} \end{cases} \tag{30}$$

Now recall equation 6 from the main text.

$$\varphi(X) = \sum_{e \in E} \alpha(e) \cdot p(D^V(e), X) \tag{31}$$

Taking the limit for $\rho \to \infty$ again we can write

$$\lim_{\rho\to\infty} \varphi(X) = \lim_{\rho\to\infty} \sum_{e\in E} \alpha(e)\cdot p(D^V(e), X) \approx \frac{\lambda}{\rho}\cdot\left|\{e \in E | D^V(e) \in X\}\right| = \frac{\lambda}{\rho}\cdot\sum_{v\in X} |A^E(v)| \tag{32}$$

Now the limit of $\varphi_\rho(X)$ as $\rho$ tends to $\infty$ can be given by:

$$\lim_{\rho\to\infty} \varphi_\rho(X) = \frac{\lim_{\rho\to\infty} \varphi(X)}{S_u} \approx \frac{\frac{\lambda}{\rho} \cdot \sum_{v\in X} |A^E(v)|}{\frac{\lambda}{\rho}} = \sum_{v\in X} |A^E(v)| \tag{33}$$

Which is a count of the ancestral edges of vertices in $X$. If we consider a tree there would be only exactly one edge leading towards each vertex and thus the result could equally be considered as a count of vertices (species richness if vertices represent species).

##### 1.4.2 Proof that $\varphi_\rho$ converges to PD when $\rho \to 0$

We next prove that $\lim_{\rho\to 0} \varphi_\rho(X) = PD(X)$. In words, $\varphi_\rho(X)$ converges to classic PD of the set of vertices when $\rho \to 0$. Let us begin, as in section 1.4.1, by considering the values of $\alpha(e)$ (net EvoHeritage accumulation along edge $e$), $\beta(e)$ (proportion of EvoHeritage lost along edge $e$) and $S_u$ (the size of a standardised unit) in this limit. Recall that

$$\alpha(e) = \frac{\lambda}{\rho}(1 - e^{-\rho\cdot L(e)}). \tag{34}$$

Taking the limit $\rho \to 0$ gives an indeterminate form of $\frac{0}{0}$. Applying L'Hôspital's Rule allows us to evaluate this limit:

$$\lim_{\rho\to 0} \alpha(e) = \lim_{\rho\to 0} \frac{\lambda L(e) e^{-\rho\cdot L(e)}}{1} = \lambda L(e). \tag{35}$$

Although we have already defined $S_u = \lambda$ at $\rho = 0$, in the limit as $\rho \to 0$ we can apply L'Hôspital's Rule in a similar manner to prove consistency

$$\lim_{\rho\to 0} S_u = \lim_{\rho\to 0} \frac{\lambda}{\rho}(1 - e^{-\rho}) = \lim_{\rho\to 0} \frac{\lambda e^{-\rho}}{1} = \lambda. \tag{36}$$

Additionally, note

$$\lim_{\rho\to 0} \beta(e) = \lim_{\rho\to 0} e^{-\rho\cdot L(e)} = 1. \tag{37}$$

This final limit simplifies the calculation of the $p(v, X)$ terms used in the calculation of $\varphi$. As $\rho$ tends to 0 we get:

$$p(v, X) = \begin{cases} 1, & \text{when } v \in X \\ 1 - \prod_{j\in D^E(v)} (1 - p(d^V(j), X)), & \text{otherwise,} \end{cases} \tag{38}$$

which means that

$$p(D^V(e), X) = \begin{cases} 1, & \text{when at least one descendant of } e \text{ is in set } X \\ 0, & \text{otherwise.} \end{cases} \quad (39)$$

Which is equivalent to saying $p(D^V(e), X) = 1$ if and only if $e$ is part of the minimum spanning set of edges for the vertices in $X$ including the root vertex. Hence

$$\lim_{\rho \to 0} \varphi(X) = \sum_{e \in E} \alpha(e) \cdot p(d^V(e), X) = \sum_{e \in E} \lambda L(e) \cdot p(d^V(e), X) = \lambda PD(X). \quad (40)$$

This last expression is the product of $\lambda$ and classic PD of the set of vertices including the root vertex. To complete the proof,

$$\lim_{\rho \to 0} \varphi_\rho(X) = \lim_{\rho \to 0} \frac{\varphi(X)}{S_u} = \frac{\lambda PD(X)}{\lambda} = PD(X) \quad (41)$$

##### 1.4.3 Conditions for $\varphi$ to correspond to PD

In this section we consider the question of whether the EvoHeritage ($\varphi$) measure may simply be replaced by PD, after an adjustment of edge lengths (where necessary) to absorb the consequences of attrition. We show that there is only a small family of phylogenetic trees for which this adjustment is always possible, whatever the pattern of accumulation and attrition on their edges. Furthermore, we show that such an adjustment is possible on trees outside this family only in very constrained circumstances, where attrition is isolated to a select set of edges near the root of the tree. This shows that $\varphi$ represents a true generalisation of PD.

Let $T = (V, E)$ be a rooted phylogenetic tree. We say that $\varphi$ *corresponds to PD* on $T$ if for every set of leaves $X$ in $T$ we have $\varphi(X) = PD(X)$. For each edge $e \in E(T)$, the EvoHeritage accumulation and attrition along $e$ are, respectively, denoted by $\alpha(e)$ and $\beta(e)$. To achieve greater generality, we allow the $\alpha$ and $\beta$ values of edges to be independent of each other (that is, not necessarily determined by a specific $\rho$ value across an entire tree). Recall that the equation $\beta(e) = 1$ describes the situation where no EvoHeritage attrition occurs along $e$, whereas the equation $\beta(e) = 0$ describes complete attrition occurring along edge $e$.

Let $L(e)$ denote the length of $e$. Because we allow $T$ to contain polytomies, we require that the length of every edge in $E(T)$ is strictly positive (including after any rescaling). We also make use of the terms parent and child vertex. If $v_1, v_2 \in V(T)$ share an edge, we call $v_1$ the *parent* of $v_2$ if $v_1$ is closer to the root of $T$. In this case we call $v_2$ a *child* of $v_1$. Additionally, we will use the term clade in the following specific sense. Let $e \in E(T)$ be an edge whose parent vertex is the root of $T$. The *clade of $T$ determined by $e$* is a subtree of $T$ comprised of the root vertex, $e$, and the entire subtree descending from $e$.

To clarify the notion of $\varphi$ corresponding to PD, we give an example of a tree for which a rescaling of edge lengths does indeed lead to $\varphi(X) = PD(X)$ for every set of leaves $X$. Consider the tree $T$ in Figure 1, and suppose that $\alpha(i) = 1$ for every edge $i$ in $E(T)$. Moreover suppose that $\beta(i) = 1$ for $i \in \{a, b, c, d\}$ and $\beta(i) = \frac{1}{3}$ for $i \in \{e, f, z\}$.

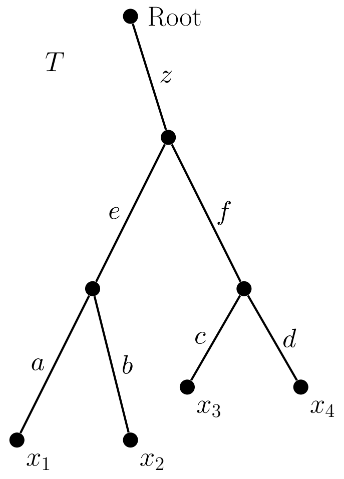

Figure 1: Suppose attrition is confined to edges $e, f$ and $z$ of the tree $T$ above. It is then possible to re-scale edge lengths such that the PD score of any set of terminal vertices on the rescaled tree matches the EvoHeritage value of that set. In contrast, if any attrition occurs on any of the four pendant edges of this tree, then no rescaling of edges is able to result in $\varphi$ corresponding to PD.

Then the following selection of edge lengths results in $\varphi$ corresponding to PD on $T$: $L(a) = L(b) = L(c) = L(d) = 1$ and $L(e) = L(f) = \frac{11}{9}$ and $L(z) = \frac{1}{9}$. Table 1 gives $\varphi(X)$ and $PD(X)$ values for selected sets of leaves $X$. In each case the desired equality holds. Values for other sets of leaves may be determined using the symmetry of $T$. Hence in this case $\varphi$ can be said to correspond to PD. We now proceed to show that such a correspondence is constructable only in very particular circumstances.

In light of Lemma 1.1 we consider phylogenetic trees with no monotomy vertices. After monotomy vertices are suppressed, the clades of a rooted phylogenetic tree may be categorised into three types. The types are differentiated by how many edges descend from the child of the root vertex.

**Definition** A *type-A clade* is one where the child of the root vertex has 3 or more descendant edges. A *type-B clade* is one where the child of the root vertex has exactly 2 descendant edges. If a type-$B$ clade contains exactly two leaves we call it a *cherry*. A *type-C clade* is one which consists of a single edge.

The tree in Figure 1 can itself be seen as a type-B clade. Figure 2 shows a tree $T = (V, E)$ with one of each of the clade types. Triangles represent subtrees

| $X$ | $\varphi(X)$ | $PD(X)$ | Value |
|---|---|---|---|
| $\{x_1\}$ | $\alpha(a) + \alpha(e) + \frac{1}{3}\alpha(z)$ | $L(a) + L(e) + L(z)$ | $2\frac{1}{3}$ |
| $\{x_1, x_2\}$ | $\alpha(a) + \alpha(b) + \alpha(e) + \frac{1}{3}\alpha(z)$ | $L(a) + L(b) + L(e) + L(z)$ | $3\frac{1}{3}$ |
| $\{x_2, x_3\}$ | $\alpha(b) + \alpha(c)$ $+\alpha(e) + \alpha(f) + \frac{5}{9}\alpha(z)$ | $L(b) + L(c)$ $+L(e) + L(f) + L(z)$ | $4\frac{5}{9}$ |
| $\{x_1, x_2, x_3\}$ | $\alpha(a) + \alpha(b) + \alpha(c)$ $+\alpha(e) + \alpha(f) + \frac{5}{9}\alpha(z)$ | $L(a) + L(b) + L(c)$ $+L(e) + L(f) + L(z)$ | $5\frac{5}{9}$ |
| $\{x_1, x_2, x_3, x_4\}$ | $\alpha(a) + \alpha(b) + \alpha(c) + \alpha(d)$ $+\alpha(e) + \alpha(f) + \frac{5}{9}\alpha(z)$ | $L(a) + L(b) + L(c) + L(d)$ $+L(e) + L(f) + L(z)$ | $4\frac{5}{9}$ |

Table 1: Expressions for $\varphi$ and PD for selected terminal vertex sets (leaf sets) from the tree $T$ in Figure 1. PD values are calculated for edge lengths given in the text, and in each case these match the $\varphi$ value for the same set.

of $T$, possibly containing no more than the single vertex indicated. The dotted line divides $E(T)$ into two parts, according to the conditions of Theorem 1.3.

Now suppose EvoHeritage attrition occurs along every edge of a phylogenetic tree. If $\varphi$ is to correspond to PD for every set of leaves of this tree, its structure is quite limited. Any descendant of the root can descend to at most two leaves. Figure 3 illustrates some of these limited structures, which contain only single-edge clades or $Y$-shaped ones. The results that follow formalise this idea.

**Proposition 1.2** *Let $T = (V, E)$ be a rooted phylogenetic tree, on terminal vertexset $S$, containing no monotomy vertices. Suppose $\beta(e) < 1$ for every edge $e \in E(T)$. Then $\varphi(X) = PD(X)$ for all sets $X \subseteq S$ if and only if the clades of $T$ are all cherries, single edges or a combination of both.*

The proof of Proposition 1.2 follows as a consequence of the following more general result.

**Theorem 1.3** *Let $T = (V, E)$ be a rooted phylogenetic tree containing no monotomy vertices. Let $S$ be the terminal vertex set of $T$, and $r$ denote the root vertex. Then $\varphi(X) = PD(X)$ for every set of leaves $X \subseteq S$ if and only if each of the following conditions hold:*

- *In type-A clades:*
  1. *Every edge $e$ not incident with the root has $\beta(e) = 1$.*
  2. *For the edge $z_A$ incident with the root, $0 \leq \beta(z_A) \leq 1$.*
  3. *For every edge $e$ in the clade, we have $\alpha(e) = L(e)$.*
- *In type-B clades:*

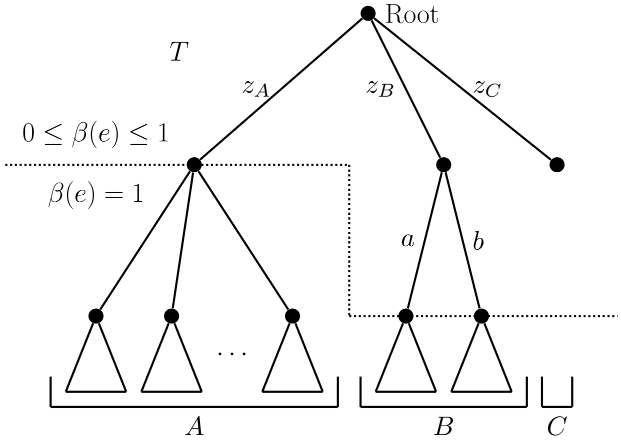

Figure 2: An example of the three clade types: A, B and C. Triangles represent sub trees. A clade of type B is considered a 'cherry' if it contains exactly two terminal vertices

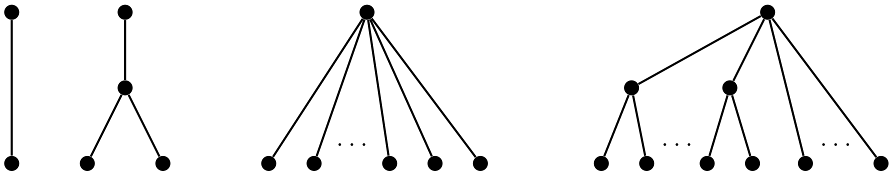

Figure 3: Examples of trees with EvoHeritage attrition occurring on every edge where $\varphi$ may equal PD for every set of leaves: two trivial trees consisting of only a single edge or a cherry, a star tree, a tree composed entirely of cherries and single-edge clades.

1. *Every edge e not incident with the child of the root has* $\beta(e) = 1$.
2. *For the edge* $z_B$ *incident with the root, and edges* $a, b$ *immediate descendants of* $z_B$*, we allow* $0 \leq \beta(z_B), \beta(a), \beta(b) \leq 1$*, provided that* $\alpha(z_B) = \frac{L(z_B)}{\beta(a)\beta(b)}$, $\alpha(a) = L(a) + L(z_B)(1 - \frac{1}{\beta(b)})$*, and* $\alpha(b) = L(b) + L(z_B)(1 - \frac{1}{\beta(a)})$.
3. *For every edge* $e \notin \{a, b, z_B\}$*, we have* $\alpha(e) = L(e)$.

- *In type-C clades:*
  1. *For the edge* $z_C$ *incident with the root,* $0 \leq \beta(z_C) \leq 1$.
  2. $\alpha(z_C) = L(z_C)$.

##### 1.4.4 Proof of Theorem 1.3

Let $\mathcal{T} = (V(\mathcal{T}), E(\mathcal{T}))$ be a clade of $T$ containing the edge $z = (r, v')$. Observe that terms derived from distinct clades do not interact in either PD or $\varphi$ calculations. As such, characterising the values of $\alpha(e)$ and $\beta(e)$ for each edge $e \in E(\mathcal{T})$ will characterise these for $T$ as a whole.

If $\mathcal{T}$ is a type-C clade, then it consists of only the edge $z$. The only set for which $\varphi$ needs to correspond to PD is $\{v'\}$. Hence $\alpha(z) = \varphi(\{v'\}) = PD(\{v'\}) = L(z)$. Otherwise, we suppose $\mathcal{T}$ contains at least 3 edges.

For a vertex $v$ of $\mathcal{T}$, we say it satisfies the $(\star)$ property if the following condition holds:

$(\star)$ Every edge $e$ in the subtree descending from $v$ has $\beta(e) = 1$.

Note that every terminal vertex of $\mathcal{T}$ satisfies $(\star)$. The proof proceeds by showing that the $(\star)$ property can be extended iteratively from the leaves to interior vertices apart from perhaps $v'$. Following this we prove that if $v'$ has three or more descendant edges, then it too has the $(\star)$ property. This means that we require $v'$ to have exactly two descendant edges for some edge (other than $z$) to exhibit EvoHeritage attrition.

Let $v$ be an interior vertex of $\mathcal{T}$ whose children all satisfy the $(\star)$ property. If $v$ is the root vertex, then the conditions of the theorem hold regardless of whether $\mathcal{T}$ is a type-A or type-B clade. Next suppose $v$ is not the root, and let $u$ be the parent vertex of $v$. We consider two cases: either $u$ is the root vertex or it is not.

**Case One:** Suppose $u$ is not the root vertex.

Since $u$ is not the root, it has a child vertex $w$ distinct from $v$. Select two leaves $x_1, x_2$ from the descendants of $v$ such that the path from $x_1$ (resp. $x_2$) to $v$ contains edge $a$ (resp. $b$) incident to $v$, and $a \neq b$. Further, select terminal vertex $x_3$ descended from $w$. Figure 4(a) illustrates these choices.

In calculating the $\varphi$ score of any subset of $\{x_1, x_2, x_3\}$, only those edges and paths shown can possibly make a non-zero contribution to the calculation.

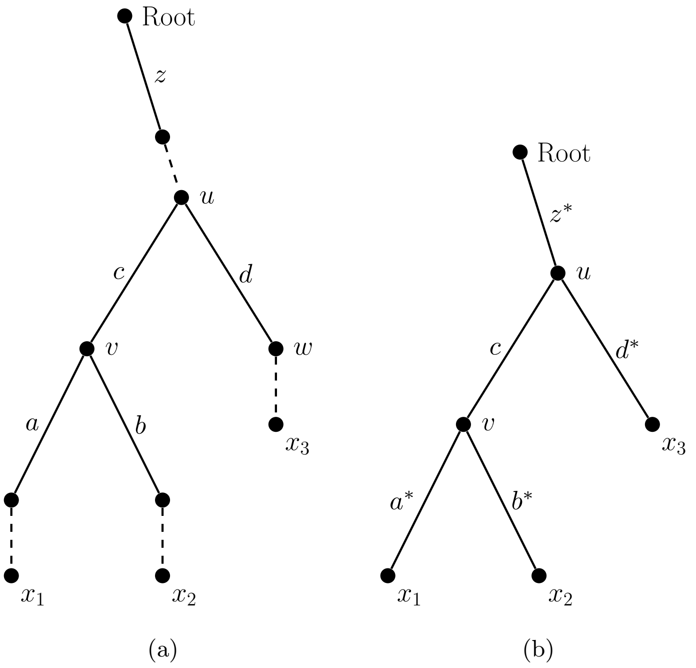

Figure 4: Selected parts of the tree $T$ referred to in the main narrative

Hence we can use Lemma 1.1 to replace the path from $v$ to $x_1$ (resp. $x_2$) with the edge $a^*$ (resp. $b^*$), noting that $\beta(a) = 1$ if and only if $\beta(a^*) = 1$, and $\beta(b) = 1$ if and only if $\beta(b^*) = 1$. Similarly, we replace the path from vertex $u$ to the root vertex (resp. $x_3$) with the edge $z^*$ (resp. $d^*$). The result of these changes is shown in Figure 4(b). Table 2 shows $\varphi$ and PD calculations for subsets of $\{x_1, x_2, x_3\}$, and a calculation of $\varphi(\{u\})$.

Observe that

$$\varphi(\{x_1\}) + \varphi(\{x_3\}) - \varphi(\{x_1, x_3\}) = \beta(a^*)\beta(c)\beta(d^*)\varphi(\{u\}). \qquad (42)$$

Now, under the assumption that $\varphi$ corresponds to PD for these sets, and noting also that $PD(\{x_1\}) + PD(\{x_3\}) - PD(\{x_1, x_3\}) = L(z^*) > 0$, it follows that the left hand side of (42), and hence the right hand side also, must be strictly positive. This implies each factor is nonzero: $\beta(a^*), \beta(c), \beta(d^*), \varphi(\{u\}) > 0$. Next, by expressing $L(a^*)$ in terms of $\varphi$, we will determine that $\beta(b^*) = 1$.

| $X$ | $\varphi(X)$ | $PD(X)$ |
|---|---|---|
| $\{u\}$ | $\alpha(z^*)$ | |
| $\{x_1\}$ | $\alpha(a^*) + \alpha(c)\beta(a^*) + \alpha(z^*)\beta(a^*)\beta(c)$ | $L(a^*) + L(c) + L(z^*)$ |
| $\{x_2\}$ | $\alpha(b^*) + \alpha(c)\beta(b^*) + \alpha(z^*)\beta(b^*)\beta(c)$ | $L(b^*) + L(c) + L(z^*)$ |
| $\{x_3\}$ | $\alpha(d^*) + \alpha(z^*)\beta(d^*)$ | $L(d^*) + L(z^*)$ |
| $\{x_1, x_2\}$ | $\alpha(a^*) + \alpha(b^*) + \alpha(c)[\beta(a^*) + \beta(b^*) - \beta(a^*)\beta(b^*)]$ $+\alpha(z^*)[\beta(a^*) + \beta(b^*) - \beta(a^*)\beta(b^*)]\beta(c)$ | $L(a^*) + L(b^*) + L(c) + L(z^*)$ |
| $\{x_1, x_3\}$ | $\alpha(a^*) + \alpha(d^*) + \alpha(c)\beta(a^*)$ $+\alpha(z^*)[\beta(d^*) + \beta(a^*)\beta(c) - \beta(a^*)\beta(c)\beta(d^*)]$ | $L(a^*) + L(c) + L(d^*) + L(z^*)$ |
| $\{x_2, x_3\}$ | $\alpha(b^*) + \alpha(d^*) + \alpha(c)\beta(b^*)$ $+\alpha(z^*)[\beta(d^*) + \beta(b^*)\beta(c) - \beta(b^*)\beta(c)\beta(d^*)]$ | $L(b^*) + L(c) + L(d^*) + L(z^*)$ |
| $\{x_1, x_2, x_3\}$ | $\alpha(a^*) + \alpha(b^*) + \alpha(d^*) + \alpha(c)[\beta(a^*) + \beta(b^*)$ $-\beta(a^*)\beta(b^*)] + \alpha(z^*)[\beta(d^*) + \beta(a^*)\beta(c)$ $+\beta(b^*)\beta(c) - \beta(a^*)\beta(b^*)\beta(c) - \beta(a^*)\beta(c)\beta(d^*)$ $-\beta(b^*)\beta(c)\beta(d^*) + \beta(a^*)\beta(b^*)\beta(c)\beta(d^*)]$ | $L(a^*) + L(b^*) + L(c) + L(d^*) + L(z^*)$ |

Table 2: $\varphi$, PD values for Figure 4(b)

Using Table 2:

$$\begin{aligned}L(a^*) &= PD(\{x_1, x_2\}) - PD(\{x_2\})\\ &= \varphi(\{x_1, x_2\}) - \varphi(\{x_2\})\\ &= \alpha(a^*) + \alpha(c)[\beta(a^*) - \beta(a^*)\beta(b^*)] + \alpha(z^*)[\beta(a^*)\beta(c) - \beta(a^*)\beta(b^*)\beta(c)]\end{aligned}$$

and
$$\begin{aligned}L(a^*) &= PD(\{x_1, x_2, x_3\}) - PD(\{x_2, x_3\})\\ &= \varphi(\{x_1, x_2, x_3\}) - \varphi(\{x_2, x_3\})\\ &= \alpha(a^*) + \alpha(c)[\beta(a^*) - \beta(a^*)\beta(b^*)] + \alpha(z^*)[\beta(a^*)\beta(c) - \beta(a^*)\beta(b^*)\beta(c)\\ &\quad - \beta(a^*)\beta(c)\beta(d^*) + \beta(a^*)\beta(b^*)\beta(c)\beta(d^*)]\end{aligned}$$

Hence $\alpha(z^*)\beta(a^*)\beta(c)\beta(d^*) = \alpha(z^*)\beta(a^*)\beta(b^*)\beta(c)\beta(d^*)$. As noted above, every factor on the left hand side of this equation is nonzero, therefore $\beta(b^*) = 1$. This in turn means we must have $\beta(b) = 1$.

A symmetric argument shows that $\beta(a) = 1$, and (repeating for other edges descending from $v$ if necessary) that $v$ has the ($\star$) property. By iterating this procedure, we find that every vertex, other than possibly $v'$ and the root, must satisfy the ($\star$) property.

**Case Two:** Suppose the parent of $v$ is the root vertex ($v = v'$), and $\mathcal{T}$ is a type-A clade.

We select $x_1, x_2, x_3$, three leaves of $\mathcal{T}$, such that the path from $x_1$ (resp. $x_2, x_3$) to $v$ contains edge $a$ (resp. $b, c$) incident to $v$, and $a, b, c$ are all distinct. Figure 5(a) illustrates these choices. As in Case One, we use Lemma 1.1 to replace the path from $v$ to $x_1$ (resp. $x_2, x_3$) with the edge $a^*$ (resp. $b^*, c^*$). The result of these changes is shown in Figure 5(b). Table 3 shows $\varphi$ and PD calculations for subsets of $\{x_1, x_2, x_3\}$, and a calculation of $\varphi(\{u\})$.

Observe that

$$\varphi(\{x_1\}) + \varphi(\{x_3\}) - \varphi(\{x_1, x_3\}) = \beta(a^*)\beta(c^*)\alpha(z). \tag{43}$$

Now, under the assumption that $\varphi$ corresponds to PD for these sets, and noting also that $PD(\{x_1\}) + PD(\{x_3\}) - PD(\{x_1, x_3\}) = L(z) > 0$, it follows that the left hand side of (43), and hence the right hand side also, must be strictly positive. This implies each factor is nonzero: $\beta(a^*), \beta(c^*), \varphi(\{v\}) > 0$. Again we express $L(a^*)$ in terms of two $\varphi$ expressions.

$$\begin{aligned}L(a^*) &= PD(\{x_1, x_2\}) - PD(\{x_2\})\\ &= \varphi(\{x_1, x_2\}) - \varphi(\{x_2\})\\ &= \alpha(a^*) + \alpha(z)[\beta(a^*) - \beta(a^*)\beta(b^*)]\end{aligned}$$

and
$$\begin{aligned}L(a^*) &= PD(\{x_1, x_2, x_3\}) - PD(\{x_2, x_3\})\\ &= \varphi(\{x_1, x_2, x_3\}) - \varphi(\{x_2, x_3\})\\ &= \alpha(a^*) + \alpha(z)[\beta(a^*) - \beta(a^*)\beta(b^*) - \beta(a^*)\beta(c^*) + \beta(a^*)\beta(b^*)\beta(c^*)]\end{aligned}$$

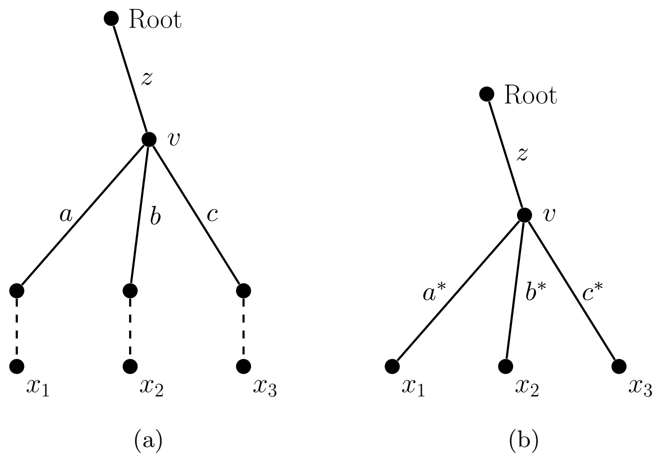

Figure 5: Diagram to show key edges and vertices referred to in the main narrative

| $X$ | $\varphi(X)$ | $PD(X)$ |
|---|---|---|
| $\{v\}$ | $\alpha(z)$ | |
| $\{x_1\}$ | $\alpha(a^*) + \alpha(z)\beta(a^*)$ | $L(a^*) + L(z)$ |
| $\{x_2\}$ | $\alpha(b^*) + \alpha(z)\beta(b^*)$ | $L(b^*) + L(z)$ |
| $\{x_3\}$ | $\alpha(c^*) + \alpha(z)\beta(c^*)$ | $L(c^*) + L(z)$ |
| $\{x_1, x_2\}$ | $\alpha(a^*) + \alpha(b^*) + \alpha(z)[\beta(a^*) + \beta(b^*) - \beta(a^*)\beta(b^*)]$ | $L(a^*) + L(b^*) + L(z)$ |
| $\{x_1, x_3\}$ | $\alpha(a^*) + \alpha(c^*) + \alpha(z)[\beta(a^*) + \beta(c^*) - \beta(a^*)\beta(c^*)]$ | $L(a^*) + L(c^*) + L(z)$ |
| $\{x_2, x_3\}$ | $\alpha(b^*) + \alpha(c^*) + \alpha(z)[\beta(b^*) + \beta(c^*) - \beta(b^*)\beta(c^*)]$ | $L(b^*) + L(c^*) + L(z)$ |
| $\{x_1, x_2, x_3\}$ | $\alpha(a^*) + \alpha(b^*) + \alpha(c^*) + \alpha(z)[\beta(a^*) + \beta(b^*) + \beta(c^*)$ $-\beta(a^*)\beta(b^*) - \beta(a^*)\beta(c^*) - \beta(b^*)\beta(c^*)$ $+\beta(a^*)\beta(b^*)\beta(c^*)]$ | $L(a^*) + L(b^*) + L(c^*) + L(z)$ |

Table 3: $\varphi$, PD values for Figure 5(b)

Hence $\alpha(z)\beta(a^*)\beta(c^*) = \alpha(z)\beta(a^*)\beta(b^*)\beta(c^*)$. Since every factor on the left hand side of the equation is nonzero, we get $\beta(b^*) = 1$. Symmetric arguments show that no EvoHeritage attrition occurs on any edge descending from $v$. Therefore every edge $e \in E(\mathcal{T})\backslash\{z\}$ has $\beta(e) = 1$ when $\mathcal{T}$ is a type-A clade.

Accordingly, $L(a^*) = \alpha(a^*) + \alpha(z)[\beta(a^*) - \beta(a^*)\beta(b^*)] = \alpha(a^*)$. Hence $\alpha(a) = L(a)$ (by Lemma 1.1). Again, by symmetry $\alpha(e) = L(e)$ for every edge $e \in E(\mathcal{T})\backslash\{z\}$. Finally, taking $X = \{x_1\}$ in Table 3 we obtain $\alpha(z) = L(z)$.

**Case Three:** Suppose the parent of $v$ is the root vertex ($v = v'$), and $\mathcal{T}$ is a type-B clade.

We proceed as in Case Two, using Figure 5 and Table 3 as if $L(c^*) = \alpha(c^*) = \beta(c^*) = 0$. By Case One and Lemma 1.1, $\beta(a^*) = \beta(a)$, and $\beta(b^*) = \beta(b)$. Then

$$\begin{aligned}\varphi(\{x_1\}) + \varphi(\{x_2\}) - \varphi(\{x_1, x_2\}) &= PD(\{x_1\}) + PD(\{x_2\}) - PD(\{x_1, x_2\}) \\ \alpha(z)\beta(a^*)\beta(b^*) &= L(z) \\ \alpha(z) &= \frac{L(z)}{\beta(a)\beta(b)}\end{aligned}$$

Furthermore,

$$\begin{aligned}\varphi(\{x_1, x_2\}) - \varphi(\{x_2\}) &= PD(\{x_1, x_2\}) - PD(\{x_2\}) \\ \alpha(a^*) + \frac{L(z)}{\beta(a)\beta(b)}(\beta(a) - \beta(a)\beta(b)) &= L(a^*) \\ \alpha(a^*) &= L(a^*) + L(z)\left(1 - \frac{1}{\beta(b)}\right) \\ \alpha(a) &= L(a) + L(z)\left(1 - \frac{1}{\beta(b)}\right) \qquad \text{[by Lemma 1.1]}\end{aligned}$$

A symmetric argument gives the required value of $\alpha(b)$.

We have now shown that if $\varphi(X) = PD(X)$ for every set $X \subseteq S$ then the conditions listed in the theorem hold. To complete the proof we must now show the converse. It can be shown by direct calculation that if the conditions listed in the theorem hold, then $\varphi(X) = PD(X)$ for every set $X \subseteq S$. For type-$A$ and type-$C$ clades this is immediate, since the $\varphi$ formula derived in the main body of the paper uses the value of neither $\beta(z_A)$ nor $\beta(z_C)$. There being no EvoHeritage preceding the root, there is no need to account for the rate at which it could be lost.

We introduce some shorthand for the type-$B$ case. Suppose $X$ is a set of leaves within a type-$B$ clade. Let $\Sigma_a$ denote that portion of $PD(X)$ which is derived from edges descended from $a$. We write $\Sigma_b$ similarly for edges descended from $b$.

Firstly, if $X$ is entirely descended from $a$:

$$\begin{aligned}\varphi(X) &= \Sigma_a + \alpha(a) + \alpha(z_B)\beta(a) \\ &= \Sigma_a + L(a) + L(z_B) - \frac{L(z_B)}{\beta(b)} + \frac{L(z_B)}{\beta(a)\beta(b)}\beta(a) \\ &= \Sigma_a + L(a) + L(z_B) \\ &= PD(X)\end{aligned}$$

A symmetric argument holds when $X$ is entirely descended from $b$. Finally, if $X$ is descended from both $a$ and $b$:

$$\begin{aligned}\varphi(X) &= \Sigma_a + \Sigma_b + \alpha(a) + \alpha(b) + \alpha(z_B)(\beta(a) + \beta(b) - \beta(a)\beta(b)) \\ &= \Sigma_a + \Sigma_b + L(a) + L(z_B) - \frac{L(z_B)}{\beta(b)} + L(b) + L(z_B) - \frac{L(z_B)}{\beta(a)} \\ &\quad + \frac{L(z_B)}{\beta(a)\beta(b)}(\beta(a) + \beta(b) - \beta(a)\beta(b)) \\ &= \Sigma_a + \Sigma_b + L(a) + L(b) + L(z_B) \\ &= PD(X)\end{aligned}$$

□

##### 1.4.5 Proof that maximising PD does not always maximise $\varphi_\rho$

The tree in Figure 6 provides an example where a set that maximises PD does not maximise $\varphi_{0.01}$ (i.e., $\varphi_\rho$ for $\rho = 0.01$). Consider choosing a set of four out of five species in Figure 6. If we wish to maximise PD, the one species we would exclude to leave a set of four would clearly be either $y$ or $z$. This gives a maximal PD of

$$\text{PD}(\{v, w, x, y\}) = 17.99 + 17.99 + 18 + 0.01 + 2 + 17.98 + 2.02 = 75.99. \quad (44)$$

By comparison, if we were to exclude $v$, we would be left with a PD score of

$$\text{PD}(\{w, x, y, z\}) = 17.99 + 0.01 + 18 + 2 + 2.02 + 17.98 + 17.98 = 75.98. \quad (45)$$

Now let us consider $\varphi_{0.01}$ on the same tree. In this case we do not add the origin of life as this does not change the comparison between the $\{v, w, x, y\}$ scenario and the $\{w, x, y, z\}$ scenario.

$$\begin{aligned}\varphi_{0.01}(\{v, w, x, y\}) =& 2 \cdot \frac{1 - e^{-0.1799}}{0.01} + \frac{1 - e^{-0.18}}{0.01} + \frac{1 - e^{-0.2}}{0.01} + \frac{1 - e^{-0.0001}}{0.01}\left(1 - (1 - e^{-0.1799})^2\right) \\ &+ \frac{1 - e^{-0.02}}{0.01}\left[1 - (1 - (1 - (1 - e^{-0.1799})^2)e^{-0.0001}) \cdot (1 - e^{-0.18})\right] \\ =& 69.51 \quad \text{(2 d.p.)}\end{aligned} \quad (46)$$

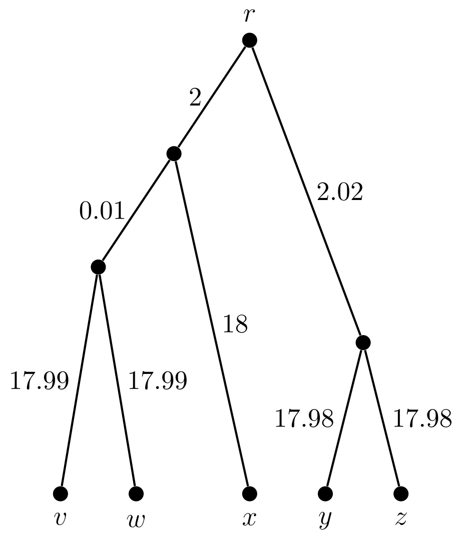

Figure 6: A binary phylogenetic tree, with edge lengths marked. A set of four leaves that maximises the PD score, $\{v, w, x, y\}$, does not maximise $\varphi_{0.01}$ for sets of this size. This set also does not maximise feature diversity under the stochastic model when $\rho = 0.01$. In both cases, the set $\{w, x, y, z\}$ gives larger values.

In contrast,

$$\begin{aligned}\varphi_{0.01}(\{w, x, y, z\}) =& 2 \cdot \frac{1 - e^{-0.18}}{0.01} + 2 \cdot \frac{1 - e^{-0.1798}}{0.01} \\ &+ \frac{1 - e^{-0.02}}{0.01}\left(1 - (1 - e^{-0.18})^2\right) + \frac{1 - e^{-0.0202}}{0.01}\left(1 - (1 - e^{-0.1798})^2\right) \\ =& 69.73 \quad \text{(2 d.p.)},\end{aligned} \tag{47}$$

and so $\varphi_{0.01}(\{w, x, y, z\}) > \varphi_{0.01}(\{v, w, x, y\})$. However, $\mathrm{PD}(\{w, x, y, z\}) < \mathrm{PD}(\{v, w, x, y\})$ so the set of nodes that maximises PD is not the same as that which maximises $\varphi_{0.01}$. Of course, this is a contrived example, and even so the set of vertices that maximise PD may often also maximise $\varphi_\rho$ depending on the value of $\rho$. What matters in practice, however, are the relative diversity values of a wide range of different sets of vertices (or species). In terms of this we expect $\varphi_\rho$ to behave very differently to PD. If we are performing an analysis based on more than just the tree, for example incorporating extinction risk as an additional distinguishing factor in a conservation prioritisation analysis, the effect of changing to $\varphi_\rho$ would be expected to be more substantial.

The same effect as above occurs within the framework of our stochastic model of feature diversity. For the stochastic comparison, we assume there are no features present at the root vertex of the tree in Figure 6. We calculate the expected feature diversity of the sets $\{v, w, x, y\}$ and $\{w, x, y, z\}$, with values of $\lambda = 1$ and $\rho = 0.01$.

$$\begin{aligned}\mathbb{E}[FD(\{v,w,x,y\})] =& \mathbb{E}[FD(v)] + \mathbb{E}[FD(w)] + \mathbb{E}[FD(x)] + \mathbb{E}[FD(y)] \\ &- \mathbb{E}[FD(v \cap w)] - \mathbb{E}[FD(v \cap x)] \\ &- \mathbb{E}[FD(w \cap x)] + \mathbb{E}[FD(v \cap w \cap x)] \\ =& 4 \cdot \frac{1}{0.01}(1 - e^{-0.2}) - 2 \cdot \frac{1}{0.01} e^{-0.36}(1 - e^{-0.02}) \\ &- \frac{1}{0.01} e^{-0.3599}(1 - e^{-0.0201}) + \frac{1}{0.01} e^{-0.5399}(1 - e^{-0.02}) \\ =& 69.51 \quad \text{(2 d.p.)}\end{aligned} \tag{48}$$

$$\begin{aligned}\mathbb{E}[FD(\{w,x,y,z\})] =& \mathbb{E}[FD(w)] + \mathbb{E}[FD(x)] + \mathbb{E}[FD(y)] + \mathbb{E}[FD(z)] \\ &- \mathbb{E}[FD(w \cap x)] - \mathbb{E}[FD(y \cap z)] \\ =& 4 \cdot \frac{1}{0.01}(1 - e^{-0.2}) - \frac{1}{0.01} e^{-0.36}(1 - e^{-0.02}) \\ &- \frac{1}{0.01} e^{-0.3596}(1 - e^{-0.0202}) \\ =& 69.73 \quad \text{(2 d.p.)}\end{aligned} \tag{49}$$

Similar to the deterministic setting above, our calculations illustrate that $\mathbb{E}[FD(\{w, x, y, z\})] > \mathbb{E}[FD(\{v, w, x, y\})]$, even though $\{v, w, x, y\}$ is a set of size four that maximises PD.

#### 1.5 EvoHeritage under stochastic conditions

Our stochastic model considers one form of EvoHeritage measured in discrete units. These discrete EvoHeritage units may be thought of as (presence / absence) features which arise and are lost along the edges of a phylogenetic tree. The features exhibited by a species are precisely those that persist until the terminal vertex representing it. Feature gains and losses will be described by exponential distributions, with respective rates $\lambda(e)$ and $\rho(e)$ along edge $e$. Here we consider the case where these rates have the same values for every edge in a tree, namely $\lambda$ and $\rho$.

We assume that the gain and loss of a feature is independent of the presence or absence of other features. We also require that a feature be gained before it can be lost, and that each feature is distinct from all others. Thus the development of a new feature is not conditional on the number of features already arisen. We phrase this as a 'constant birth' process. But the attrition of features does depend on how many features exist, so we have a 'linear death' process. We model these processes by a continuous time Markov chain, using constant birth and linear death rates, as shown in Figure 7. We note that this model is described in Queueing Theory as an infinite server queue, or $M/M/\infty$-queue. This connection provides a limiting distribution for the model, given by $p_j = e^{-\frac{\lambda}{\rho}} \frac{(\frac{\lambda}{\rho})^j}{j!}$, for $j \geq 0$.

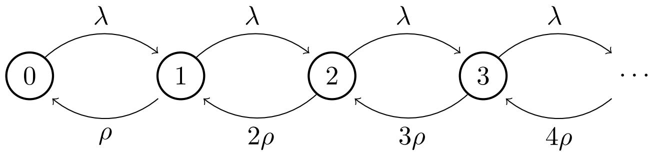

Figure 7: Diagram showing evolution of the system of feature gain and loss along a single edge under stochastic standard conditions. The numbers inside circles correspond to states of different total numbers of features. The arrows show transition between these states. $\lambda$ is the rate of new feature generation (accumulation). $\rho$ is the per feature rate of feature loss (attrition). In this system, more features means more opportunities for attrition.

Let $T = (V, E)$ be a rooted phylogenetic tree, where each edge $e \in E(T)$ has a corresponding positive length $L(e)$. For a vertex $v \in V$, we denote the set of features present at $v$ by $\mathbb{F}_v$. This set consists of both new features, those which have been gained on the incident ancestral edge, and persistent features which accumulated along earlier edges and survived the attrition process.

Let $N_e(t)$ be the net number of new features present on edge $e$ at distance $t$ from the start of $e$. Then $N_e(0) = 0$ and $N_e(t)$ is a Poisson random variable with mean $m_t = \int_0^t \lambda e^{-\rho s} ds = \frac{\lambda}{\rho} e^{-\rho s}\Big|_t^0 = \frac{\lambda}{\rho}(1 - e^{-\rho t})$, i.e. $N_e(t) \sim P_0(m_t)$. Hence $\mathbb{E}[N_e(t)] = m_t$.

We can now derive an expression for the expected number of features which arise along edge $e$ and are still present at the end of $e$.

$$\mathbb{E}[N_e(L(e))] = m_{L(e)} = \frac{\lambda}{\rho}(1 - e^{-\rho L(e)})$$

Next, let $v, v' \in V(T)$, where $v'$ is a child of $v$ and $(v, v') = e \in E(T)$. We write $F_v$ for the random variable that counts the number of features present at vertex $v$. Then we define $P_{v'}$ to be the number of these $F_v$ features which persist along the entirety of edge $e$ to be present at $v'$. Conditional on $F_v$, $P_{v'}$ is described by a binomial distribution $P_{v'} \sim \mathrm{Bin}(F_v, e^{-\rho L(e)})$, hence we have $\mathbb{E}[P_{v'}|F_v] = F_v e^{-\rho L(e)}$.

Finally, we can write $\mathbb{E}[F_{v'}|F_v] = F_v e^{-\rho L(e)} + \frac{\lambda}{\rho}(1 - e^{-\rho L(e)})$. When $\rho = 0$, we define $\mathbb{E}[F_{v'}|F_v]$ as $\lim_{\rho \to 0} \mathbb{E}[F_{v'}|F_v] = F_v + \lambda L(e)$.

**Lemma 1.4** *Let $T$ be a rooted phylogenetic tree, and let $P$ be a path in $T$, beginning at $v$ and ending at $v'$. Then $\mathbb{E}[F_{v'}|F_v]$ may be calculated as if $v$ and $v'$ were connected by a single edge whose length is the length of $P$ as a whole.*

##### 1.5.1 Proof of Lemma 1.4

Let $u, v, w \in V(T)$ and $e, f \in E(T)$. Suppose that $e = (u, v), f = (v, w)$. Then

$$\begin{aligned}\mathbb{E}[F_w|F_u] &= \mathbb{E}[F_v|F_u]e^{-\rho L(f)} + \frac{\lambda}{\rho}(1 - e^{-\rho L(f)}) \\ &= [F_u e^{-\rho L(e)} + \frac{\lambda}{\rho}(1 - e^{-\rho L(e)})]e^{-\rho L(f)} + \frac{\lambda}{\rho}(1 - e^{-\rho L(f)}) \\ &= F_u e^{-\rho(L(e)+L(f))} + \frac{\lambda}{\rho}(1 - e^{-\rho(L(e)+L(f))})\end{aligned}$$

So the expected number of features is the same if we take the entire path as if it were composed of a single edge of length $L(e) + L(f)$. This observation can be applied iteratively to extend to paths of any finite length. □

**Definition:** Let $T$ be a rooted phylogenetic tree with terminal vertex set $S$. The *feature diversity* (FD) of a set $X \subseteq S$ is the size of the set of features present among the taxa in $X$. Let $\mathbb{F}_X = \bigcup_{x \in X} \mathbb{F}_x$ and let $FD(X) = |\mathbb{F}_X|$. We note that FD by this definition is identical to the total EvoHeritage on a tree under stochastic standard conditions. Similar to the deterministic setting, we find that FD does not always match Phylogenetic Diversity (PD). Proposition 1.5 gives conditions for when the two measures do align. If $T$ does contain monotomy vertices, we suppress these first using Lemma 1.4.

**Theorem 1.5** *Let $T = (V, E)$ be a rooted phylogenetic tree containing no monotomy vertices, where $S$ is the terminal vertex set of $T$. Let $r$ denote the root vertex and let $L : E(T) \to \mathbb{R}$ and $L' : E(T) \to \mathbb{R}$ be two edge length assignments for $T$. Suppose at least one clade of $T$ contains more than 3 edges.*

*If $\mathbb{E}[FD_{T,L}(X)] = PD_{T,L'}(X)$ for every set of leaves $X \subseteq S$ then $\rho = 0$.*

*Conversely, if $\rho = 0$ and $L(e) = cL'(e)$ for every edge $e \in E(T)$ then $\mathbb{E}[FD_{T,L}(X)] = c\lambda PD_{T,L'}(X)$ for every set of leaves $X \subseteq S$.*

##### 1.5.2 Proof of Theorem 1.5

Let $\mathcal{T} = (V(\mathcal{T}), E(\mathcal{T}))$ be a clade of $T$ containing the edge $z = (r, v')$. We will determine that $\rho = 0$ or $v'$ has less than 3 descendant edges. Since terms derived from distinct clades do not interact in either PD or FD calculations, this will prove the result for $T$ as a whole. For simplicity's sake we consider the most natural scenario where the root vertex begins with no features. However this assumption may be relaxed without impacting the results shown below.

Let $v$ be an interior vertex of $\mathcal{T}$, and let $u$ be the parent vertex of $v$. We consider two cases: either $u$ is the root vertex or it is not.

**Case One:** Suppose $u$ is not the root vertex.

Since $u$ is not the root, it has a child vertex $w$ distinct from $v$. Select two leaves $x_1, x_2$ from the descendants of $v$ such that the path from $x_1$ (resp. $x_2$)

| $X$ | Expected FD of $X$ |
|---|---|
| $\lvert\mathbb{F}_1\rvert$ | $\frac{\lambda}{\rho}(1 - e^{-\rho(L(a^*)+L(c)+L(z^*))})$ |
| $\lvert\mathbb{F}_2\rvert$ | $\frac{\lambda}{\rho}(1 - e^{-\rho(L(b^*)+L(c)+L(z^*))})$ |
| $\lvert\mathbb{F}_3\rvert$ | $\frac{\lambda}{\rho}(1 - e^{-\rho(L(d^*)+L(z^*))})$ |
| $\lvert\mathbb{F}_1 \cap \mathbb{F}_3\rvert$ | $\frac{\lambda}{\rho}e^{-\rho(L(a^*)+L(c)+L(d^*))}(1 - e^{-\rho L(z^*)})$ |
| $\lvert\mathbb{F}_1 \cap \mathbb{F}_2 \cap \mathbb{F}_3\rvert$ | $\frac{\lambda}{\rho}e^{-\rho(L(a^*)+L(b^*)+L(c)+L(d^*))}(1 - e^{-\rho L(z^*)})$ |

Table 4: Expected FD for sets in Figure 4(b)

to $v$ contains edge $a$ (resp. $b$) incident to $v$, and $a \neq b$. Further, select terminal vertex $x_3$ descended from $w$. Figure 4(a) illustrates these choices.

In calculating FD of any subset of $\{x_1, x_2, x_3\}$, only those edges and paths shown can possibly make a non-zero contribution to the calculation. Hence we can use Lemma 1.4 to replace the path from $v$ to $x_1$ (resp. $x_2, x_3, r$) with the edge $a^*$ (resp $b^*, d^*, z^*$). The result of these changes is shown in Figure 4(b).

Writing $\mathbb{F}_i$ as shorthand in place of $\mathbb{F}_{x_i}$ for $i \in \{1, 2, 3\}$, Table 4 shows calculations of expected FD for selected sets. The expected FD for $|\mathbb{F}_1 \cap \mathbb{F}_3|$ is arrived at by noticing that any feature in this intersection must arise along $z^*$. Then it must survive attrition along edges $c$ and $a^*$ to be included in $\mathbb{F}_1$ and also along $d^*$ to be included in $\mathbb{F}_3$. The expression for $|\mathbb{F}_1 \cap \mathbb{F}_2 \cap \mathbb{F}_3|$ is obtained similarly. If FD corresponds to PD then the last two lines of Table 6 must be equal, because

$$PD(\{x_1, x_2\}) - PD(\{x_2\}) = L'(a^*) = PD(\{x_1, x_2, x_3\}) - PD(\{x_2, x_3\}).$$

Hence,

$$|\mathbb{F}_1 \cap \mathbb{F}_3| = |\mathbb{F}_1 \cap \mathbb{F}_2 \cap \mathbb{F}_3|$$
$$e^{-\rho(L(a^*)+L(c)+L(d^*))}(1 - e^{-\rho L(z^*)}) = e^{-\rho(L(a^*)+L(b^*)+L(c)+L(d^*))}(1 - e^{-\rho L(z^*)}).$$

Suppose $\rho \neq 0$. Then $1 = e^{-\rho L(b^*)}$, giving $L(b^*) = 0$ which contradicts our earlier stipulation that all edge lengths be strictly positive. Therefore $\rho = 0$ in this case.

**Case Two:** Suppose $u$ is the root vertex.

Assume $v$ has at least three descendant edges. We select $x_1, x_2, x_3$, three leaves of $\mathcal{T}$, such that the path from $x_1$ (resp. $x_2, x_3$) to $v$ contains edge $a$ (resp. $b, c$) incident to $v$, and $a, b, c$ are all distinct. Figure 5(a) illustrates these choices, and all edges relevant to the calculation of FD and PD appear in this figure. We use Lemma 1.4 to replace the path from $v$ to $x_1$ (resp. $x_2, x_3$) with the edge $a^*$ (resp. $b^*, c^*$). The result of these changes is shown in Figure 5(b).

| $X$ | Expected FD of $X$ |
|---|---|
| $\lvert\mathbb{F}_1\rvert$ | $\frac{\lambda}{\rho}(1 - e^{-\rho(L(a^*)+L(z))})$ |
| $\lvert\mathbb{F}_2\rvert$ | $\frac{\lambda}{\rho}(1 - e^{-\rho(L(b^*)+L(z))})$ |
| $\lvert\mathbb{F}_3\rvert$ | $\frac{\lambda}{\rho}(1 - e^{-\rho(L(c^*)+L(z))})$ |
| $\lvert\mathbb{F}_1 \cap \mathbb{F}_3\rvert$ | $\frac{\lambda}{\rho}e^{-\rho(L(a^*)+L(c^*))}(1 - e^{-\rho L(z)})$ |
| $\lvert\mathbb{F}_1 \cap \mathbb{F}_2 \cap \mathbb{F}_3\rvert$ | $\frac{\lambda}{\rho}e^{-\rho(L(a^*)+L(b^*)+L(c^*))}(1 - e^{-\rho L(z)})$ |

Table 5: Expected FD for sets in Figure 5(b)

| | |
|---|---|
| $\lvert\mathbb{F}_1 \cup \mathbb{F}_2\rvert$ | $\lvert\mathbb{F}_1\rvert + \lvert\mathbb{F}_2\rvert - \lvert\mathbb{F}_1 \cap \mathbb{F}_2\rvert$ |
| $\lvert\mathbb{F}_2 \cup \mathbb{F}_3\rvert$ | $\lvert\mathbb{F}_2\rvert + \lvert\mathbb{F}_3\rvert - \lvert\mathbb{F}_2 \cap \mathbb{F}_3\rvert$ |
| $\lvert\mathbb{F}_1 \cup \mathbb{F}_2 \cup \mathbb{F}_3\rvert$ | $\lvert\mathbb{F}_1\rvert + \lvert\mathbb{F}_2\rvert + \lvert\mathbb{F}_3\rvert - \lvert\mathbb{F}_1 \cap \mathbb{F}_2\rvert - \lvert\mathbb{F}_1 \cap \mathbb{F}_3\rvert - \lvert\mathbb{F}_2 \cap \mathbb{F}_3\rvert + \lvert\mathbb{F}_1 \cap \mathbb{F}_2 \cap \mathbb{F}_3\rvert$ |
| $\lvert\mathbb{F}_1 \cup \mathbb{F}_2\rvert - \lvert\mathbb{F}_2\rvert$ | $\lvert\mathbb{F}_1\rvert - \lvert\mathbb{F}_1 \cap \mathbb{F}_2\rvert$ |
| $\lvert\mathbb{F}_1 \cup \mathbb{F}_2 \cup \mathbb{F}_3\rvert - \lvert\mathbb{F}_2 \cup \mathbb{F}_3\rvert$ | $\lvert\mathbb{F}_1\rvert - \lvert\mathbb{F}_1 \cap \mathbb{F}_2\rvert - \lvert\mathbb{F}_1 \cap \mathbb{F}_3\rvert + \lvert\mathbb{F}_1 \cap \mathbb{F}_2 \cap \mathbb{F}_3\rvert$ |

Table 6: Equal-sized feature sets

As in Case One, we have

$$|\mathbb{F}_1 \cap \mathbb{F}_3| = |\mathbb{F}_1 \cap \mathbb{F}_2 \cap \mathbb{F}_3|$$
$$e^{-\rho(L(a^*)+L(c^*))}(1 - e^{-\rho L(z)}) = e^{-\rho(L(a^*)+L(b^*)+L(c^*))}(1 - e^{-\rho L(z)}).$$

Again, the supposition that $\rho \neq 0$ leads to the contradiction that $L(b^*) = 0$. So $\rho = 0$ when $v$ has at least three descendant edges.

The remaining possibility for $T$ is that every clade consists of a single edge, or an edge incident with the root and exactly two descending edges.

Now we prove the second statement of the theorem. Suppose that $\rho = 0$, and for some constant $c$, that $L(e) = cL'(e)$ for every $e \in E(T)$. Let $X = \{x_1, \ldots, x_k\}$ be a set of leaves from $T$. Let $p_i$ denote the path from the root vertex $r$ to terminal vertex $x_i$, and $P_X$ be the set of edges common to all $p_i$, for $1 \leq i \leq k$. Finally, let $Q_X = \bigcup p_i - P_X$, i.e. the set of edges contained in at least one of the paths $p_i$, but not all of them. Then

$$\mathbb{E}\left[|\bigcap_{x_i \in X} \mathbb{F}_{x_i}|\right] = \lim_{\rho \to 0} \frac{\lambda}{\rho} e^{-\rho \sum_{e \in Q_X} L(e)} \left(1 - e^{-\rho \sum_{e \in P_X} L(e)}\right) \tag{50}$$
$$= \lambda \sum_{e \in P_X} L(e) \tag{51}$$

For tree $T$ and edge length assignment $L$,

$$\begin{aligned}\mathbb{E}[FD(X)] &= \mathbb{E}[|\mathbb{F}_X|]\\ &= \mathbb{E}[|\mathbb{F}_{x_1} \cup \cdots \cup \mathbb{F}_{x_k}|]\\ &= \mathbb{E}[|\mathbb{F}_{x_1}|] + \cdots + \mathbb{E}[|\mathbb{F}_{x_k}|]\\ &\quad - \mathbb{E}[|\mathbb{F}_{x_1} \cap \mathbb{F}_{x_2}|] - \mathbb{E}[|\mathbb{F}_{x_1} \cap \mathbb{F}_{x_3}|] - \cdots - \mathbb{E}[|\mathbb{F}_{x_{k-1}} \cap \mathbb{F}_{x_k}|]\\ &\quad + \ldots\\ &\quad \pm \mathbb{E}[|\mathbb{F}_{x_1} \cap \cdots \cap \mathbb{F}_{x_k}|]\end{aligned}$$

Now let $e$ be an edge in $\bigcup p_i$, from which $m$ members of $X$ descend. When the final expression above is evaluated using Eqn. (51), then $L(e)$ is added a net $\binom{m}{1} - \binom{m}{2} + \binom{m}{3} - \cdots \pm \binom{m}{m}$ times. For all $m \geq 1$, we have $\sum_{j=0}^{m} (-1)^j \binom{m}{j} = 0$, and $\binom{m}{0} = 1$. Therefore $L(e)$ is added exactly once more than it is subtracted.

Hence $\mathbb{E}[FD_{T,L}(X)] = \lambda \sum_{e \in \cup p_i} L(e) = \lambda PD_{T,L}(X) = c\lambda PD_{T,L'}(X)$.

□

#### 1.6 Generalisations of EvoHeritage to multiple independent forms

The measures discussed here may be straightforwardly extended to account for different 'forms' of EvoHeritage. We shall consider EvoHeritage to be of the same 'form' if it is subject to the same rules of accumulation and attrition across an EvoHeritage graph. We may specify one or more forms of EvoHeritage, each with its own independent rules and characteristics of accumulation and attrition. For example, we may have 'immortal' EvoHeritage as one form that is never lost, to enable rare and truly irreversible evolutionary steps to be accounted for. The accumulation and attrition vaules for different forms of EvoHeritage may be indicated with subscripts, writing $\alpha_i(e)$ for the accumulation of EvoHeritage of form $i$ along edge $e$.

The formulae presented in both the main text and supplementary information all contain a single form of EvoHeritage. Each measure in the EvoHeritage calculus can be adjusted to incorporate the different forms of EvoHeritage. We simply calculate the measure of interest separately for each form and then sum these values together. As each form of EvoHeritage acts independent of each other, this gives the value across the set of forms. For instance, the equation for $\varphi$ becomes:

$$\varphi(X) = \sum_{i \in I} \sum_{e \in E} \alpha_i(e) \cdot p_i(d^V(e), X) \qquad (52)$$

In words, $\varphi$ of a set of vertices $X$ is the sum, across all forms of EvoHeritage (first summation term) and across all edges (second summation term) in the EvoHeritage graph. Inside the sum is the EvoHeritage of form $i$ accumulated along each each edge, $\alpha_i(e)$, multiplied by the proportion of that EvoHeritage surviving attrition to be counted in the set $X$ (using the $p_i$ expression relevant to this form).

### 2 Developing a complete EvoHeritage calculus

#### 2.1 Relationship between diff-$\varphi$ and classic phylogenetic distance for a pair of vertices

Suppose we measure phylogenetic distance as the date $D$ of common ancestry between two terminal vertices $x$ and $y$ on a dated phylogenetic tree. We seek to calculate EvoHeritage difference, diff-$\varphi$, between these terminal vertices under standard conditions with attrition at rate $\rho$. First, we express the phylogenetic tree as an EvoHeritage graph under standard conditions. Next we collapse all vertices except those corresponding to the two species, their common ancestor, and the root vertex at age $d \geq D$. By setting $X = \{x\}$ and $Y = \{y\}$ we are left with the following calculation for diff-$\varphi$.

$$\text{diff-}\varphi(\{x\},\{y\}) = \frac{\text{unique-}\varphi(\{x\},\{y\}) + \text{unique-}\varphi(\{y\},\{x\})}{\varphi(\{x,y\})} \tag{53}$$

Which, by expanding the unique-$\varphi$ components can be written as

$$\text{diff-}\varphi(\{x\},\{y\}) = \frac{2 \cdot \varphi(\{x,y\}) - \varphi(\{x\}) - \varphi(\{y\})}{\varphi(\{x,y\})} \tag{54}$$

Note that we could measure $\varphi$ in standardised units but as we are taking a ratio of different $\varphi$ measures any effect of units would immediately cancel out. We determine expressions for the terms appearing on the right hand side of the above equation. Let us begin with $\varphi(\{a\})$ and $\varphi(\{b\})$:

$$\begin{aligned}\varphi(\{a\}) = \varphi(\{b\}) &= \frac{\lambda}{\rho}(1 - e^{-\rho \cdot D}) + \frac{\lambda}{\rho}(1 - e^{-\rho \cdot (\tau - D)}) \cdot e^{-\rho \cdot D} \\ &= \frac{\lambda}{\rho}(1 - e^{-\rho \cdot \tau})\end{aligned} \tag{55}$$

Where $\tau$ is the date of the origin of life as described earlier. Now let us consider $\varphi(\{a, b\})$:

$$
\begin{aligned}
\varphi(\{a,b\}) &= 2\frac{\lambda}{\rho}(1-e^{-\rho\cdot D}) + \frac{\lambda}{\rho}(1-e^{-\rho\cdot(\tau-D)})\cdot(1-(1-e^{-\rho\cdot D})^2) \\
&= \frac{\lambda}{\rho}(2(1-e^{-\rho\cdot D}) + (1-e^{-\rho\cdot(\tau-D)})\cdot(1-(1-e^{-\rho\cdot D})^2)) \\
&= \frac{\lambda}{\rho}(2(1-e^{-\rho\cdot D}) + (1-e^{-\rho\cdot(\tau-D)})\cdot(2e^{-\rho\cdot D}-e^{-2\rho\cdot D})) \\
&= \frac{\lambda}{\rho}(2-2e^{-\rho\cdot D}+2e^{-\rho\cdot D}-e^{-2\rho\cdot D}-e^{-\rho\cdot(\tau-D)}\cdot(2e^{-\rho\cdot D}-e^{-2\rho\cdot D})) \\
&= \frac{\lambda}{\rho}(2-e^{-2\rho\cdot D}-2e^{-\rho\cdot\tau}+e^{-\rho\cdot(\tau+D)})
\end{aligned}
\tag{56}
$$

Substituting these expressions into Eqn. (54) and cancelling out the factor of $\frac{\lambda}{\rho}$ from both the numerator and demomenator gives:

$$
\begin{aligned}
\text{diff-}\varphi(\{a\},\{b\}) =& \frac{2(2-e^{-2\rho\cdot D}-2e^{-\rho\cdot\tau}+e^{-\rho\cdot(\tau+D)})-2(1-e^{-\rho\cdot\tau})}{2-e^{-2\rho\cdot D}-2e^{-\rho\cdot\tau}+e^{-\rho\cdot(\tau+D)}} \\
=& \frac{4-2e^{-2\rho\cdot D}-4e^{-\rho\cdot\tau}+2e^{-\rho\cdot(\tau+D)}-2+2e^{-\rho\cdot\tau}}{2-e^{-2\rho\cdot D}-2e^{-\rho\cdot\tau}+e^{-\rho\cdot(\tau+D)}} \\
=& \frac{2-2e^{-2\rho\cdot D}-2e^{-\rho\cdot\tau}+2e^{-\rho\cdot(\tau+D)}}{2-e^{-2\rho\cdot D}-2e^{-\rho\cdot\tau}+e^{-\rho\cdot(\tau+D)}}
\end{aligned}
\tag{57}
$$

Let us consider what happens in the limit as $\rho \to \infty$. All of the exponential terms tend to 0 and therefore

$$
\lim_{\rho\to\infty} \text{diff-}\varphi(\{a\},\{b\}) = \frac{2}{2} = 1. \tag{58}
$$

Recall that diff-$\varphi$ is the proportion of total EvoHeritage across two sets which is unique to one set or the other. The above limit conforms with the intuition that with infinite EvoHeritage attrition, these two species will be maximally distant from one another. Now consider a fixed finite $\rho > 0$, if we set $D = \tau$ then diff-$\varphi(\{a\},\{b\}) = 1$ again, its maximal value. As $D$ approaches $\tau$ diff-$\varphi$ should saturate so that increasing $D$ further towards $\tau$ has negligible impact.

Now, let us consider what happens when $\rho \to 0$. In this case substituting $\rho = 0$ into Eqn. (57) gives an indeterminate expression $(\frac{0}{0})$. We use L'Hôpital's Rule to calculate this limit.

$$
\begin{aligned}
\lim_{\rho\to 0}\text{diff-}\varphi(\{a\},\{b\}) &= \lim_{\rho\to 0}\frac{2-2e^{-2\rho\cdot D}-2e^{-\rho\cdot\tau}+2e^{-\rho\cdot(\tau+D)}}{2-e^{-2\rho\cdot D}-2e^{-\rho\cdot\tau}+e^{-\rho\cdot(\tau+D)}}\\
&= \lim_{\rho\to 0}\frac{4De^{-2\rho\cdot D}+2\tau e^{-\rho\cdot\tau}-2(\tau+D)e^{-\rho\cdot(\tau+D)}}{2De^{-2\rho\cdot D}+2\tau e^{-\rho\cdot\tau}-(\tau+D)e^{-\rho\cdot(\tau+D)}}\\
&= \frac{4D+2\tau-2(\tau+D)}{2D+2\tau-(\tau+D)}\\
&= \frac{2D}{\tau+D}
\end{aligned}
\tag{59}
$$

Which has classic phylogenetic distance on the numerator but normalises by the total PD of the two vertices together with the root. Again, if we set $D=\tau$ then diff-$\varphi(\{a\},\{b\})=1$ as expected.

#### 2.2 Relationship to genetic diversity

Let $T_u$ be an unrooted binary phylogenetic tree with $n$ leaves. Let $\{e_1,\ldots,e_{2n-3}\}$ be the set of edges in $T_u$. For each edge $e_i$ in this set, let $0\le L(e_i)\le 1$ be the length of $e_i$, corresponding to the probability of a difference in character state between vertices at either end of $e_i$.

The genetic diversity measure, introduced by (Crozier, 1992), measures the probability of there being two or more character states across the $n$ leaves. The measure is given by Equation 3 in (Crozier, 1992):

$$
P(\ge 2) = 1-\prod_{k=1}^{2n-3}(1-b_k), \tag{60}
$$

where $b_k$ gives the probability of a difference along a branch. So $1-b_k$ gives the probability that both endpoints of edge $e_k$ have matching character states, and hence the product term gives the probability that the character states match on every vertex.

To see how this corresponds to a measure in the $\varphi$ world first add a stem and root to the tree from any interior vertex and direct all edges away from the root. Let us refer to this new edge as having index $2n-2$ so that we can write the edge as $e_{2n-2}$ with all the other edges being indexed as before. Now set

$$
\alpha(e_i) = \begin{cases} 1, & \text{when } i = 2n-2;\\ 0, & \text{otherwise.}\end{cases} \tag{61}
$$

$$
\beta(e_i) = \begin{cases} 0, & \text{when } i = 2n-2;\\ b_i, & \text{otherwise.}\end{cases} \tag{62}
$$

The proportion of ancestral EvoHeritage (from edge $e_{2n-2}$) that *does* survive to all terminal vertices is the amount that survives EvoHeritage attrition on all branches other than the stem (which is where the EvoHeritage is generated). This is given by

$$\prod_{k=1}^{2n-3}(1-\beta(e_k)) \tag{63}$$

The proportion that *does not* survive is just one minus that giving

$$1-\prod_{k=1}^{2n-3}(1-\beta(e_k)) = 1-\prod_{k=1}^{2n-3}(1-b_k) = P(\geq 2) \tag{64}$$

And hence the relationship to genetic diversity from the main text is proven.

#### 2.3 Proof that the power set based calculation of $\varphi$ is equivalent to direct calculation

We will make use of the following lemma in the proof of Theorem 3 from the main text

**Lemma 2.1** *For a set $X$, and function $\beta : X \to \mathbb{R}$:*

$$\sum_{G\in\mathcal{P}(X)}\prod_{g\in G}\beta(g)\prod_{g\in X\setminus G}(1-\beta(g)) = 1$$

##### 2.3.1 Proof of Lemma 2.1

$$1 = \prod_{g\in X}(\beta(g)+(1-\beta(g))) = \sum_{G\in\mathcal{P}(X)}\prod_{g\in G}\beta(g)\prod_{g\in X\setminus G}(1-\beta(g)) \qquad \square$$

To prove Theorem 4 from the main text, we first recall the recursive formulation of $\varphi$.

$$\varphi(X) = \sum_{e\in E}\alpha(e)\cdot p(d^V(e), X), \tag{65}$$

where $p(v, X)$ is given by

$$p(v,X) = \begin{cases} 1, & \text{when } v \in X; \\ 0, & \text{when } v \notin X \text{ and } D^E(v) = \emptyset; \\ 1-\prod_{j\in D^E(v)}[1-p(d^V(j),X)\cdot\beta(j)], & \text{otherwise.} \end{cases} \tag{66}$$

On the other hand, starting with the expression for $\kappa(\mathbb{I}, X)$ and rearranging the order of summation we get:

$$\kappa(\mathbb{I}, X) = \sum_{G \in \mathcal{P}(E)} \left( \prod_{g \in G} \beta(g) \prod_{g \in E \setminus G} (1 - \beta(g)) \sum_{e \in E} \alpha(e) \cdot \mathbb{I}(d^V(e), G, X) \right)$$

$$= \sum_{e \in E} \alpha(e) \cdot \sum_{G \in \mathcal{P}(E)} \left( \prod_{g \in G} \beta(g) \prod_{g \in E \setminus G} (1 - \beta(g)) \cdot \mathbb{I}(d^V(e), G, X) \right)$$

Hence, the theorem holds provided, for every edge $e \in E$, we can show

$$p(d^V(e), X) = \sum_{G \in \mathcal{P}(E)} \left( \prod_{g \in G} \beta(g) \prod_{g \in E \setminus G} (1 - \beta(g)) \cdot \mathbb{I}(d^V(e), G, X) \right). \tag{67}$$

Recall from that main text that we defined the indicator function $\mathbb{I} : V \times \mathcal{P}(E) \times \mathcal{P}(V) \to \{0, 1\}$ by:

$$\mathbb{I}(v, G, X) = \begin{cases} 1, & \text{when } c(v, G, X) \geq 1; \\ 0, & \text{when } c(v, G, X) = 0. \end{cases} \tag{68}$$

$$c(v, G, X) = |\{x \in X : \text{there is a path from } v \text{ to } x \text{ using only edges in } G\}|. \tag{69}$$

The proof proceeds by induction on the number of edges in $E$. As the base case we consider the phylogentic tree with one edge, which we call $e$. Suppose the unique terminal vertex is $v = d^V(e)$. Note that $\mathbb{I}(v, G, X) = 1$ if and only if $v \in X$, for both $G = \emptyset$ and $G = \{e\}$. So in this case, beginning from the right hand side of Eqn. (67):

$$\sum_{G \in \mathcal{P}(E)} \left( \prod_{g \in G} \beta(g) \prod_{g \in E \setminus G} (1 - \beta(g)) \cdot \mathbb{I}(d^V(e), G, X) \right) \tag{70}$$

$$= \prod_{g \in \emptyset} \beta(g) \prod_{g \in \{e\}} (1 - \beta(g)) \cdot \mathbb{I}(v, \emptyset, X) + \prod_{g \in \{e\}} \beta(g) \prod_{g \in \emptyset} (1 - \beta(g)) \cdot \mathbb{I}(v, \{e\}, X) \tag{71}$$

$$= (1 - \beta(e)) \cdot \mathbb{I}(v, \emptyset, X) + \beta(e) \cdot \mathbb{I}(v, \{e\}, X) \tag{72}$$

$$= \begin{cases} 1, & \text{when } v \in X \\ 0, & \text{when } v \notin X \end{cases} \tag{73}$$

$$= p(v, X), \tag{74}$$

and the base case holds.

Now suppose that $T = (V, E)$ is a phylogenetic tree, and that Eqn. (67) holds for all phylogenetic trees with less than $|E|$ edges. We select an edge $e \in E$ and split up the sum on the right hand side of Eqn. (67) into two parts. The first part consists of subsets of $E$ that contain $e$, and the second consists of those subsets of $E$ which exclude $e$. In this way, we create the following expression:

$$
\begin{aligned}
&(1-\beta(e)) \cdot \sum_{G \in \mathcal{P}(E \backslash \{e\})} \left( \prod_{g \in G} \beta(g) \prod_{g \in E \backslash (G \cup \{e\})} (1-\beta(g)) \cdot \mathbb{I}(d^V(e), G, X) \right) \\
&+ \beta(e) \cdot \sum_{G \in \mathcal{P}(E \backslash \{e\})} \left( \prod_{g \in G} \beta(g) \prod_{g \in E \backslash (G \cup \{e\})} (1-\beta(g)) \cdot \mathbb{I}(d^V(e), G \cup \{e\}, X) \right).
\end{aligned} \tag{75}
$$

We consider two cases: either $d^V(e) \in X$ or $d^V(e) \notin X$.

Firstly, assume $d^V(e) \in X$. Then $\mathbb{I}(d^V(e), G, X) = \mathbb{I}(d^V(e), G \cup \{e\}, X) = 1$ for all $G \in \mathcal{P}(E \backslash \{e\}$, because we consider the path (with no edges) from $d^V(e)$ to itself to be a valid path. Then the expression (75) is equal to 1 by Lemma 2.1. Since $p(d^V(e), X) = 1$ whenever $d^V(e) \in X$, we have shown that Eqn. (67) holds in this case.

Now assume $d^V(e) \notin X$. Then for every set $G$ of edges, $\mathbb{I}(d^V(e), G, X) = \mathbb{I}(d^V(e), G \cup \{e\}, X)$ because $e$ doesn't help us to connect to any vertex in $X$ which was not already reachable via edges in $G$. Then the expression (75) becomes

$$
\sum_{G \in \mathcal{P}(E \backslash \{e\})} \left( \prod_{g \in G} \beta(g) \prod_{g \in E \backslash G} (1-\beta(g)) \cdot \mathbb{I}(d^V(e), G, X) \right). \tag{76}
$$

By the induction assumption this expression is equal to $p(d^V(e), X)$, and Eqn. (67) holds.

#### 2.4 Summing the partitioned-$\varphi$ of individual vertices

Many practical applications will find it useful to partition the EvoHeritage of all extant vertices between each individual extant vertex, however, we could use the same solution to partition between any mixture of extant and extinct vertices. This is not made explicit when partitioning classic PD, for example with the Evolutionary Distinctiveness (ED), or Equal Splits (ES) methods, because extinct species are never considered.

The sum of all partitioned-$\varphi$ values returns the total $\varphi$ of all extant vertices together: $\varphi(X) = \sum_{x \in X}$ partitioned-$\varphi(\{x\}, X)$. To see this, note that the total number of paths $c(v, G, X)$ can be found by calculating the number of paths from $v$ to each distinct $b$ in $X$, then summing these values together. In symbols: $c(v, G, X) = \sum\limits_{x \in X} c(v, G, \{x\})$.

$$\sum_{x \in X} \text{partitioned-}\varphi(\{x\}, X) \tag{77}$$
$$= \sum_{x \in X} \kappa(R, (\{x\}, X) \tag{78}$$
$$= \sum_{x \in X} \sum_{G \in \mathcal{P}(E)} \left( \prod_{g \in G} \beta(g) \prod_{g \in E \setminus G} (1 - \beta(g)) \sum_{e \in E} \alpha(e) \cdot R(d^V(e), G, \{x\}, X) \right) \tag{79}$$
$$= \sum_{G \in \mathcal{P}(E)} \left( \prod_{g \in G} \beta(g) \prod_{g \in E \setminus G} (1 - \beta(g)) \sum_{e \in E} \alpha(e) \cdot \frac{\sum_{x \in X} c(d^V(e), G, \{x\})}{c(d^V(e), G, X)} \right) \tag{80}$$
$$= \sum_{G \in \mathcal{P}(E)} \left( \prod_{g \in G} \beta(g) \prod_{g \in E \setminus G} (1 - \beta(g)) \sum_{e \in E} \alpha(e) \cdot \mathbb{I}(d^V(e), G, X) \right) \tag{81}$$
$$= \varphi(X) \tag{82}$$

#### 2.5 Values of partitioned-$\varphi$ and unique-$\varphi$ for extremal values of attrition

Let $T = (V, E)$ be an EvoHeritage tree, and for each edge $e \in E$, let $L(e)$ give the length of $e$. Let $X \subset V$ be the set of terminal vertices of $T$. Suppose $x$ is a terminal vertex of $T$, and that $e_x$ is its corresponding terminal edge. We write $ED(x)$ for the evolutionary distinctiveness score of $x$ in the tree $T$, and $\mathcal{D}(e)$ for the set of terminal vertices descended from edge $e$.

**Proposition 2.2** *Let $T$ be an EvoHeritage tree under standard conditions, with attrition parameter $\rho$. The extremal values of partitioned-$\varphi_\rho$ and unique-$\varphi_\rho$ are given by:*

- partitioned-$\varphi_0(\{x\}, X) = ED(x)$
- unique-$\varphi_0(\{x\}, X - \{x\}) = L(e_x)$
- partitioned-$\varphi_\infty(\{x\}, X) = 1$
- unique-$\varphi_\infty(\{x\}, X - \{x\}) = 1$

Recall that $\varphi(X) = \kappa(\mathbb{I}, X)$ and partitioned-$\varphi(X, Y) = \kappa(R, (X, Y))$. We begin by proving the statements where $\rho = 0$. From section 1.4.2 we have $\alpha(e) = \lambda L(e)$ and $\beta(e) = 1$ for all $e \in E$. Also, note that in this case $S_u = \lambda$.

Whenever $G \neq E$, the product $\prod_{g \in E \setminus G} (1 - \beta(g))$ is nonempty, and equals zero, making these terms vanish in the relevant $\kappa$ expressions. For $G = E$, the function $c(d^V(e), G, X)$ equals the number of terminal vertices in $X$ that descend from $e$, that is: $|\mathcal{D}(e)|$. In particular, $c(d^V(e), G, \{x\})$ equals one when

$x$ is descended from $e$, and zero for all other edges. Therefore, dividing through by $S_u$ to get a standardised value,

$$\begin{aligned}\frac{\text{partitioned-}\varphi_0(\{x\},X)}{S_u} &= \frac{\kappa(R,(\{x\},X))}{S_u} \\ &= \frac{1}{\lambda}\prod_{e\in E}1\prod_{e\in\emptyset}0\sum_{e\in E}\lambda L(e)R(d^V(e),E,\{x\},X) \\ &= \sum_{e\in E}L(e)R(d^V(e),E,\{x\},X) \\ &= \sum_{\substack{e\in E,\\ x\in\mathcal{D}(e)}}L(e)\frac{1}{|\mathcal{D}(e)|} \\ &= ED(x).\end{aligned} \tag{83}$$

Let $\bar{E}(x)$ be the set of edges that have a descendant terminal vertex distinct from $x$. Then, noting that $\{x\}\cup(X-\{x\})=X$,

$$\begin{aligned}\frac{\text{unique-}\varphi_0(\{x\},X-\{x\})}{S_u} &= \frac{\varphi_0(X)-\varphi_0(X-\{x\})}{S_u} \\ &= \frac{1}{\lambda}\left(\kappa(\mathbb{I},X)-\kappa(\mathbb{I},X-\{x\})\right) \\ &= \frac{1}{\lambda}\left(\sum_{e\in E}\lambda L(e)-\sum_{e\in\bar{E}(x)}\lambda L(e)\right) \\ &= L(e_x).\end{aligned} \tag{84}$$

Next, we prove the statements where $\rho\to\infty$. From section 1.4.1, as $\rho$ increases we have $\alpha(e)\approx\frac{\lambda}{\rho}$ and $\beta(e)\to 0$ for all $e\in E$. We use the approximation $S_u\approx\frac{\lambda}{\rho}$, valid for large values of $\rho$.

Whenever $G\neq\emptyset$, the product $\prod_{g\in E\setminus G}(\beta(g))$ is nonempty, and equals zero, making these terms vanish in the relevant $\kappa$ expressions. For $G=\emptyset$, the function $c(d^V(e),G,X)$ equals one if and only if $d^V(e)\in X$, and equals zero otherwise. Therefore, dividing through by $S_u$ to get a standardised value,

$$\begin{aligned}
\frac{\text{partitioned-}\varphi_\infty(\{x\},X)}{S_u} &= \frac{\kappa(R,(\{x\},X))}{S_u} \\
&= \frac{\prod_{e\in\emptyset}0\prod_{e\in E}1\sum_{e\in E}\alpha(e)R(d^V(e),\emptyset,\{x\},X)}{S_u} \\
&= \frac{\alpha(e_x)}{S_u} \\
&\approx \frac{\frac{\lambda}{\rho}}{\frac{\lambda}{\rho}} = 1
\end{aligned} \tag{85}$$

and

$$\begin{aligned}
\frac{\text{unique-}\varphi_0(\{x\},X-\{x\})}{S_u} &= \frac{\varphi_0(X)-\varphi_0(X-\{x\})}{S_u} \\
&= \frac{\kappa(\mathbb{I},X)-\kappa(\mathbb{I},X-\{x\})}{S_u} \\
&= \frac{\sum_{e\in E}\alpha(e)\mathbb{I}(d^V(e),\emptyset,X)-\sum_{e\in E}\alpha(e)\mathbb{I}(d^V(e),\emptyset,X-\{x\})}{S_u} \\
&= \frac{\alpha(e_x)}{S_u} \\
&\approx \frac{\frac{1}{\rho}}{\frac{1}{\rho}} = 1.
\end{aligned} \tag{86}$$

#### 2.6 Complexity of calculating partitioned-$\varphi$

In this section we show that the calculation of partitioned-$\varphi$ on a tree with $n$ leaves can in some cases require the addition of $2^{n-2}$ terms. Hence, this calculation cannot be completed in polynomial time in $n$. Recall the forms of the function $\kappa$, the path-count function $c$ and the path-ratio function $R$ from the main text.

$$\kappa(F,X) = \sum_{G\in\mathcal{P}(E)}\left(\prod_{g\in G}\beta(g)\prod_{g\in E\setminus G}(1-\beta(g))\sum_{e\in E}\alpha(e)\cdot F(d^V(e),G,X)\right). \tag{87}$$

For a set of edges $G\in\mathcal{P}(E)$, we wish to count the number of different paths which use only edges from within $G$, and connect from a vertex $v\in V$ to any vertex $a$ in set $X\subseteq V$. To do this, we introduce a function $c: V\times\mathcal{P}(E)\times\mathcal{P}(V)\to\mathbb{Z}_0^+$

$$c(v, G, X) = |\{g \in \mathcal{P}(G) : g \text{ is a path from } v \text{ to any } x \in X\}| \quad (88)$$

Suppose $X \subseteq Y$. The function $R(d^V(e), G, X, Y)$ gives the ratio of the number of unique paths from the vertex descending from the edge $e$ to any vertex in the set $Y$ that are also paths to a vertex in the set $X$.

$$R(d^V(e), G, X, Y) = \begin{cases} \dfrac{c(d^V(e), G, X)}{c(d^V(e), G, Y)} & \text{when } c(d^V(e), G, Y) \geq 1; \\ 0 & \text{when } c(d^V(e), G, Y) = 0. \end{cases} \quad (89)$$

Applying $\kappa$ to the function $R$ gives us partitioned-$\varphi(X, Y)$.

$$\text{partitioned-}\varphi(X, Y) = \kappa\left(R, (X, Y)\right) \quad (90)$$

We now turn to an example that shows that the calculation of partitioned-$\varphi$ may require an large number of terms, i.e., non-polynomial in the size of the terminal vertex set. Suppose that $T = (V, E)$ is the six terminal vertex caterpillar tree in Figure 8 and that we wish to calculate partitioned-$\varphi(\{x_2\}, X)$, where $X$ is the set of leaves of $T$. During the calculation of partitioned-$\varphi(\{x_2\}, X)$ a number of terms cancel, leaving a final sum of 'essential terms' required for the end result. For example, consider the sets of edges $\{c, d, g\}$ and $\{c, d, g, i\}$ in $E$. In $\kappa(R, (\{x_5\}, X))$ the following terms appear that correspond to these sets of edges:

$$\beta(c)\beta(d)\beta(g)(1-\beta(a))(1-\beta(b))(1-\beta(f))(1-\beta(h))(1-\beta(i))(1-\beta(j))(1-\beta(k)) \cdot \left[\sum_{e \in E} \alpha(e) \cdot R(d^V(e), \{c, d, g\}, (\{x_2\}, X))\right], \text{ and} \quad (91)$$

$$\beta(c)\beta(d)\beta(g)\beta(i)(1-\beta(a))(1-\beta(b))(1-\beta(f))(1-\beta(h))(1-\beta(j))(1-\beta(k)) \cdot \left[\sum_{e \in E} \alpha(e) \cdot R(d^V(e), \{c, d, g, i\}, (\{x_2\}, X))\right]. \quad (92)$$

The addition of edge $i$ to the set $\{c, d, g\}$ does not create any new paths that connect an interior vertex to $X$. Thus, the values of $R(d^V(e), \{c, d, g\}, (\{x_2\}, X))$ exactly match $R(d^V(e), \{c, d, g, i\}, (\{x_2\}, X))$ for all $e \in E$. Specifically, the sum in the square brackets in both terms (91) and (92) equals $\alpha(b)$. Noting this, we add the terms (91) and (92) together. The $\beta(i)$ and $(1 - \beta(i))$ parts cancel, leaving:

$$\beta(c)\beta(d)\beta(g)(1-\beta(a))(1-\beta(b))(1-\beta(f))(1-\beta(h))(1-\beta(j))(1-\beta(k)) \cdot \left[\sum_{e \in E} \alpha(e) \cdot R(d^V(e), \{c, d, g\}, (\{x_2\}, X))\right], \quad (93)$$

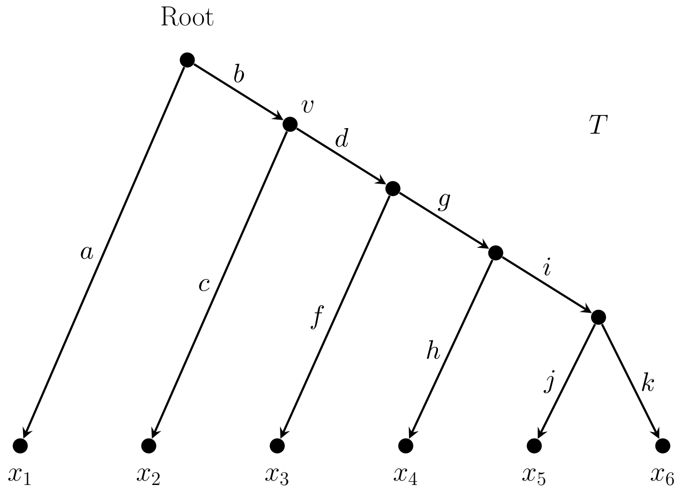

Figure 8: A caterpillar tree on six leaves. For the family of caterpillar trees, calculating partitioned-$\varphi$ may require exponentially many terms.

which does not contain any $i$ terms. We continue this process of selecting pairs of sets of edges in $E$ that differ in exactly one edge, and have identical connections to the terminal vertex set.

Yet, even after simplifying $\kappa(R, (\{x_2\}, X))$ as much as possible in this manner, a large number of terms remain. Each such term corresponds to the minimal subtree that connects each subset of $\{x_2, x_3, x_4, x_5, x_6\}$ to vertex $v$. Examples of these terms are given in Table 7. There is one term for each subset of $\{x_3, x_4, x_5, x_6\}$ (because $x_2$ must always appear); that is, $2^4 = 16$ separate terms to calculate. Partitioned-$\varphi(\{x_2\}, X)$ equals the sum of these terms. Each term must be included, even if we are using standard conditions and there is a uniform value of $\beta(e)$ for each edge $e$ in $E$. This is because sets of leaves of the same size may have different numbers of edges connecting these leaves to vertex $v$ and thus different numbers of factors in their corresponding terms.

We can extend this example to caterpillar trees with $n$ leaves and find that at least $2^{n-i}$ terms are required to calculate partitioned-$\varphi(\{x_i\}, X)$.

### 3 Supplementary information related to the application to living fossils

#### 3.1 living-fossil-ness measures based on ED

For comparison, we have included a figure equivalent to Fig. 9 in the main text only using ED instead of Partitioned-$\varphi$ (see Fig. 9)

| Set of leaves | Edge set | Associated term |
|---|---|---|
| $\{x_2\}$ | $\{c\}$ | $\alpha(b)$ |
| $\{x_2, x_4\}$ | $\{c, d, g, h\}$ | $-\frac{1}{2}\alpha(b)\beta(c)\beta(d)\beta(g)\beta(h)$ |
| $\{x_2, x_3, x_5\}$ | $\{c, d, g, i, j\}$ | $\frac{1}{3}\alpha(b)\beta(c)\beta(d)\beta(g)\beta(i)\beta(j)$ |
| $\{x_2, x_3, x_4, x_5\}$ | $\{c, d, f, g, h, i, j\}$ | $-\frac{1}{4}\alpha(b)\beta(c)\beta(d)\beta(f)\beta(g)\beta(h)\beta(i)\beta(j)$ |
| $\{x_2, x_3, x_4, x_5, x_6\}$ | $\{c, d, f, g, h, i, j, k\}$ | $\frac{1}{5}\alpha(b)\beta(c)\beta(d)\beta(f)\beta(g)\beta(h)\beta(i)\beta(j)\beta(k)$ |

Table 7: Example sets of leaves, the edges that connect these sets to $v$ and their associated terms.

##### 3.2 Comparison between ED and our living-fossil-ness measure based on paritioned-$\varphi$

Living-fossil-ness ranks of species based on ED are identical to those based on paritioned-$\varphi$ for the Quaternary period ('Quaternary' case) with $\rho = 0$. This is expected given the mathematical analysis of the $\rho = 0$ case in section 2.5. Living-fossil-ness ranks of species based on ED are closely correlated, but not identical, to the 'Quaternary' case with $\rho = 0.01$. The relationship of Living-fossil-ness ranks between the Quaternary and Cretaceous cases (both for $\rho = 0$ and for $\rho = 0.01$) is extremely weak overall, though the very highest ranking 'Cretaceous $\rho = 0$' species also tend to have high values under 'Quanternary $\rho = 0$'. The 'Jurassic $\rho = 0$' case yields a distribution with only three possible values (corresponding to monotremes, marsupials, and everything else). The 'Jurassic' case with ($\rho = 0.01$) predicts very different ranks to the 'Jurassic $\rho = 0$' case particularly in the sense that it creates much finer distinctions between species instead of three discrete categories. The 'Jurassic' case with $\rho = 0.01$ compares closely to the 'Cretaceous' case with $\rho = 0.01$ though with some clear deviations from a 1:1 relationship in the form of tight groups within which ranks may rearrange between the cases, but between which ranks will not change much (see Fig. 10).

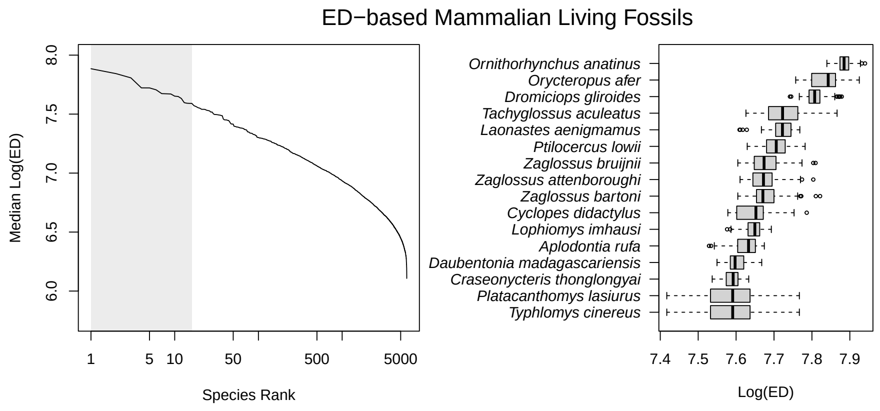

Figure 9: Living-fossil-ness for mammal species calculated using classic ED. The left panel shows the living-fossil-ness with rank in log scale. The box and whisker plot on the right shows the range of values for the top 16 species only. Each tree gives rise to one value in the distribution shown (see methods in the main text).

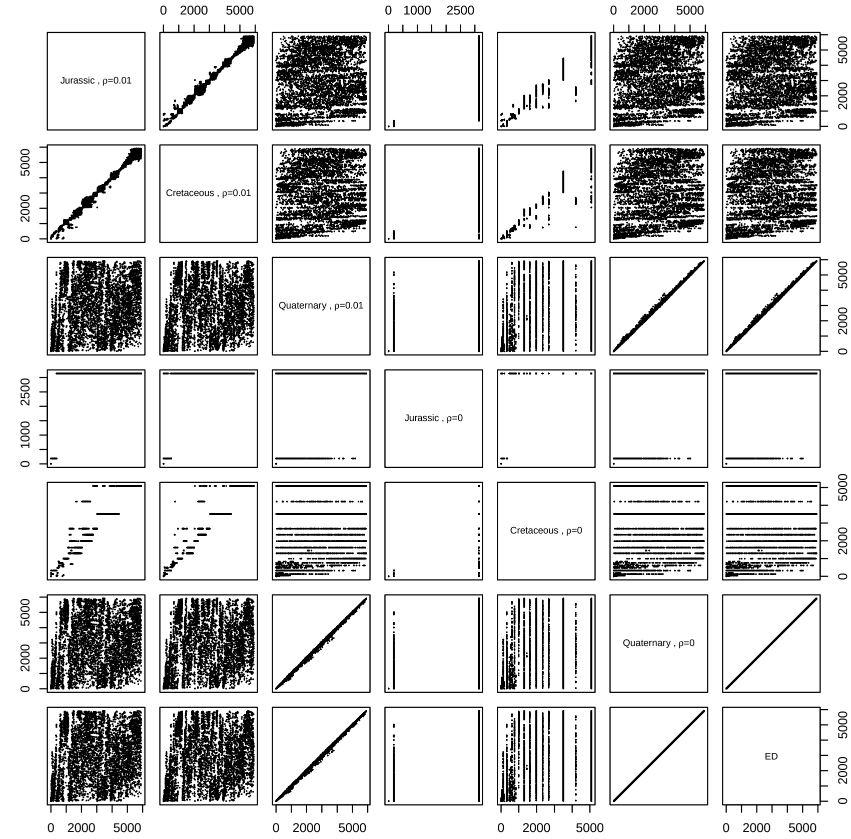

Figure 10: Paired scatter plot comparing the ranks of species according to different living-fossil-ness measures. We consider three periods of geological history: Jurassic (EvoHeritage from 145 million years ago, or earlier), Cretaceous (EvoHeritage from 66 million years ago, or earlier) and Quaternary (EvoHeritage from any period). Each of these is evaluated with $\rho = 0.01$ and $\rho = 0$. Quaternary living fossils with $\rho = 0$ correspond precisely to rankings under classic ED as expected from the results given in section 2.5. Cretaceous living fossils with $\rho = 0$ (labelled 'Cretaceous $\rho = 0$') correspond to ED measured on a tree with everything post-Cretaceous removed. Likewise for the 'Jurassic $\rho = 0$' results.
