## Supplementary Glossary for "Phylogenetic Biodiversity Metrics Should Account for Both Accumulation and Attrition of Evolutionary Heritage"

September 4, 2023

James Rosindell[1,2,*], Kerry Manson[2], Rikki Gumbs[3], Will Pearse[1] and Mike Steel[2]

[1] *Department of Life Sciences, Silwood Park Campus, Imperial College London, Buckhurst Road, Ascot, Berkshire, SL5 7PY, United Kingdom*
[2] *Biomathematics Research Centre, University of Canterbury, Christchurch, New Zealand*
[3] *EDGE of Existence Programme, Zoological Society of London, Regent's Park, London NW1 4RY, UK*

### Glossary of EvoHeritage Terms

**Tree**: In this work we consider only rooted trees which are a set of vertices and a set of directed edges, each edge connecting a pair of vertices. A (rooted) tree has the property that for any vertex, there exists a unique path to it, through the edges, starting at the root vertex.

**Root vertex**: A vertex of a rooted tree which has no edges directed towards it, equivalently this means it has no ancestral edges.

**Terminal vertex**: A vertex of a tree which has no edges directed away from it, equivalently that means it has no descendant edges. A terminal vertex is sometimes referred to as a leaf or a tip.

**Interior vertex**: A vertex that is neither a terminal vertex nor a root vertex.

**Terminal edge**: An edge of a tree directed towards a terminal vertex.

**Path**: Any set of edges that define a directed route between two given vertices of a tree.

$\hat{o}$: A root vertex corresponding to the origin of life.

**EvoHeritage**: short for Evolutionary Heritage this is a broad term for biological features in the most general sense including traits, functions, genetic sequences, aesthetic features, and future options for humanity, but explicitly only the components that are heritable and present in a given set of vertices. EvoHeritage may be measured as a continuous or discrete quantity.

**Accumulation**: A natural process by which novel EvoHeritage is generated along edges with copies being passed on to descendants.

**Attrition**: A natural process by which EvoHeritage is lost over time without extinction having occurred, for example, by not being passed on to any surviving descendants.

**EvoHeritage tree**: A rooted tree where the accumulation and attrition of EvoHeritage occurs in a specified way along each edge.

**EvoHeritage form**: EvoHeritage is considered to be of the same 'form' if it is subject to the same rules of accumulation and attrition on an EvoHeritage tree. An EvoHeritage tree may thus specify one or more forms of EvoHeritage, each with its own rules of accumulation and attrition. The majority of this work is concerned with a single form of EvoHeritage, to have multiple forms simply requires considering each independently.

**Species**: The species concept is optionally incorporated by identifying species as either a single vertex or as a set of vertices in an EvoHeritage tree.

$\boldsymbol{\varphi}$: For a given set of vertices on an EvoHeritage tree, the total quantity of EvoHeritage present on those vertices.

**Evolutionary History**: For a given set of vertices on an EvoHeritage tree, the total quantity of EvoHeritage that has ever been present on those vertices or any of their ancestors. This is equivalent to the PD of the given set of vertices, including the root, which we take to be $\hat{o}$ by default.

**Unique-$\boldsymbol{\varphi}$**: For two given sets of vertices on an EvoHeritage tree, this gives the amount of EvoHeritage that is present within the first set of vertices and absent from the second set of vertices.

**Diff-$\boldsymbol{\varphi}$**: For two sets of vertices on an EvoHeritage tree, diff-$\varphi$ is the proportion of total EvoHeritage that is only present within one of the two sets.

**Partitioned-$\boldsymbol{\varphi}$**: A partitioning of EvoHeritage between vertices on an EvoHeritage tree, where the value of any part of EvoHeritage is divided equally among all copies of it.

**Dated EvoHeritage tree**: An EvoHeritage tree where all vertices also have dates attached.

**Extant vertex**: A terminal vertex with date = 0 on a dated EvoHeritage tree.

**Extinct vertex**: Any vertex with date > 0 on a dated EvoHeritage tree.

**Targeted EvoHeritage tree**: Constructed from a dated EvoHeritage tree by artificially augmenting EvoHeritage accumulation on certain edges in order to study the fate of targeted ancestral EvoHeritage. For example, removing EvoHeritage accumulation except during a key period in geological history.

**Living fossil**: An extant vertex with a large share of rare EvoHeritage from a period of geological history. After the total EvoHeritage present at a selected period of geological history has been partitioned among the extant vertices of a dated ancestral EvoHeritage tree (defined at species level), those extant vertices (species) with a high value are living fossils.

**Standard conditions**: For a tree with edge lengths, standard conditions generate an EvoHeritage tree according to a simplified set of rules. Specifically, (i) all EvoHeritage is of the same form and measured as a continuous quantity, (ii) EvoHeritage accumulates along edges at a fixed deterministic rate of $\lambda$ units of EvoHeritage per unit of edge length and (iii) attrition occurs at a fixed deterministic rate of $\rho$ per unit of EvoHeritage per unit of edge length. The rate of attrition is thus proportionate to the total amount of EvoHeritage.

**Standard stochastic conditions**: For a tree with edge lengths, standard stochastic conditions generate an EvoHeritage tree according to a simplified set of rules. (i) all EvoHeritage is of the same form and measured as a discrete quantity, (ii) EvoHeritage accumulates along edges at a fixed stochastic rate of $\lambda$ units of EvoHeritage per unit of edge length and (iii) attrition occurs at a fixed stochastic rate of $\rho$ per unit of EvoHeritage per unit of edge length. These rules are akin to Standard conditions with the only difference being that EvoHeritage is measured in discrete units and the rates of Attrition and Accumulation describe stochastic events rather than a gradual deterministic flow.

**Standardised units**: Under standard conditions (or standard stochastic conditions), the total EvoHeritage at the end of a single disconnected edge with a unit length. Alternative standard lengths could be used e.g. the time back to $\hat{o}$, with $\hat{o}$ included as a vertex in the calculations. Alternatively, the median (or some other property) of the distribution of terminal edge lengths of species could be chosen in order to make standardised units relatable to the familiar species concept. The size of a standardised unit is dependent on both Attrition ($\rho$) and Accumulation ($\lambda$) as well as the standard edge length that it is based on.

$\boldsymbol{\varphi_\rho}$: A convenient notation for $\varphi$ under standard conditions (attrition at rate $\rho$) and measured in standardised units on an EvoHeritage tree. Measuring EvoHeritage in standardised units removes any dependency on $\lambda$ leaving only a dependency on $\rho$. $\varphi_\rho$ can thus be used in the same way as PD for any tree and a given $\rho \geq 0$. We may also write $\varphi_\rho$ in place of $\varphi$ for the other metrics involving $\varphi$. For example unique-$\varphi_\rho$.
